# Compositional memory matters for early molecular systems

**DOI:** 10.64898/2026.03.03.709225

**Authors:** Ledoux Barnabé, Ryoka Kuwabara, Norikazu Ichihashi, Ryo Mizuuchi, Lacoste David

## Abstract

The error catastrophe refers to the proliferation of non-functional molecules in conditions where molecular replication has low accuracy, which is likely to correspond to conditions present at the Origin of Life. This error catastrophe can be avoided thanks to transient compartmentalization, provided that the compartments are sufficiently tight to prevent molecular leakage. Typically, transient compartmentalization models assume that the content of the compartments is completely pooled periodically, resulting in the complete loss of the compositional memory of the compartment. Furthermore, previous models that include the possibility of ecological interactions between molecular parasites and replicators within compartments generally do not study the effect of transient compartmentalization on their coevolution. To address both issues, we develop a framework that accounts for the coevolution of molecular replicators and parasites, along with specific compartmentalization dynamics that are transient yet partial, allowing compositional memory to accumulate from one round of compartmentalization to the next. We benchmark our model with a serial dilution experiment that displays complex oscillatory dynamics among four well-characterized RNA replicators. We also perform experiments to quantify the level of mixing in compartments when stronger stirring tends to homogenize their composition. We then model stirring-induced mixing and show how stirring alters the dynamics of compartmentalized replicators. We conclude that compositional memory arising from transient compartmentalization plays a major role in the dynamics of early molecular systems.

---

While chemical systems are often studied in well-mixed conditions with only a few species, it seems more likely that life emerged in a heterogeneous medium with complex composition [1, 2, 3]. Such a medium would then contain regions in which diffusion is limited, which we call compartments. Compartmentalization, even when only partial and transient, could solve essential issues for the Origin of Life [4, 5, 6, 3, 7, 8, 9, 10, 11, 12]: It can keep important molecules nearby in a small volume, which could enhance reactivity [13, 14], it can limit side-reactions and parasite amplification, thus preventing the error catastrophe first identified by Eigen [15, 16, 17, 18], and it could lead to new forms of autocatalysis and ecological interactions, which simply do not exist in well-mixed systems [19]. Moreover, compartmentalization could facilitate compositional inheritance [20, 21, 22,, which is a primitive form of heredity that does not require information-carrying polymers nor enzymes.

Recently, a scenario for the emergence of an RNA world in three stages has been proposed [24], in which the last two stages involve a form of compartmentalization. In stage 1, network autocatalysis occurs using oligomer substrates to form catalysts [25]. In stage 2, autocatalysis coexists with templated ligation, and limited diffusion acts as a primitive form of compartmentalization [7, 4], thereby preventing the error catastrophe. Then, in stage 3, template-based replication takes over thanks to enhanced catalytic properties, smaller building blocks are used, and more efficient forms of compartmentalization appear via coacervates and lipid vesicles. The transition from stage 2 to stage 3, leading to the dominance of template-based replication, has been studied before by several authors [26, 27]; the transition is stochastic and becomes less likely in well-mixed systems, suggesting that concentration fluctuations also matter for life to appear [28, 29].

This scenario supports the view that transient compartmentalization is crucial for the maintenance and the transfer of molecular information [4, 30, 31, 32, 33, 34]. In practice, transient compartmentalization dynamics involve a periodic contact with the external environment and a certain level of spatial inhomogeneity. Some experiments confirmed the theoretical prediction that transient compartmentalization can limit parasite amplification [30, 35, 31, 34, 32, 33, 36, 37]. When performed over long timescales, experiments on compartmentalized RNA-based systems showed that RNA replicases coevolved with their parasites and formed increasingly complex molecular ecosystems [34, 38].

One question raised by these experiments [34, 38] is the role of compartmentalization during replicase-parasite coevolution [33, 31, 32]. A related issue in these experiments was the lack of quantification of the composition of individual compartments and of the degree of mixing between compartments.

To address these challenges, we first revisit and extend previous models of transient compartmentalization, which assume that compartment contents are fully mixed during periodic pooling [31, 37]. Our extension explicitly accounts for the coevolution of multiple replicases and parasites within compartments. We then introduce a model where compartments partially inherits their composition from one round to the next, linking this compositional memory to the average degree of stirring applied to the compartment population. Stirring induces fusion and mixing among compartments, thereby modulating the inheritance of molecular compositions.

Our model successfully reproduces the four-species oscillatory dynamics observed in a serial dilution experiment using previously characterized RNA replicators, which replicate via protein translation [34]. These replicators include self-replicating RNA molecules (replicases) and parasitic sequences that exploit replicases for their own replication. Although the experimental system is based on protein translation, the theoretical model is more general and describes other replicator-parasite systems such as ribozymes and autocatalytic networks. We demonstrate that when parasites dominate the system—reducing the average number of molecules per compartment to near unity—sufficient mixing is essential to spatially isolate parasites from replicases and purify the system. Without adequate mixing, parasites persist even at low densities, leading to complete washout through successive dilutions. This behavior echoes the role of dispersal in other replicase-parasite models [39]: parasites cannot proliferate if they fail to encounter replicases (e.g., due to limited diffusion).

We performed competition experiments using compartmentalized RNA-replicases and parasites, and we confirmed our theoretical expectation that stirring generally improves the survival of replicases. Weak stirring, conversely, preserves compositional memory, promoting parasite fixation. Using fluorescent dyes, we quantified the degree of mixing between compartments as a function of stirring strength, analyzing fluorescence distributions across droplets. Collectively, our findings reveal that compartmentalization alone is insufficient to prevent parasite invasion. Instead, controlled mixing is critical for enabling early molecular replicators to evolve into robust, self-sustaining systems.

## Model and observables

We model the serial transfer experiment with 4 species introduced in Figs. 1A-B, where we specify the mechanisms of replication of RNA-replicases and RNA-parasites. Transient compartmentalization dynamics of RNA molecules has been theoretically modeled in previous work [31, 32, 33, 34, 37] as a cycle consisting of four steps shown in Fig. 1C : (i) *compartmentalization* : from an initial pool of replicases and parasites, compartments are formed by seeding them with molecules according to a Poisson distribution [40]; (ii) *maturation* : each compartment evolves depending on its initial composition during a maturation time *T*_*mat*_ set by the environment. This time is assumed to be long enough for each compartment to reach its carrying capacity; (iii) *pooling* : after maturation, compartments fuse and release their contents in a common pool, and the fractions of each subpopulations are updated; (iv) *dilution* : the content of the pool is diluted homogeneously with a factor *d*, without affecting the fractions of each subpopulations. The process can start again.

**Figure 1.**
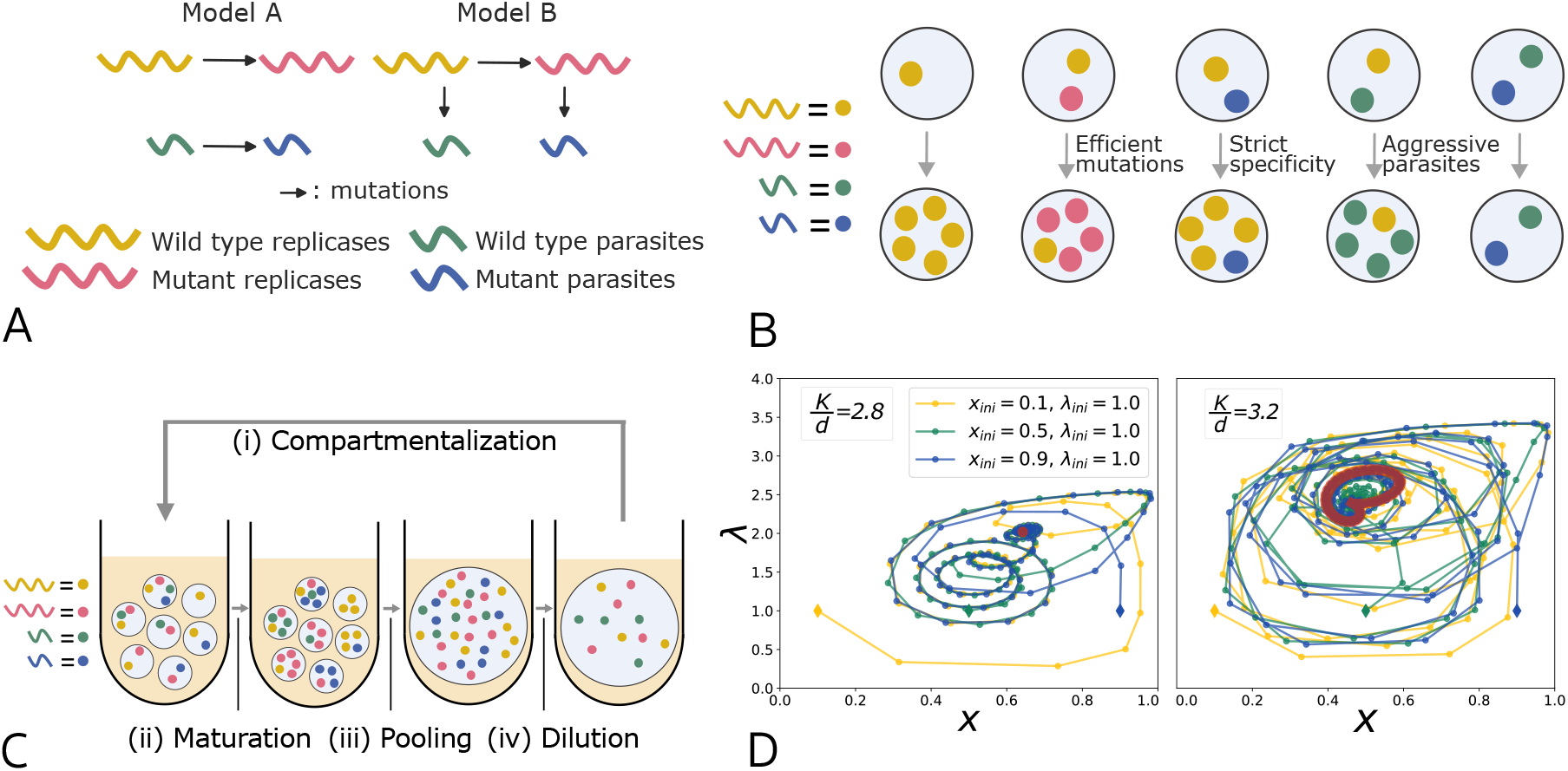
**A** Representation of two possible evolution models of the RNA replication system containing two replicases and two parasites. **B** Illustration of the deterministic assumptions. **C** Transient compartmentalization with pooling (homogeneous case). **D** Illustration of the typical oscillatory dynamics obtained with the deterministic approach with second order replication rates and mutations. Here we show the results for model B, we used the rates *α*_0,0_ = 0.5*h*^1^, *α*_1,1_ = 2.0*h*^1^, *γ*_0,0_ = 7.0*h*^1^, *γ*_1,1_ = 20.0*h*^1^, *µ*_0_ = 1.0 ×10^−2^*h*^1^, *ν*_0_ = 1.0 ×10^−2^*h*^1^ and *ν*_1_ = 1.0 ×10^−2^*h*^1^. The carrying capacities are *K* = *K*_*r*_ = *K*_*p*_ = 28 on the left panel and *K* = *K*_*r*_ = *K*_*p*_ = 32 on the right panel, and the dilution factor is *d* = 10. Limit cycles are shown in red (which is a single point for a steady state). We observe damped oscillations reaching a steady state on the left panel, and sustained oscillations (limit cycles) on the right panel.

### Maturation

In the maturation step, we consider two different models, defined in Fig. 1B depending on how parasites are created. Both models are similar to that of [31] except that they account for the possible coevolution of replicases and parasites through mutations. Model A considers that mutant parasites (resp. replicases) evolve only from wild type parasites (resp. replicases) due to mutations. The dynamical equations [41] for the populations of the various species are given in Eq. 5.

In model B, parasites can evolve from replicases. In practice, this can happen due to deletions in the sequence of replicases (by recombination for instance [34]), which yield smaller sequences unable to replicate by themselves. The dynamical equation for the populations of the various species in this model is given by Eq. 14. Both models are based on experimental observations obtained by tracking evolutionary lineages of replicase and parasite RNAs [34].

### Pooling

After the maturation step, the contents of all compartments are pooled (they fuse and all their contents are mixed in a common pool). Therefore, compartments do not inherit from their content before pooling (compositional memory is erased). The new fraction of replicases *k* in the pool is expressed in terms of the value at the beginning of the round by the following equation:

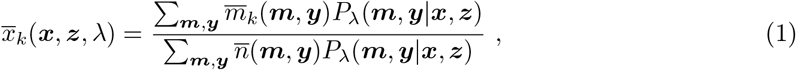

where 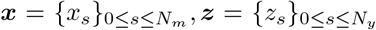 are vectors describing the fractions of each mutant replicases (where *N*_*m*_ is the total number of mutant replicases) and parasites (where *N*_*y*_ is the number of mutant parasites), respectively. Further, *λ* is the parameter of the Poisson distribution, and *P*_*λ*_(***m, y x, z***) is the multinomial probability to have a vector ***m*** for the numbers of each replicase and a vector ***y*** for the numbers of each parasite starting with a pool with fractions ***x*** of replicases and fractions ***z*** of parasites. Then 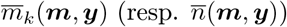 is the final number of replicases after maturation of type *k* (resp. final number of molecules after maturation) in the compartment given that it was initially filled with *n* molecules among which ***m*** were replicases and ***y*** parasites. In principle, 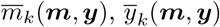 should be random variables but they become fully determined by ***m*** and ***y*** in our *deterministic approach* (see below). We can set the time of maturation to be *T*, or fix the final number in each compartment as a constant value 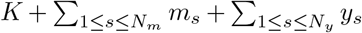. In the following, we assume that *T* is large enough for both conditions to be equivalent.

## Results

### Deterministic approach

There are two sources of stochasticity in the dynamics we consider: the first arises from inoculation [40], i.e. the preparation of the initial composition of compartments by sampling molecules from the pool. The second arises during the maturation phase, in which each compartment evolves stochastically through mutation or replication events.

In the deterministic approach, it is assumed that the maturation of a compartment is fully determined by its initial composition, which means that the only remaining source of stochasticity in the system is the inoculation step. This has the advantage of providing simple dynamics that are independent of the kinetic parameters of the maturation step, such as the replication rates. The only parameters controlling the evolution of the system are then the carrying capacity (assuming resources are homogeneously distributed among compartments), the dilution factor, the maturation time and mutation rates for mutations. The carrying capacity arises from the limitation of available resources in one compartment and constrains the total number of molecules maximally produced in that compartment.

This approach relies on specific assumptions about species interactions. In SI appendix Sections 5-8, we explore how varying these assumptions—such as the potential for parasites to evolve and acquire generality (i.e., the ability to replicate with all replicases)—affects the system dynamics. For Fig. 1D, Fig. 2B, and Fig. 3, we adopt the following key assumptions:

**Figure 2.**
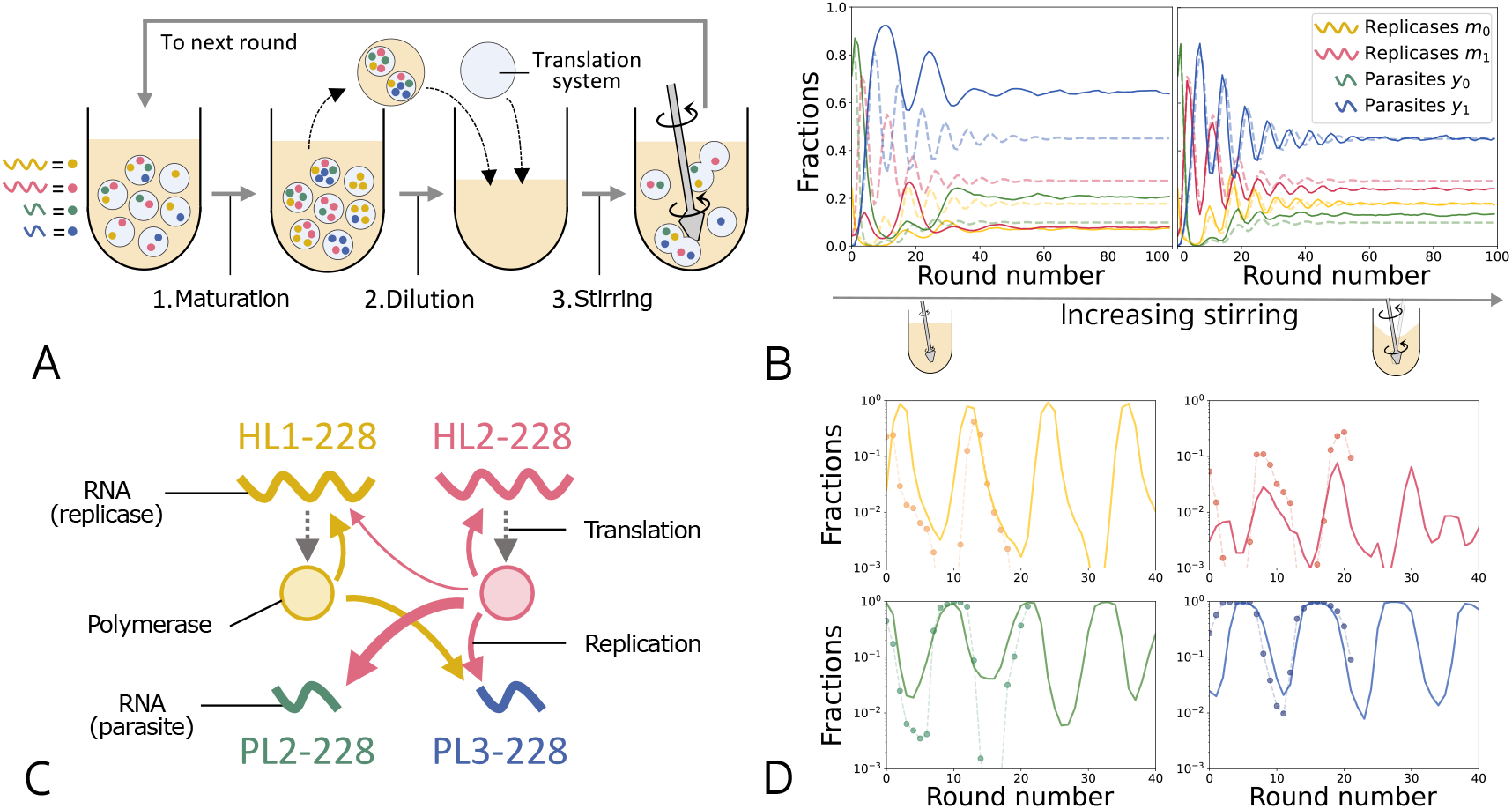
**A** Schematic representation of the long-term replication experiment. 1) RNAs were replicated in water-in-oil droplets at 37°*C* for 5 h. 2) Droplets were five-fold diluted with a fresh oil phase and the translation system. 3) The droplet population was stirred, allowing fusion and division, thereby redistributing RNAs among droplets. **B** Effect of stirring on the dynamics, we used the same deterministic rules and parameters but for different values of the stirring parameter *s* (s=0.5 and s=0.99). The solid lines represent the simulations with stirring while the dotted lines represent the perfectly homogeneous case. Simulations with stirring were performed with 50000 compartments, the carrying capacities are *K*_*p*_ = *K*_*r*_ = 30 and dilution factor *d* = 10. Oscillations are damped in this panel because *K/d* is small enough, (as illustrated in Fig. 1D and detailed in SI appendix Section 1.F). **C** The RNA replication system containing two replicases (HL1-228 in yellow and HL2-228 in red) and two parasites (PL2-228 in green and PL3-228 in blue). These RNAs replicate by utilizing polymerases translated from the replicases. The widths of the colored arrows represent relative replication efficiencies [37]. **D** We can reproduce experimental dynamics of four RNA species (2 replicases and 2 parasites), represented by solid circles connected by thin dashed lines. The solid lines show simulations performed using the same dilution factor *d* = 5 and the experimentally determined replication rates of the individual species [34]. Each species has a different carrying capacity, determined by its length. Other parameters are detailed in SI appendix Section 2.D.

**Figure 3.**
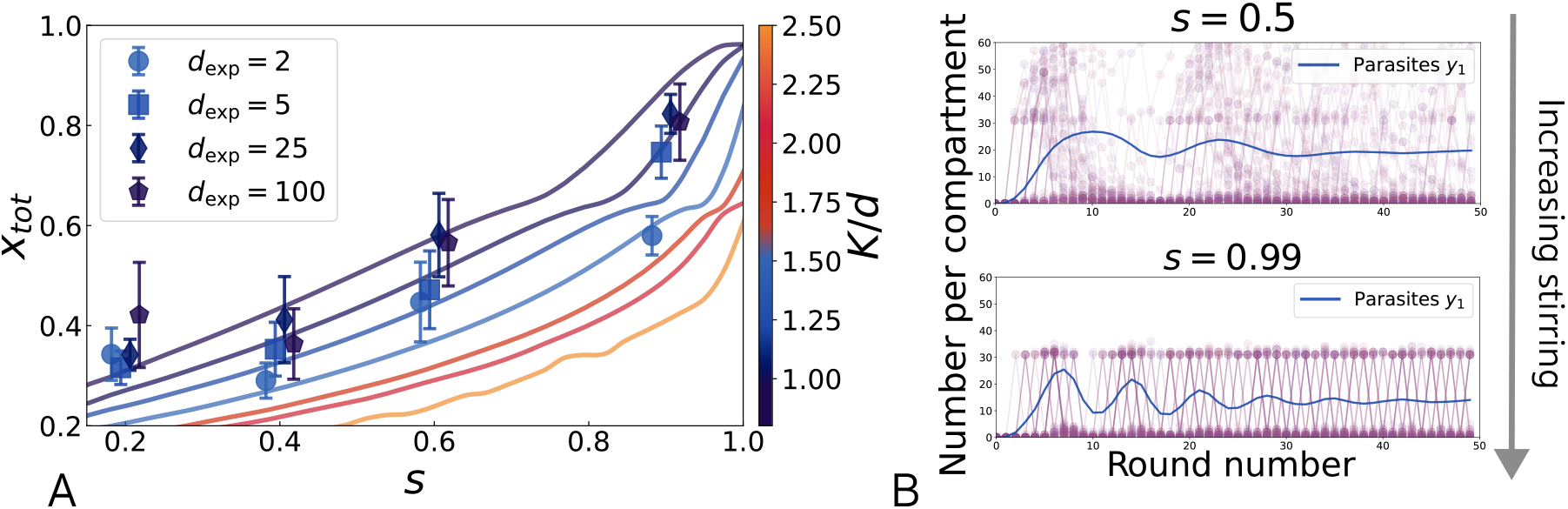
**A** Total fraction of replicases in the system, *x*_*tot*_. Theoretical expectations (solid lines) are shown for different values of the ratio of carrying capacity to dilution factor *K/d*. Experimental data (solid symbols) are plotted against stirring strength (2.0, 5.2, 16, and 30 krpm), converted into the corresponding stirring parameters *s*. For visualization, the data points are plotted at the expected *s* ±0.01 0.02. The experiments were performed at different dilution factors, *d*_exp_, as indicated in the panel. The same trends are observed in theory and experiments. Error bars indicate the mean ±one standard deviation (n = 3–4). **B** Effect of stirring on the compartment composition. In purple, we show the trajectories of individual compartments (for the number of mutant parasites *y*_1_), while in solid blue lines we show the average over all compartments. As stirring gets stronger, the contents of compartments homogenize, and the compartments end up either full of mutant parasites or containing mostly no mutant parasites. In this limit, all compositional memory is lost.

#### Aggressive parasites

In compartments containing both parasites and their associated replicases (i.e., replicases capable of replicating these parasites), parasites dominate and replicate until they reach their carrying capacity, which may differ for replicases and parasites depending on sequence length [31].

#### Strict specificity

Parasites can replicate only with replicases of the same type, reflecting experimental observations of parasite-replicase ecosystems [38]. Consequently, in compartments containing parasites of type *k* and replicases of type *j* ≠ *k*, only replicases replicatMutated replicases or parasites exhibit significantly higher replication rates than their wild-type counterparts, with no back-mutations. For example, in compartments containing only replicases, those with the highest fitness (i.e., replication rate) dominate. Under these assumptions, each initial condition maps to a unique final composition.

Further, we assume that the maturation time is long enough for every compartment to reach its carrying capacity providing it contains replicases initially. The lengths of dominant parasite RNAs that emerged from replicases in previous evolution experiments were approximately 4 times smaller than that of replicases [34]. If one assumes that the carrying capacity originates from a limit in terms of the total number of nucleotides, then this factor four represents the ratio between the carrying capacity of the parasites and of the replicases as explained in SI appendix Section 11. This effect is not taken into account in most simulations except for the comparison to experimental data in Fig. 2D. This effect mostly increases the amplitude of replicase-parasite oscillations and prevents them from reaching steady states. In this case, the average fraction of replicases tends to be lower, and the isolation of parasites in distinct compartments becomes less likely. Yet the effects of dilution and stirring are similar to the case where all species have the same carrying capacity (see SI appendix Section 11), and the general conclusions we draw are not modified.

Using this method, we obtain oscillatory dynamics between the different species, as can be seen in Fig. 1D. We see in Fig. 1D that oscillations manifest as cycles on the plane (*x, λ*). When both *λ* and *x* are small, parasites become isolated, leading to an increase in the replicase fraction *x*. This subsequently raises the average number of individuals per compartment *λ*. At higher *λ*, parasites will more likely find replicases and start invading the system, thereby decreasing *x*. The resulting reduction in replicases leads to fewer replication events and thus a decrease in *λ*, and the cycle is ready to start again. The mechanism underlying these oscillations is detailed in SI appendix Section 1.F. In contrast with the theoretical study of two-species dynamics [31], we explore the effects of mutations and four-species interactions.

### Stochastic vs. deterministic approaches

The deterministic approach ignores the stochasticity of mutations, but fortunately, their effect can be incorporated by computing the average numbers of mutants of each species appearing during one cycle (averaged over the initial compositions of the compartment), as explained in SI appendix Section 1.C. In that way, the deterministic method can account for different values of the replication and mutation rates. To assess if this improved deterministic approach fully captures the dynamics that arise from stochastic chemical reactions, we compare it to Gillespie simulations (Direct Stochastic Simulation Algorithm) [41] in Figs. S4 and S5. These simulations include the full stochasticity of all reactions and therefore account for all sources of stochasticity in the system. We find that the deterministic approach provides an accurate description in the regime where mutations are not too rare. Whenever mutation rates are small enough that mutations do not necessarily occur within one cycle, the dynamics predicted by the deterministic approach differ from those obtained with Gillespie simulations, yet it still predicts the correct average fractions of each species over cycles. In this limit, mutations become rare events that drastically modify the system. Depending on the conditions (typically the ratio *K/d*), we observe oscillations (either stable or unstable) or a stable node [31]. In particular, some conditions lead to limit cycles, and the complexity of orbits increases with four species as compared to two species (see SI appendix Section 1.F). These complex dynamics stabilize the four species system (see SI appendix Section 1.F), because the threshold between stable and unstable spirals is shifted towards larger *K/d* with four species.

This formalism can be extended to include more complex interactions between replicases and parasites, as observed experimentally [34]. In one extension, we have distinguished generalist-parasites, which are able to replicate with all types of replicases, and specialist-parasites [42], which are able to replicate with only one type of replicases but more efficiently as explained in SI appendix Sections 5A and B. We have also studied the role of various parameters such as the carrying capacities of replicases and parasites, mutation rates and maturation time in SI appendix Sections 9-12.

### Stirring and compositional memory

Despite these modeling efforts, experiments with the compartmentalized RNA systems still do not exactly match the assumptions of the models outlined above, because in practice, the solution is stirred periodically. This stirring causes some material exchange between compartments (sketched in Fig. 2A) due to fusion/fission events [34, 38] or other transfer mechanisms [43, 44], which are not described explicitly in the model. As a result, the composition of a given compartment before stirring is correlated with its composition after stirring in contrast to the assumptions made in the pooling step. We refer to this phenomenon as compositional memory, a feature which is central in some origin of life scenarios [21, 20, 45]. This compositional memory is expected to be relevant irrespective of whether compartments are made of droplets, coacervates or vesicles [46]. The effect of mixing from one round of compartmentalization to the next has previously been assessed numerically using simulations [37], but that work did not model the complex ecological interactions between molecular species precisely.

To address the issue of stirring, we now consider the experimental protocol described in Fig. 2A. After an initial *compartmentalization* step, each subsequent cycle consists of a *maturation* step as before, followed by a *dilution* step at the compartment level, in which randomly chosen mature compartments are removed and replaced by empty ones to increase the total available volume. We also assume that new droplets are enriched (for instance by enclosing resources present in the environment). Then, the *pooling* step is replaced by a *stirring* step, in which the solution is stirred so as to induce fusions and divisions of compartments. If this stirring was strong enough to completely erase the influence of the former content of compartments on their new composition, we would recover equivalent dynamics as with *pooling*, as illustrated in Fig. 2B.

The stronger the stirring, the weaker the compositional memory should be. In the limit of strong stirring, we expect to recover the result of complete pooling, meaning that the system has no memory. To include compositional memory, we consider that the composition of compartments is updated to a new composition determined by an interpolation between their former compositions and the average composition of the whole population. This is a mean field assumption because every compartment is then interacting with the average composition of the total population. This approach does not allow us to study spatially localized effects or the fusion/fission dynamics of individual compartments, but it is sufficient to capture the average effect of stirring on droplet content. In practice, we find that the new composition of a compartment depends on its former composition because stirring is often not strong enough to fully homogenize the system. We quantify the average level of stirring-induced mixing by a parameter *s* ∈ [0, 1] (*s* = 0 means that compartments do not interact with each others at all resulting in the absence of population scale dynamics, while *s* = 1 is the complete pooling limit), and we denote with a hat the quantities computed after the stirring step. For instance, 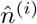 (the total number of individuals in compartment *i* after dilution) is drawn according to a Poisson distribution of parameter:

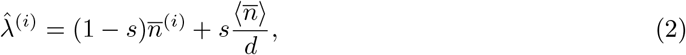

where 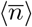 is the average number of individuals after maturation and before dilution, and 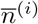 is the number of individuals in compartment *i* after maturation if it has not been removed during dilution, and 0 otherwise. We deduce from this expression that *s* = *d/*(*d* + 1) corresponds to the level of stirring where the old composition and the average composition after dilution contribute equally to the definition of 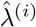. Thus, when *s > d/*(*d* + 1), we expect to recover the perfectly homogeneous case. For instance, when *d* = 10, the corresponding value is *s* ≃ 0.91. From Eq. 2, we also deduce that the standard deviation of 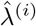 scales as 1 − *s* and we recover that the compartments are perfectly homogeneous for *s* = 1.

Then, the new numbers 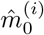 (and similar for other species) are drawn according to a multinomial distribution with parameters

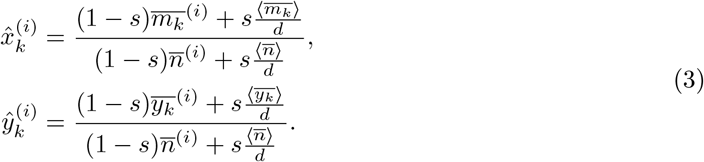

We keep the notations 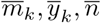 for the numbers of each species in one compartment after maturation and before dilution, 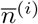 is the number of individuals in compartment *i*. If *s* = 1, we recover the dynamics discussed in the previous sections (Fig. 2B) and the distribution is homogeneous (no memory). Whenever *s <* 1, we keep track of inhomogeneities (memory effect).

Note that this procedure implies following the trajectory of every compartment individually; this assumes a finite (fixed and large) number of compartments, in contrast to the previous assumption of an infinite number of compartments. For *s* →1 (strong stirring), the system loses its compositional memory, and we recover the results for the homogeneous case (perfect pooling). On the other hand, if *s* →0 (weak stirring), all the compartments are conserved over time and the system does not evolve (except for mutations).

To examine whether our theoretical model could reproduce experimental observations, we performed a serial dilution experiment using a previously constructed compartmentalized RNA replication system [34]. This system comprises single-stranded RNA molecules functioning as replicases and parasites, together with a reconstituted *Escherichia coli* translation system, all encapsulated in water-in-oil droplets. Each replicase encodes the catalytic subunit of an RNA-dependent RNA polymerase, which becomes active within the translation system. Both replicases and parasites compete for polymerases translated from replicases and for nucleotide triphosphates (NTPs). We initiated the serial-dilution experiment with two replicases (HL1-228 and HL2-228, 2041 nt and 2042 nt, respectively) and two parasites (PL2-228 and PL3-228, both 514 nt), all previously characterized [34]. Their replication network is detailed in Fig. 2C. Each round consisted of (1) RNA replication at 37°*C* for 5 h, (2) five-fold dilution of droplets with freshly prepared droplets, and (3) stirring (16 krpm, 1 min) to induce droplet fusion and division (Fig. 2A). Step (3) not only replenished the translation system in individual droplets but also redistributed RNA molecules across the new droplet population, thereby representing the partial pooling of replicases and parasites. We found that the four RNA species exhibited sustained replication with oscillatory dynamics for at least 22 rounds. The dynamics were consistent with those observed previously using the same four RNA species together with another weakly replicating RNA [34]. We then compared these dynamics with those generated by our theoretical model, based on the four RNA species with their experimentally determined replication rates [34] (Fig. 2D). We confirmed that the model (*s* = 0.79) largely reproduced the experimental dynamics. Details of the calculations are given in SI appendix Section 2.D.

### Stirring and replicases fraction

Let us now study how stirring affects the degree by which molecular information is maintained. This can be assessed by following the total fraction of replicases *x*_*tot*_ = *x*_0_ + *x*_1_ as a function of *s* for a given set of parameters. As stirring becomes weaker, the increased heterogeneity of compartment composition decreases the probability for parasites to be isolated, resulting in a lower fraction of replicases. We find that the dependence of *x*_*tot*_ as a function of *s* is governed by the ratio of the carrying capacity to the dilution factor *K/d*, and that it displays different curvatures in the regimes *K/d* ≫1 and *K/d <* 1 (Fig. 3A).

To test whether these results could capture the experimental behavior, we subjected a replicase (HL2-228) and a parasite (PL2-228) in the RNA replication system (Fig. 2C) to a competition assay. The assay was performed as a single-round transfer experiment (Fig. 2A) with varying stirring strengths and dilution factors *d*_exp_. First, we prepared droplets containing HL2-228 and PL2-228. Next, the droplets were diluted at different ratios (2to 100-fold) with new droplets containing neither replicases nor parasites. We then induced droplet fusion and division by stirring at various speeds (2–30 krpm) for 1 min using a homogenizer. The different stirring strengths barely affected the droplet size distributions (Fig. S14), and the average number of HL2-228 or PL2-228 molecules per droplet was likely less than one under all tested conditions. We did not vary other parameters that determine the carrying capacity, such as NTP concentrations. After incubating the droplets at 37°*C* for 5 h to allow RNA replication, we measured the concentrations of replicases and parasites, and calculated *x*_*tot*_. We found that *x*_*tot*_ increased with both stirring strength and dilution factor (Fig. 3A), consistent with the theoretical prediction. The correspondence between experimental stirring strengths in *krpm* and stirring parameters *s* was assessed from experiments detailed in the following section. These results further validate the theoretical model and the importance of accounting for stirring in the model, as it matters for the coevolution of replicases and parasites.

To further investigate how varying stirring strengths influence replicase-parasite competition, we used our model to calculate the number of mutant parasites per compartment as a function of *s* (Fig. 3B), which is difficult to assess experimentally. Interestingly, we find that the number of mutant parasites remains heterogeneous under moderate stirring, whereas strong stirring homogenizes compartments, which end up either full or almost empty of mutant parasites. In this limit, all compositional memory is lost. These results can explain how the fraction of replicases increases under stronger stirring, as parasites are more frequently isolated.

#### Quantification of the level of stirring in experiments

In the experiments, the degree of stirring mainly depends on the applied stirring strength (in rotations per minutes or rpm). To quantify the extent of content mixing between droplets as a function of stirring strengths, we prepared two droplet populations containing either green or red fluorescent dyes—fluorescein isothiocyanate (FITC)-labeled dextran or tetramethylrhodamine isothiocyanate (TRITC)-labeled dextran)—hereafter referred to as G and R, respectively. The two droplet populations were mixed, stirred at varying strengths, and subjected to a two-round transfer experiment (Fig. 4A). The degree of content mixing was assessed from measurements of the fluorescence intensities of G and R in individual droplets after rounds 1 and 2 (Fig. S13). To model this experiment, we rescale the fluorescence intensities by the average fluorescence per compartment for each color, and we call the corresponding distributions of green (resp. red) molecules 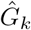 (resp. 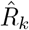). In addition, we define 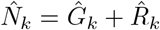 .

**Figure 4.**
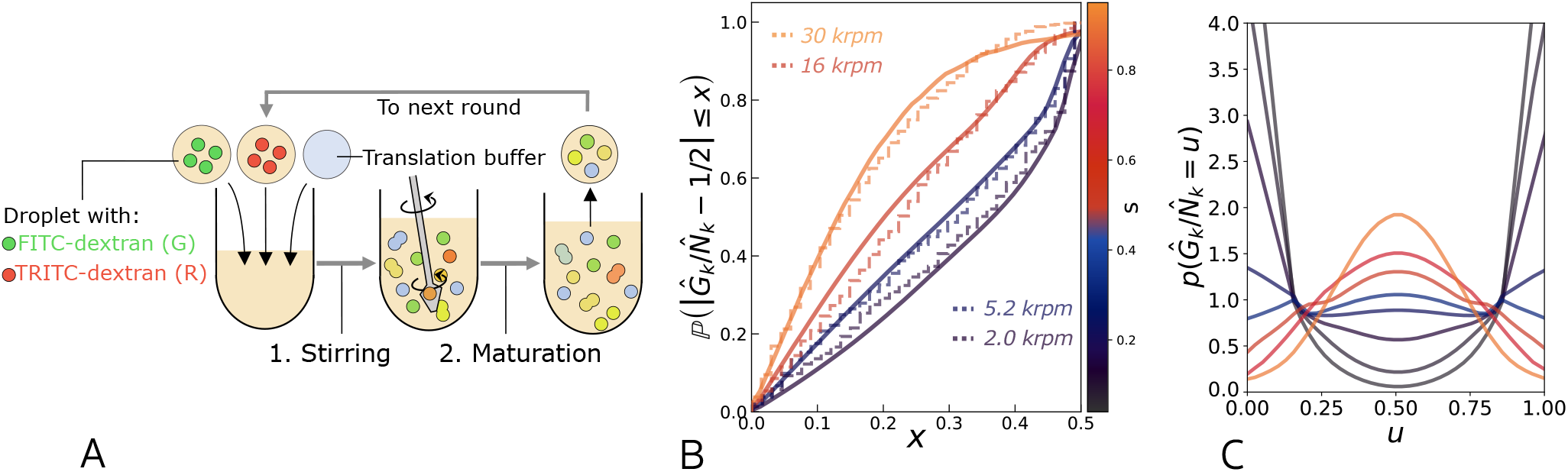
Influence of the stirring parameter *s* on the distribution of the fraction of one type of individuals, in a population with only two types of individuals. **A** Experimental procedure to assess the effect of stirring with two fluorescent dyes. After preparing the system from droplets containing either FITC-or TRITC-labeled dextran (*G* or *R*) together with the translation buffer, as described in the Materials Methods, the system was stirred for 1 min, followed by a 5 *h* incubation (maturation) phase. **B** Evolution of the experimental distributions (dotted lines, after round 1) for different stirring strengths and comparison with theoretical distributions (solid lines) starting from a perfectly bimodal distribution of 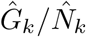 for different values of the stirring parameter *s*. We observe that the distribution of 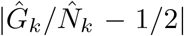 shifts towards 0 as stirring strengths increases, meaning that the distribution of 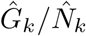 shifts towards a monomodal, with a peak close to 0.5. We used *n*_*C*_ = 80 (estimated from data, see SI appendix Section 2) interacting compartments. Experimental stirring wass performed for 1 min at four different strengths (2.0, 5.2, 16, 30*krpm*). For each stirring strength, we evaluated the stirring parameter *s* yielding a theoretical distribution corresponding the experimental distribution. **C** Effect of stirring on the probability distribution function of the fraction 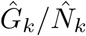 (probability that 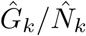 is between *u* and *u* + *du*) with *n*_*C*_ = 80 interacting compartments, starting from a perfectly bimodal distribution with peaks at 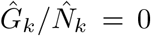 and 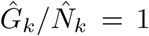. The distribution shifts towards a monomodal distribution as *s* increases.

This rescaling assumes that during typical fusion/division events, droplets exchange a number of fluorescent molecules of the order of the average fluorescence per compartment. We call 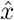 the fraction of fluorescence intensity due to green fluorescence in one compartment, which serves as a proxy for the fraction of G in the compartment, and we study the cumulative distribution function of 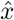(the probability of having 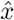lower than a given value *x*). In practice, we employ 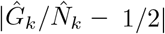 to correct for experimental deviations toward either 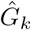 or 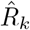, introduced during microscopic observation. Values close to 0 indicate well-mixed compartments containing approximately equal amounts of G and R, whereas values close to 0.5 indicate compartments that contain either G or R almost exclusively. We found that the fraction of droplets with similar amounts of G and R gradually increased with stronger stirring and over two experimental rounds (Fig. 4B and Fig. S13).

To model the observed shift in the distribution, we start with a bimodal distribution of 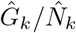, assuming that compartments initially contain either only *G* or no *G*, and use the experimental distribution of 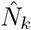 . We then apply a theoretical transformation to describe the effect of stirring on these distributions, yielding the final distribution conditioned on the initial state (see Materials and Methods, section B for details). This transformation depends primarily on the stirring parameter *s* and an effective number of compartments *n*_*C*_. We calibrate *n*_*C*_ by assuming that the maximal stirring condition (30 krpm) corresponds to *s* →1, as detailed in SI appendix Section 2B. Additionally, we confirm in SI appendix Section 2 that the transformed distribution approaches a fixed point as *s*→ 1, consistent with the distribution of *N*_*k*_, where *s* = 1 represents maximal stirring. The impact of this transformation on the probability density of 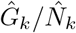 is shown in Fig. 4C. With increasing stirring strength, the distribution transitions from a bimodal to a centered unimodal form.

In Fig. 4B we compare the result of this transformation, starting from an initially bimodal distribution, expressed as the cumulative distribution of 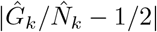 (solid lines for different values of *s*), with the experimental distributions after round 1 (dotted lines), and obtain a good agreement between them. In practice, increasing the number of effective compartments *n*_*c*_ renders the distributions sharper (due to central limit theorem). Here we use an effective number of compartments of *n*_*C*_ = 80 (estimated from data as explained in SI appendix Section 2), and by calibrating *s* = 1 to 30 krpm, we associate a value of *s* with each stirring strength. We observe a good agreement between our theory and experimental data, with both showing a shift of the distribution towards a more monomodal form. Accordingly, our parameter *s* captures the effect of stirring on droplet contents. We also recover that weak stirring (less than 5.2 krpm) corresponds to small values of the stirring parameter *s* ≤ 0.4, whereas stronger stirring at 16 krpm and 30 krpm correspond to *s* ≥ 0.5. We confirmed the robustness of these estimations by applying the same transformation and stirring parameters *s* to experimental distributions after round 2, which are successfully reproduced by our model (Fig. S11). The correspondence between stirring strengths and stirring parameters *s* has been used to plot the data for *x*_*tot*_ against *s* in Fig. 3A.

### Survival and predominance

Our theoretical description of stirring can now be used to study how this process impacts the dilution and maturation in the replicase-parasite dynamics. We study various networks of interactions between replicases and parasites. To this end, we build coexistence diagrams representing the expected fractions of each co-existing species in SI appendix Section 4. In Figs. S15-18, we build these diagrams for models A and B. In both cases, a higher dilution factor favors replicases by isolating parasites more efficiently. In model B, a longer maturation time also favors parasites due to more frequent mutations from replicases to parasites. In Figs. S20-23, we examine how these coexistence diagrams depend on stirring. As detailed above, stronger stirring typically favors replicases, although it can also lead to extinction if *K/d* is sufficiently small due to the finite number of compartments. In Figs. S25-26, we observe that generalist parasites frequently take over specialist parasites. In Fig. S27, we explore the case where mutant replicases acquire a better resistance to parasites but do not improve their replication rate. In Figs. S28-29, parasites acquire generality but their replicability decreases; nevertheless, generalist parasites remain favored.

We observe that weaker stirring tends to favor parasites, as concluded in [37] and detailed above in the present work. We recover the complete pooling limit for *s* →1. If the system has a strong compositional memory, dilution only weakly affects the system, whereas mutations occur even if the compartment is not diluted. With a weaker stirring, parasites are favored (as predicted in [37]), because parasites would less likely be isolated from replicases.

When the number of compartments is small enough, we may observe extinction of the whole population in the regime of strong dilution, provided that stirring is also strong enough. Under weak dilution, the heterogeneity in the number of individuals per compartment allows some compartments to contain more than one individual even if *K/d* is smaller than 1.

## Discussion

In this work, we investigated the role of transient compartmentalization on the coevolution of parasites and replicases. The theoretical framework we have built is independent of the nature of the compartments, which may be for instance membrane-less aggregates, coacervates or droplets. We find that the capacity of compartments to inherit from their past compositions (compositional memory) matters for the dynamics of these early molecular systems in agreement with the idea of compositional genomes or compositional memory [21, 22, 23, 20].

We note that our experimental RNA replication system relied on a modern translation machinery, and replication occurred using encoded proteins. Such a complex replication scheme is unlikely to have existed in early stages of the origins of life. Unfortunately, more primitive RNA-based systems, such as ribozymes capable of sustained co-replication of replicases and parasites, are not yet available. The development of such systems would enable a more direct test of our theoretical model in a prebiotically relevant context. Nevertheless, the experimentally observed replicase-parasite dynamics were generally consistent with theoretical predictions based on simpler RNA replicators (Fig. 2D, 3A), suggesting that underlying compartmental effects may be broadly applicable. In addition, our findings may provide insights into the development of evolvable artificial cell systems in the context of bottom-up synthetic biology [47, 48]. Genomic replication has been reconstituted through multiple mechanisms in diverse compartment types, including water-in-oil droplets, liposomes, and liquid–liquid phase-separated droplets, and compartmentalization has been shown to suppress parasite amplification [49, 50, 51]. Although long-term coevolution of replicases and parasites has so far been demonstrated only in specific systems [34, 38], similar dynamics may be achievable in other platforms if compartment mixing is appropriately controlled.

Compartments allow the coevolution of replicases and parasites by preventing a complete takeover by parasites. It results in complex dynamics, with oscillations between four species that we can predict theoretically and that significantly differ from the behavior expected in bulk conditions. Dilution and carrying capacity play a central role in the ability of replicases to be maintained in the population during evolution. Transient compartmentalization dynamics are only possible if the system undergoes mixing steps, where parasites may be spatially isolated from replicases, resulting in their inability to replicate. Compositional memory is observed experimentally, and its effect on the predominance of parasites has been assessed. We also quantified the relation between the strength of stirring and the mixing of compartments, thanks to a combination of experiments and modelling. In future work, it will be important to further investigate this issue by systematically tracking the composition of individual compartments in experiments with compartmentalized RNAs. We hope that our work will guide further experiments in this field, as more complex and specific interactions between RNA molecules, as well as between RNA and proteins, will need to be included in future extensions of this work.

## Materials and methods

In this section, we provide details regarding the evolution of molecular replicators undergoing transient compartmentalization dynamics and detail experimental conditions.

### RNA molecules

All RNA molecules (HL1-, HL2-, PL2-, and PL3-228) were prepared by in vitro transcription. First, the DNA template for each RNA was amplified by PCR using the corresponding plasmid obtained previously [34]. The resulting DNA products were digested with Sma I (Takara), transcribed in vitro using T7 RNA polymerase (Takara), and treated with DNase I (Takara). The transcribed RNAs were purified using the RNeasy Mini Kit (Qiagen).

### Compartmentalized RNA replication system

The RNA replication system was prepared by mixing an arbitrary set of RNA molecules in 10 *µ*L of the Escherichia coli reconstituted translation system [52], as used previously [34]. To encapsulate the replication system in water-in-oil droplets, the 10 *µ*L mixture was stirred in 1,000 *µ*L of buffer-saturated oil at various speeds for 1 min on ice using a homogenizer (Polytron PT-1300D, Kinematica). The buffer-saturated oil was prepared as follows. An oil phase composed of 95% mineral oil (Sigma-Aldrich), 2% Span 80 (Nacalai Tesque), and 3% Tween 80 (Fujifilm) was mixed with 1/20 vol. of the translation system buffer, lacking NTPs, tRNA, and translation proteins but containing an increased concentration of dithiothreitol (36 mM). After vigorous mixing, the oil mixture was incubated at 37°*C* for 10 min, centrifuged at 20,000 g for 5 min at room temperature, and the upper oil phase was collected as buffer-saturated oil.

### Long-term RNA replication experiment

The RNA replication system was prepared with 10 nM each of HL1-, HL2-, PL2-, and PL3-228. The mixture was emulsified in buffer-saturated oil at 16 krpm for 1 min to generate water-in-oil droplets. The droplets were incubated at 37°*C* for 5 h to induce RNA replication coupled with protein translation, completing one replication round. For the next round, 200 *µ*L of the droplet suspension was transferred to a new mixture containing 800 *µ*L of buffer-saturated oil and 10 *µ*L of the translation system, and homogenized as above. The new droplets were then incubated again at 37°*C* for 5 h. These cycles were repeated for 22 rounds. RNA concentrations were determined at the end of each cycle (after 5 h incubation) by intercalating dye-based RT-qPCR using One Step TB Green PrimeScript PLUS RT-PCR Kit (Takara) and sequence-specific primers. The primers were designed to specifically amplify each RNA species, and sequence mismatches with non-target RNAs prevent cross-amplification (Table S4). RNA concentrations were quantified using dilution series of each RNA.

### Competition assay

The RNA replication system containing 0.05 nM HL2-228 and 0.15 nM PL2-228 was used to prepare water-in-oil droplets via homogenization at 16 krpm for 1 min as described above. To vary the dilution conditions, 10, 40, 200, or 500 *µ*L of the droplet suspension was mixed with 990, 960, 800, or 500 *µ*L of buffer-saturated oil and 9.9, 9.6, 8, or 5 *µ*L of the translation system, respectively. The mixtures were then stirred for 1 min at 2, 5.2, 16, or 30 krpm using the same homogenizer. After incubation at 37°*C* for 5 h, RNA concentrations were measured by RT-qPCR as described above using sequence-specific primers (Table S3).

### Evaluation of content mixing in droplet populations

Fluorescein isothiocyanate (FITC)-labeled dextran (FITC-dex, G) and tetramethylrhodamine isothiocyanate (TRITC)-labeled dextran (TRITC-dex, R), with average molecular weights of 10 kDa and 155 kDa, respectively, were purchased from Sigma-Aldrich. Alexa Fluor 647-labeled dextran (Alexa-dex; 10 kDa) was purchased from Invitrogen. Using these fluorescent dextrans, two types of waterin-oil droplets were prepared: droplets 1 containing 3 wt% FITC-dex and droplets 2 containing 3 wt% TRITC-dex, as described above but using a translation buffer lacking protein components. To examine droplet fusion induced by mechanical agitation, 100 *µ*L each of droplets 1 and 2 were mixed with 800 *µ*L of buffer-saturated oil and 10 *µ*L of the translation system buffer containing 0.01 wt% Alexa-dex. The droplet mixtures were then stirred for 1 min at 2, 5.2, 16, or 30 krpm, as described above. The droplet populations were then incubated at 37°*C* for 5 h, completing the first round of the experiment. For the second round, new droplet populations were prepared in the same manner using 200 *µ*L of each post-incubated droplet suspension in place of droplets 1 and 2, followed by incubation at 37°*C* for another 5 h. At 5 h of rounds 1 and 2, aliquots were collected for microscopic observation. Fluorescence imaging was performed using an FV3000 confocal laser-scanning microscope (Evident) equipped with a 100× oil-immersion objective at room temperature. FITC, TRITC, and Alexa Fluor 647 were excited at 488 nm and 561 nm, and 640 nm, respectively. Images were analyzed using ImageJ (NIH). In merged triple-channel images, signals with circularity ≥0.7 were identified as individual droplets, and the mean intensities of each fluorophore were recorded for every droplet. Background intensities were calculated as the mean fluorescence of droplet-free regions. Misidentified droplets (based on transmitted light images) were manually excluded. To focus on “active” droplets containing either FITC or TRITC fluorophore, droplets with both FITC and TRITC intensities below 105% of their respective background intensities were omitted as unreliable detections. The ratio of FITC to TRITC intensity was then calculated for each remaining droplet after background subtraction. Droplet size was estimated by assuming circular geometry.

### Perfectly homogeneous limit

Let us consider a pool of replicators, with a fraction *x*_*k*_ of replicases and *z*_*k*_ a fraction of parasites. In the inoculation step, compartments are created with a total population drawn from a Poisson distribution of parameter *λ*, and a number for each species drawn from a multinomial distribution of parameters *x*_*k*_ and *z*_*k*_. Therefore, the probability to have a compartment with *m*_*k*_ replicases of type *k* for 0≤ *k* ≤*K* (modeled by a vector ***m***) and *y*_*l*_ parasites of type *l* for 0≤ *l* ≤*L* (modeled by a vector ***y***) is

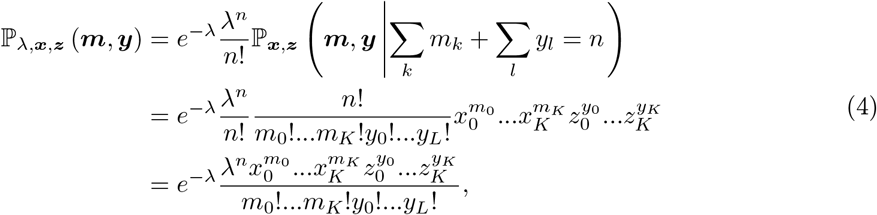

where *n* = ∑_*k*_ *m*_*k*_ +∑ _*l*_ *y*_*l*_. This step is a significant source of stochasticity and heterogeneity among compartments and the only one considered in the deterministic approach.

#### Maturation (model A)

A simple model consists in assuming that the sequence of the replicases mutates with time, in which case we can index the mutations by an integer *k* (similar mutants are grouped together). We also suppose that new types of parasites can appear. The fitness of replicases corresponding to mutant *k* replicating with replicase *j* is *α*_*j,k*_, the fitness of parasites of type *l* replicating with mutant *j* is *γ*_*j,l*_. With ***e***_*i*_ the vector of 0 except at entry *i*, we can write the master equation for the probability of having ***m, y*** at time *t* in a given compartment in Eq. 5 (see Fig. S1).

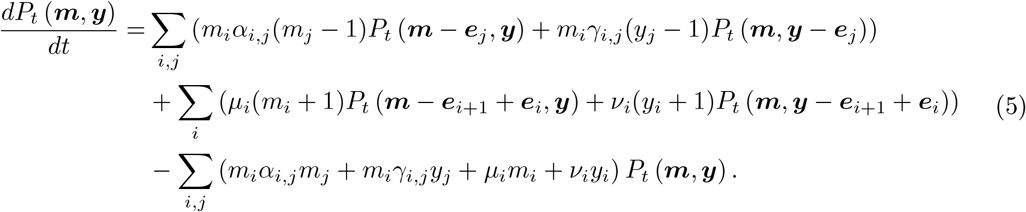

The system evolves according to this master equation until the population reaches the carrying capacity of the compartment. The carrying capacity can be thought of as the total number of molecules or as the total number of nucleotides (it depends on the molecular lengths in this case) that can be produced. The latter effect is explored in SI appendix Section 11. The processes that can occur during maturation are detailed in Fig. S1. The model is similar to that of [31] except that it accounts for the possible coevolution of replicases and parasites through mutations.

#### Maturation (model B)

In this model, we assume that parasites appear from deletions in the sequence of replicases (by recombination for instance [34]), yielding smaller sequences unable to replicate by themselves. The notations remain the same in this model. The mutations rates of the evolution equations of the maturation phase are different, and we obtain the master equation in Eq. 14 (see Fig. S2). The replication dynamics are the same in both models and we typically have that *m*_0_(*t*) = *m*_0_*/*(1 −*α*_0,0_*m*_0_*t*) during this phase. In the following, we will study the differences that result from the choice of the model.

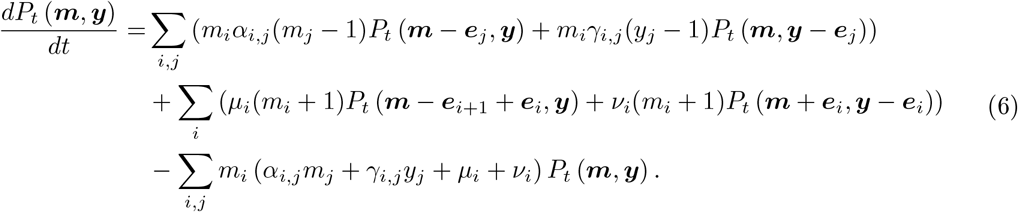

### Modeling the dye experiment

Let us call the initial distribution of fraction *x* of G in the droplets *p*_*ini*_(*x*). After stirring, we write the cumulative probability of having a fraction 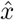 lower than *x* in a compartment *k*, using the mean field approach described above. We denote *G*_*k*_ the rescaled amount of G in compartment *k, N*_*k*_ the total rescaled fluorescence in compartment *k, n*_*c*_ the number of interacting compartments and *N* the total population of the system. We also assume that instead of taking the fraction of G as a single random variable, we have the following rules for mixing

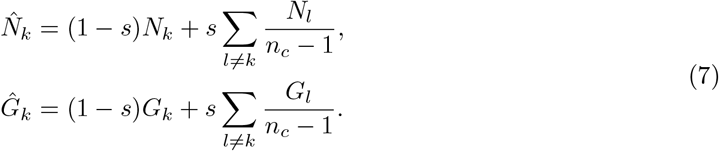

Naturally, this transformation conserves the numbers of molecules of each type because 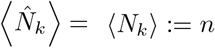 (with standard deviation *σ*_*N*_) and 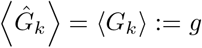 (with standard deviation *σ*_*G*_). Note also that the random variables *N*_*k*_ and *G*_*k*_ are correlated due to the fact that *N*_*k*_ = *G*_*k*_ + *R*_*k*_. The cumulative probability distribution of the updated fraction after one round of mixing is

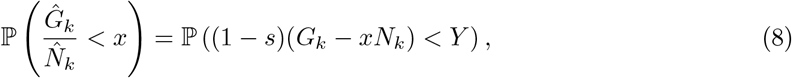

with 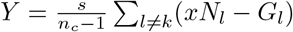.

Now, we can approximate the sums ∑_*l*≠*k*_ *G*_*l*_ and ∑_*l*≠*k*_ *N*_*l*_ as normally distributed, using the central limit theorem in the limit of a large number of compartments. Therefore, accounting for the fact that *G*_*l*_ and *N*_*l*_ are correlated we obtain

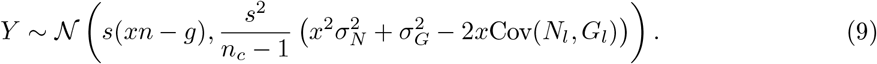

For which we can use the measured values of standard deviations and covariances (from experimental distributions). If we assume that the distribution is initially bimodal in *G*_*k*_ (initially the compartments are either full of G, or contain no G), we can also compute the statistics of *G*_*k*_ as a function of the experimental statistics 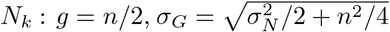 and 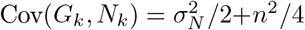. Now, we can write the cumulative distribution probability, using the law of total probability:

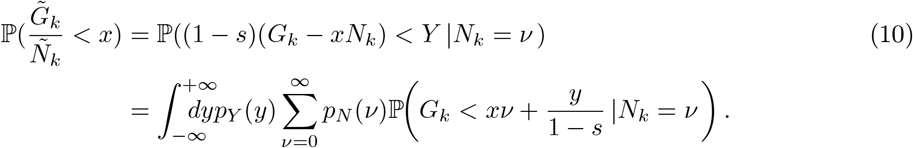

This formula depends on the initial cumulative probability, standard deviations and covariance of the distributions of *N*_*k*_ and *G*_*k*_. In Fig. 4B, we used initial experimental distributions to obtain these values and compute the theoretical cumulative distribution after stirring for different values of *s* (shown in solid lines). In the particular case where we start from a perfectly bimodal distribution for the conditional probability, we have *p*_*G,ini*_(*γ*|*N*_*k*_ = *ν*) = (*δ*_*γ*,0_ + *δ*_*γ,ν*_)*/*2, and the experimental distribution for *N* . Similarly we express the cumulative distribution function of 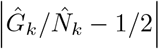 in SI appendix Section 2. A closed expression for the cumulative distribution of the fraction of 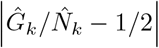 follows after one round of stirring that is compared with the experimental data in Fig. 4B.

## Acknowledgments

We thank Taro Furubayashi, Hidekazu Sono, and Philippe Nghe for helpful discussions. We thank OIST and the Thematic Program on the “Computational and Physical Understanding of Biological Information Processing” for their support. This research was supported by JSPS KAKENHI (25K22442 to R. M.

---

**Table S1.** Notations for species and dynamical parameters.

|  |  |
| --- | --- |
| $m_0$ | Number of wild-type replicases in a compartment |
| $m_k$ | Number of mutant replicases of generation $k$ in a compartment |
| $y_0$ | Number of wild-type parasites in a compartment |
| $y_k$ | Number of mutant parasites of generation $k$ in a compartment |
| $n$ | Total number of individuals in a compartment |
| $x_0$ | Fraction of wild-type replicases in the whole population |
| $x_k$ | Fraction of mutant replicases of generation $k$ in the whole population |
| $z_0$ | Fraction of wild-type parasites in the whole population |
| $z_k$ | Fraction of mutant parasites of generation $k$ in the whole population |
| $K_r$ | Carrying capacity of replicases in one compartment |
| $K_p$ | Carrying capacity of parasites in one compartment |
| $\alpha$ | Matrix of replication rates for replicases |
| $\gamma$ | Matrix of replication rates for parasites |
| $\mu$ | Vector of mutation rates of replicases |
| $\nu$ | Vector of mutation rates of parasites |
| $\bar{u}$ | Value of a quantity $u$ after maturation |
| $\hat{u}$ | Value of a quantity $u$ after stirring |

## Appendix

### Transient compartmentalization models

We model the evolution of a population (mutations following fitness gradient) along with transient compartmentalization dynamics. We want to describe the evolutions of replicases and parasites, mutating to modify their interactions or increase their relative fitness. We introduce the notations in Table 1

#### Perfectly homogeneous limit

##### Compartmentalization

Starting from a pool of individuals, with fractions *x*_*k*_ and *z*_*k*_, compartmentalization consists in creating compartments with a total population drawn from a Poisson distribution of parameter *λ* (which is itself a dynamical parameter, modified at each cycle), and the number of each species drawn from a multinomial distribution of parameters *x*_*k*_ and *z*_*k*_. Therefore, the probability to have a compartment with *m*_*k*_ replicases of type *k* for 0 ≤ *k* ≤ *K* (modeled by a vector ***m***) and *y*_*l*_ parasites of type *l* for 0 ≤ *l* ≤ *L* (modeled by a vector ***n***) is

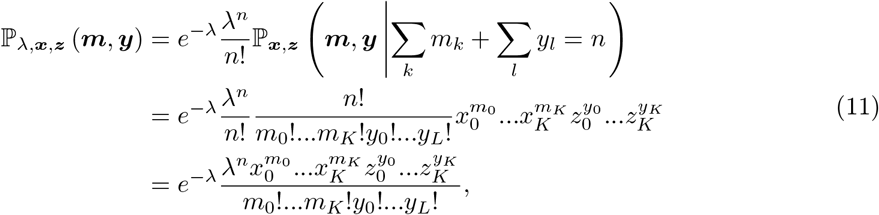

where *n* = ∑_*k*_ *m*_*k*_ + ∑ _*l*_ *y*_*l*_. This is the product of the probability to have *n* individuals in the compartment and the probability to have ***m*** replicases and ***y*** parasites given there are *n* individuals. This step is a source of stochasticity and heterogeneity among compartments. As explained in the main text, the initial condition in a compartment will determine its final composition in the deterministic approach.

##### Maturation (model A)

A simple model consists in assuming that the sequence of replicases mutates with time, in which case we we can index the mutations by an integer *k* (similar mutants are grouped together). We also suppose that new types of parasites can appear. The fitness of replicases corresponding to mutant *k* replicating with replicase *j* is *α*_*j,k*_, the fitness of parasites of type *l* replicating with mutant *j* is *γ*_*j,l*_. We can write evolution equations for the maturation phase (without noise) :

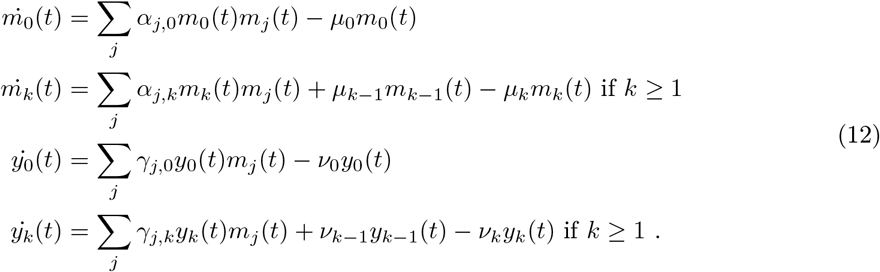

The processes that can occur during maturation are detailed in Fig. 1. We can also describe this phase in terms of a Master Equation for the probability of having ***m, y*** at time *t* in a given compartment. For this we introduce ***e***_*i*_ the vector of 0 except at entry *i* and we have

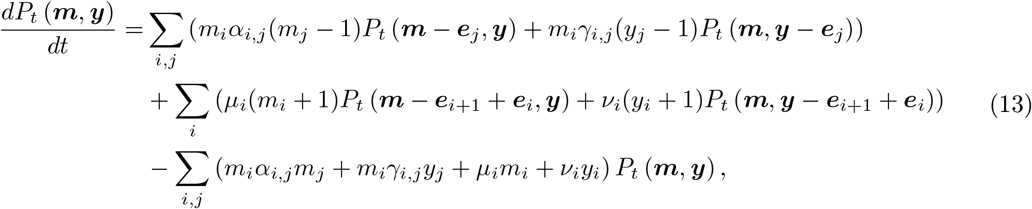

which will be used in the Gillespie simulations.

##### Maturation (model B)

In this model, we assume that parasites appear from deletions in the sequence of replicases, yielding smaller sequences unable to replicate by themselves. The mutations rates of the evolution equations of the maturation phase are different (see Fig. 2) and without noise we have

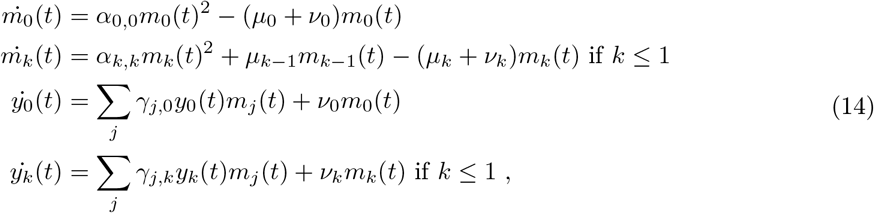

**Figure S1.**
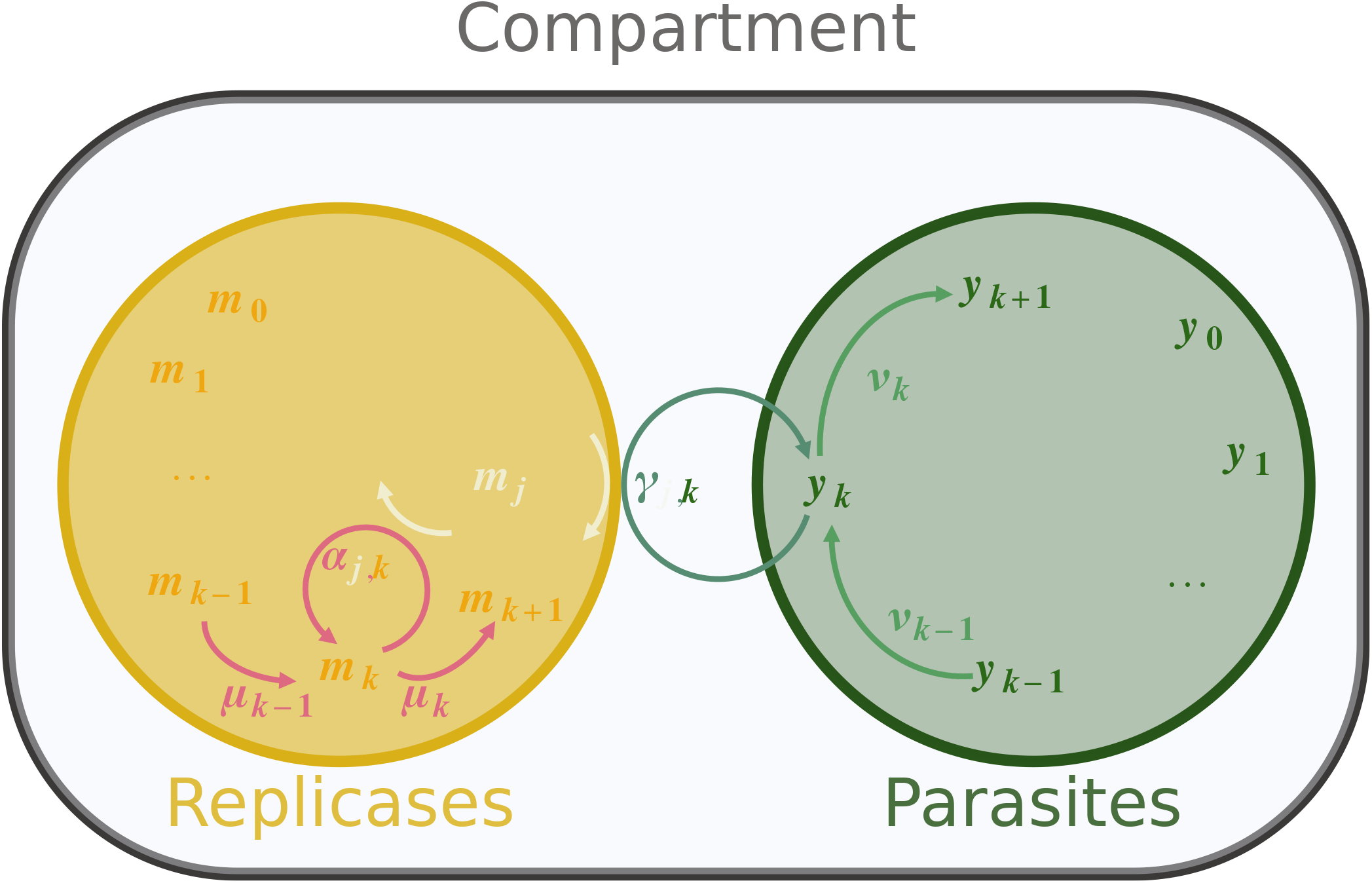
Processes occuring during maturation (model A). Replicases of type 0, 1, …, *k* + 1 are shown on the left (yellow disk) and parasites of type 0, 1, …*k* + 1 on the right (green disk). Replicase *k* can be replicated by replicase *j* at a rate *α*_*j,k*_, and mutate to replicase *k* + 1 at a rate *µ*_*k*_. Parasite *k* can be replicated by replicase *j* at a rate *γ*_*j,k*_, and mutate to replicase *k* + 1 at a rate *ν*_*k*_.

**Figure S2.**
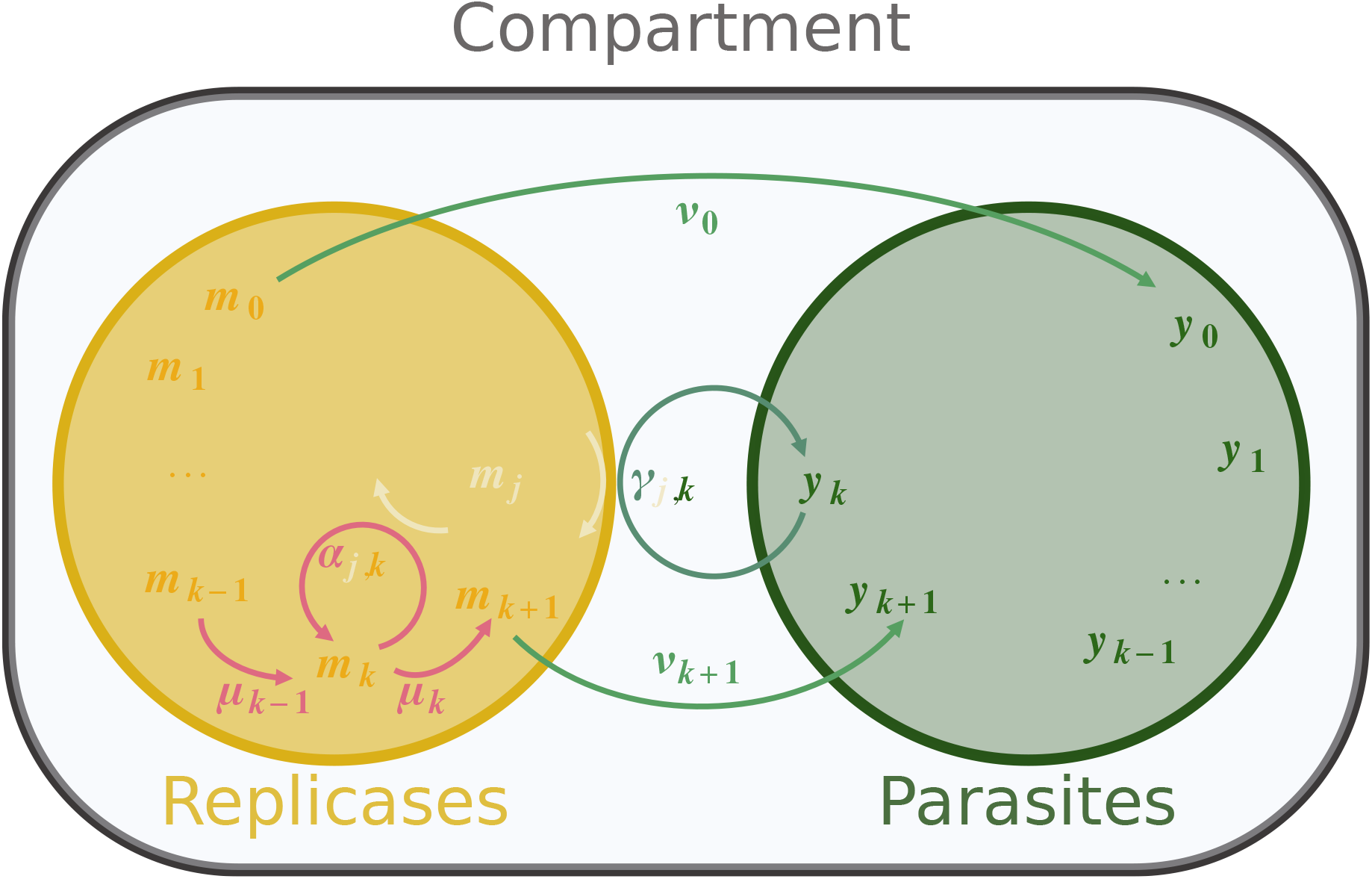
Processes occurring during maturation (model B). Replicases of type 0, 1, …, *k* + 1 are shown on the left (yellow disk) and parasites of type 0, 1, …*k* + 1 on the right (green disk). Replicase *k* can be replicated by replicase *j* at a rate *α*_*j,k*_, and mutate to replicase *k* + 1 at a rate *µ*_*k*_, and to parasite of type *k* at a rate *ν*_*k*_. Parasite *k* can be replicated by replicase *j* at a rate *γ*_*j,k*_.

the replication dynamics are the same in both models and we typically have that *m*_0_(*t*) = *m*_0_*/*(1 − *α*_0,0_*m*_0_*t*) during this phase. In the following, we will study the differences that result from the choice of the model. We can also write a Master equation for the probability of having ***m, y*** at time *t* in a compartment

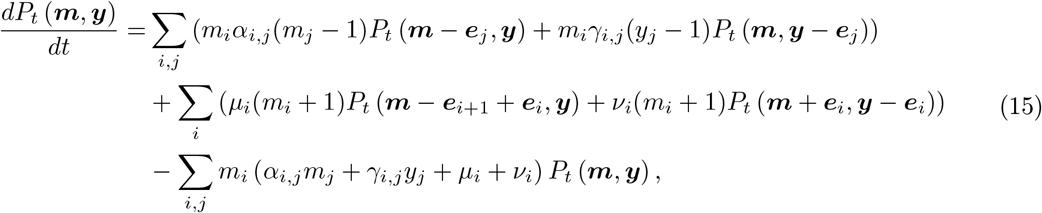

which we use in the Gillespie simulations.

#### Deterministic approximations

The deterministic maturation phase is the same for both models, which only differ in the mutation dynamics. We assume that the initial condition completely determines the post-maturation state, and this reasoning is true for both models. This allows to explore the dynamics of this process even if we cannot find analytic solutions to the non linear maturation equations. There are two sources of stochasticity in this process:

- A first source of stochasticity is due to inoculation, i.e. the preparation of the initial composition of the compartment using molecules present in the pool.
- A second source of stochasticity derives from the maturation phase, during which each compartment evolves stochastically through mutation or replication events.

In the deterministic approach, the maturation of a compartment is fully determined by its initial composition, which means that the only source of stochasticity in the system is the inoculation step. This hypothesis is key in order to simplify the theoretical description of the dynamics since the evolution equations during the maturation phase are highly non-linear. We make first make the assumptions detailed in the main text and summarized in Fig. 3. Then we study other the effect of different deterministic assumptions. As a result, only one final composition is associated to a given initial condition. In the following, we use this deterministic model and study different limits, obtained by comparing the different rates. In general, to study the number 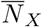 of one species *X* per cell after maturation (starting from a number *N*_*X*_), we use the assumption that the maturation time is long enough for every compartment to reach its carrying capacity (providing it contains replicases, and where the carrying capacity *K*_*X*_ may depend on wether hosts or parasites are favored):

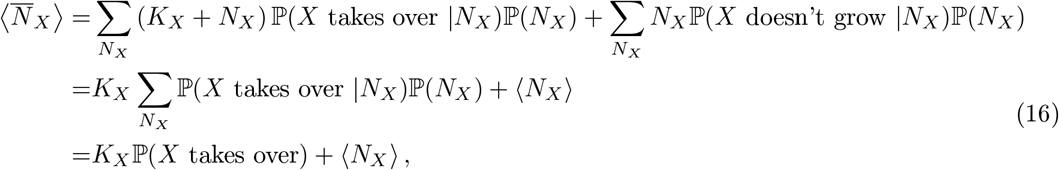

where the probabilities and averages are taken with respect to the initial distribution within the compartments.

If mutations are beneficial, we can compute the updated fraction of each species after one step following the above method:

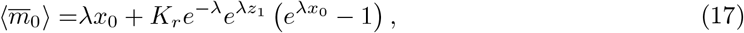

where *K*_*r*_ is the carrying capacity for replicases. Similarly, we find:

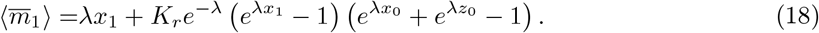

To have a complete description of the system, we need to compute 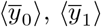. In particular:

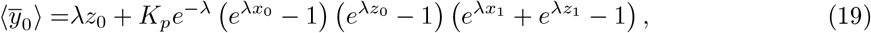

with *K*_*p*_ the carrying capacity for parasites, and similarly:

**Figure S3.**
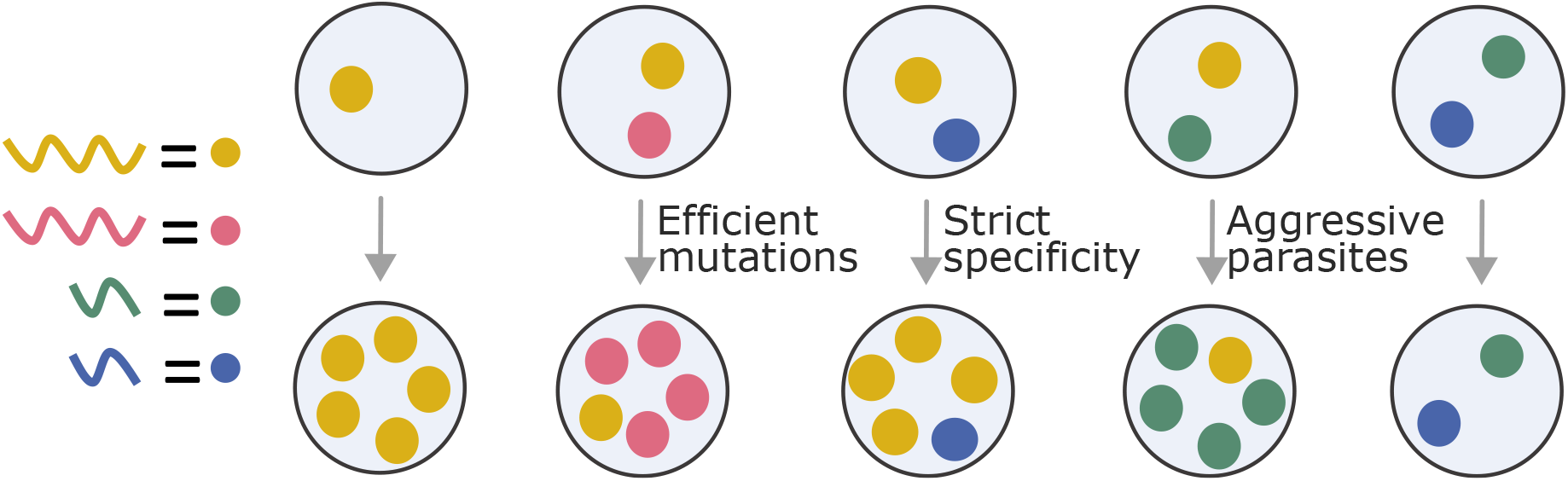
Deterministic approach of the maturation phase. A replicase isolated in a compartment will be able to be replicated freely. In a compartment with different types of replicases, the fittest will take over the compartment, and we assume that fitness increases with mutations (*efficient mutations*, otherwise we can switch the roles of mutants and WT to sort them according to their fitness). In a compartment with a replicase and a parasite which is not able to use this replicase, parasites will not be replicated (*strict specificity*). In a compartment with a replicase and parasite able to use this replicase, parasites will take over (*aggressive parasites*). In a compartment with parasites only, parasites will not be replicated.

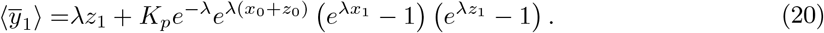

and we have:

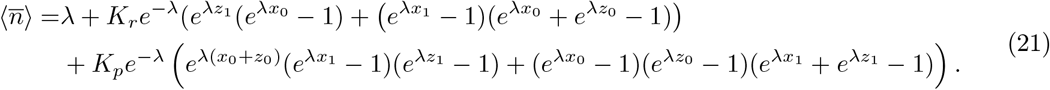

Therefore, we recover:

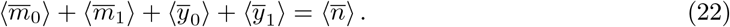

#### Mutations

##### Modeling mutations (model A)

The major issue with the deterministic approach developed above is that mutations are not directly accounted for. Even if it is enough to capture the dynamics of evolution in multiple cases, some processes cannot be described with this formalism. One way to improve the deterministic model is to assume that during each maturation phase, a number *δ*_*m*_ mutant replicases (*δ*_*y*_ mutant parasites) is created from the wild-type populations (this will depend on the model). We can estimate the number of created mutants by comparing the mutation rate and replication rate. In addition, we found that the evolution of the wild-type populations during maturation can be described as:

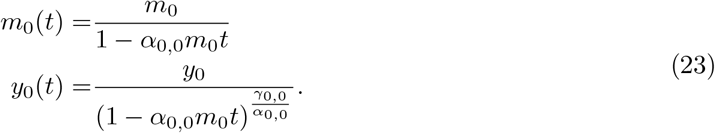

For model A, we can estimate, if wild-type replicases are favored using that 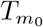 is the time to reach the carrying capacity:

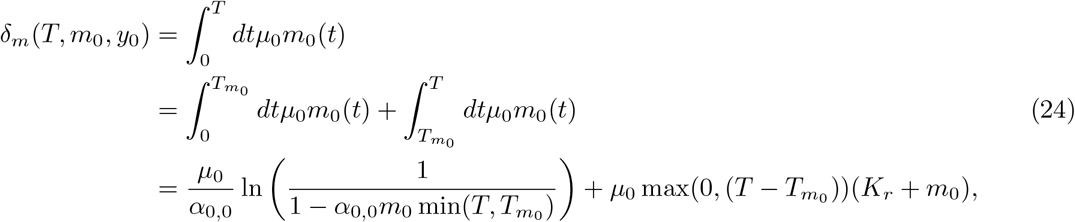

where the min in the first term represents the fact that if 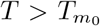 the population reaches the carrying capacity, and then the second term of the sum is different from 0. For too small maturation times, the second term is null. If the population of wild-type replicases is not growing (not favored):

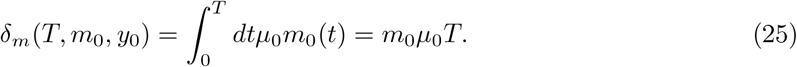

Therefore, if we consider the timescale of the maturation phase to be the time to reach the carrying capacity in a compartment starting with *m*_0_ = 1 and *n* = 1, 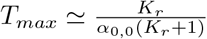:

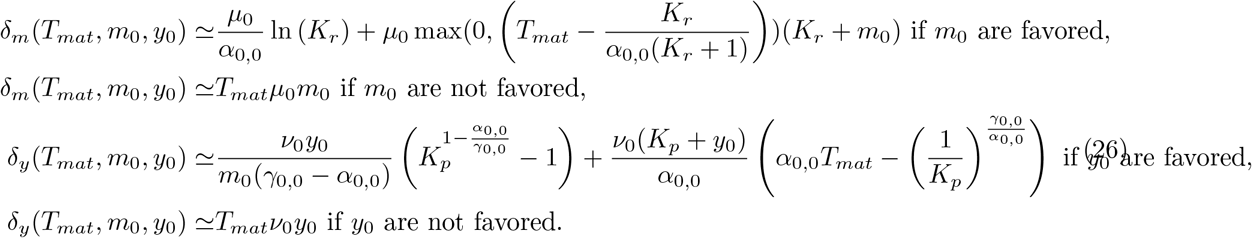

In addition, *m*_0_ are favored if and only if *m*_0_ *>* 0 and *y*_0_ = 0 and *m*_1_ = 0. Similarly, *y*_0_ are favored if and only if *y*_0_ *>* 0 and *m*_0_ *>* 0 and (*m*_1_ = 0 or *y*_1_ = 0). This is true when we do not consider the effect of the carrying capacity, possibly limiting growth when the compartment is close to being full. Therefore, the typical time to reach the carrying capacity in a compartment, starting from *m*_0_ and without parasites, is 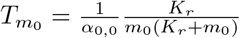. Similarly we find:

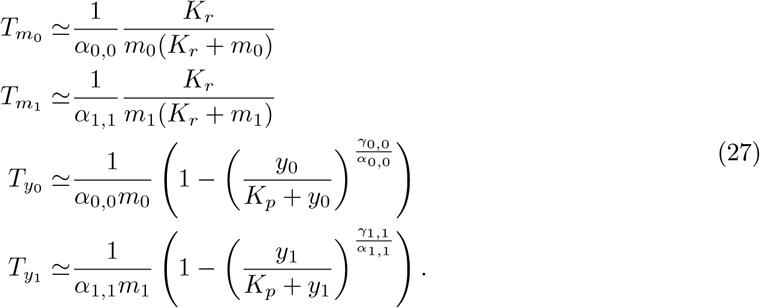

Due to the assumption of beneficial mutations, *α*_0,0_ ≪ *α*_1,1_, thus 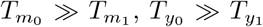 and thanks to the aggressive parasite assumption, 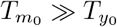. Therefore, the time of a maturation phase should be at least 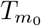. We can thus compute the average numbers of added mutants and removed wild-type individuals (due to mutations) for each species :

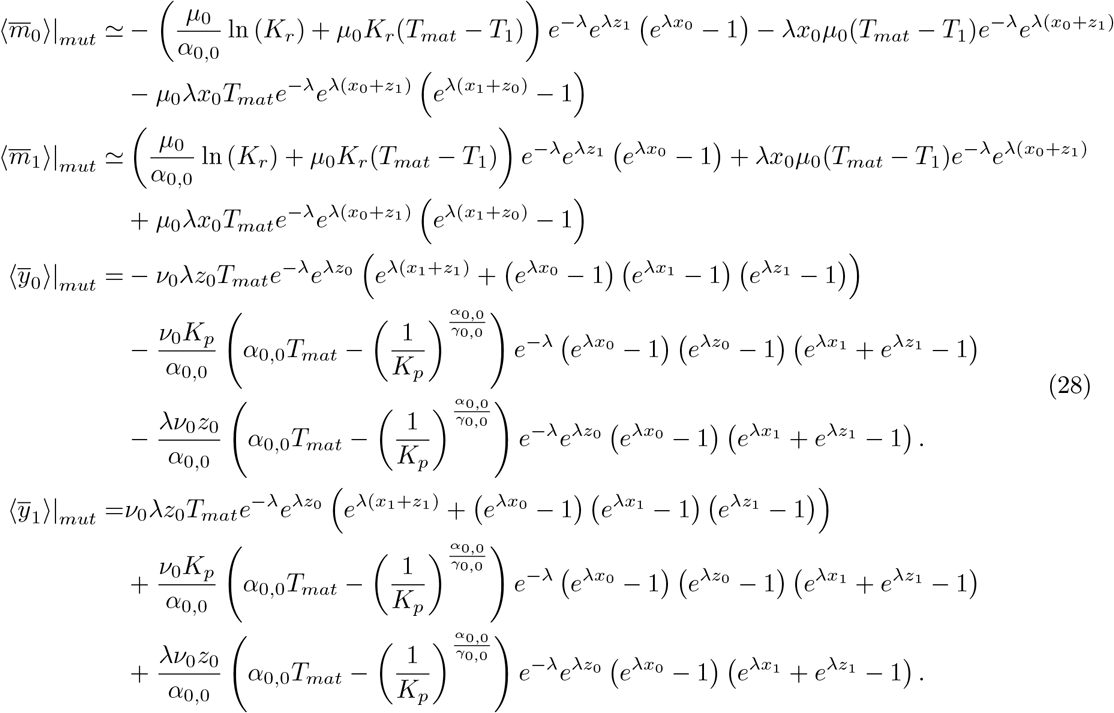

We can now simulate the whole coevolution in this simple deterministic approach and observe the effect of mutations. In addition, we compare this theory (solid line) to Gillespie simulations (dotted) in Fig. 4. The Gillespie simulations incorporate the non-linearities of maturation and preserve all sources of stochasticity. In particular, we observe that the deterministic model describes precisely the stochastic behaviour in most conditions, apart from the noise. With low probability of mutations, mutations become rare events with strong influence on the evolution. In this case the deterministic approach does not perform as well because the system is strongly driven by the first occurrence of mutation. Yet theory accuratelyely predicts the average fractions of each species even in this limit. This is however an effect due to the finite number of compartments in the simulations, and we can show that for larger numbers of compartments the simulations match the theory even in that case of rare mutations.

##### Modeling mutations (model B)

The main difference from the previous case is that parasites do not mutate directly. Replicases also mutate to form parasites, and this mechanism could favour coevolution. Indeed, we observe in Fig. 5 that more species seem to coexist in this situation. Again, Gillespie simulations agree well with the theory, up to a finite-size effect resulting in the noise observed in the simulations. As for model A, the discrepancy between theory and experiments increases as the mutation rates become smaller, due to the fact that mutations occur stochastically and wild-type populations can go extinct contrary to predictions from the deterministic model. In this case, mutant replicases accumulate because mutant parasites mutate too slowly to limit this effect, leading to the extinction of the wild-type populations in the simulations. We obtain the following numbers of mutants

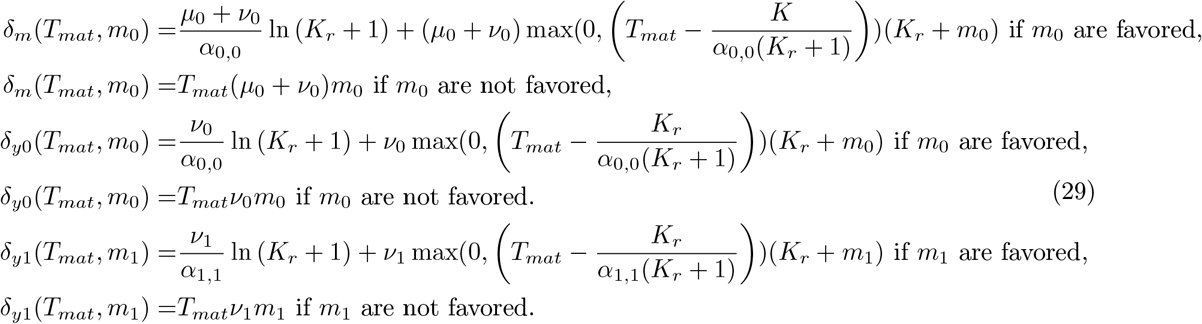

And by averaging, we find that:

**Figure S4.**
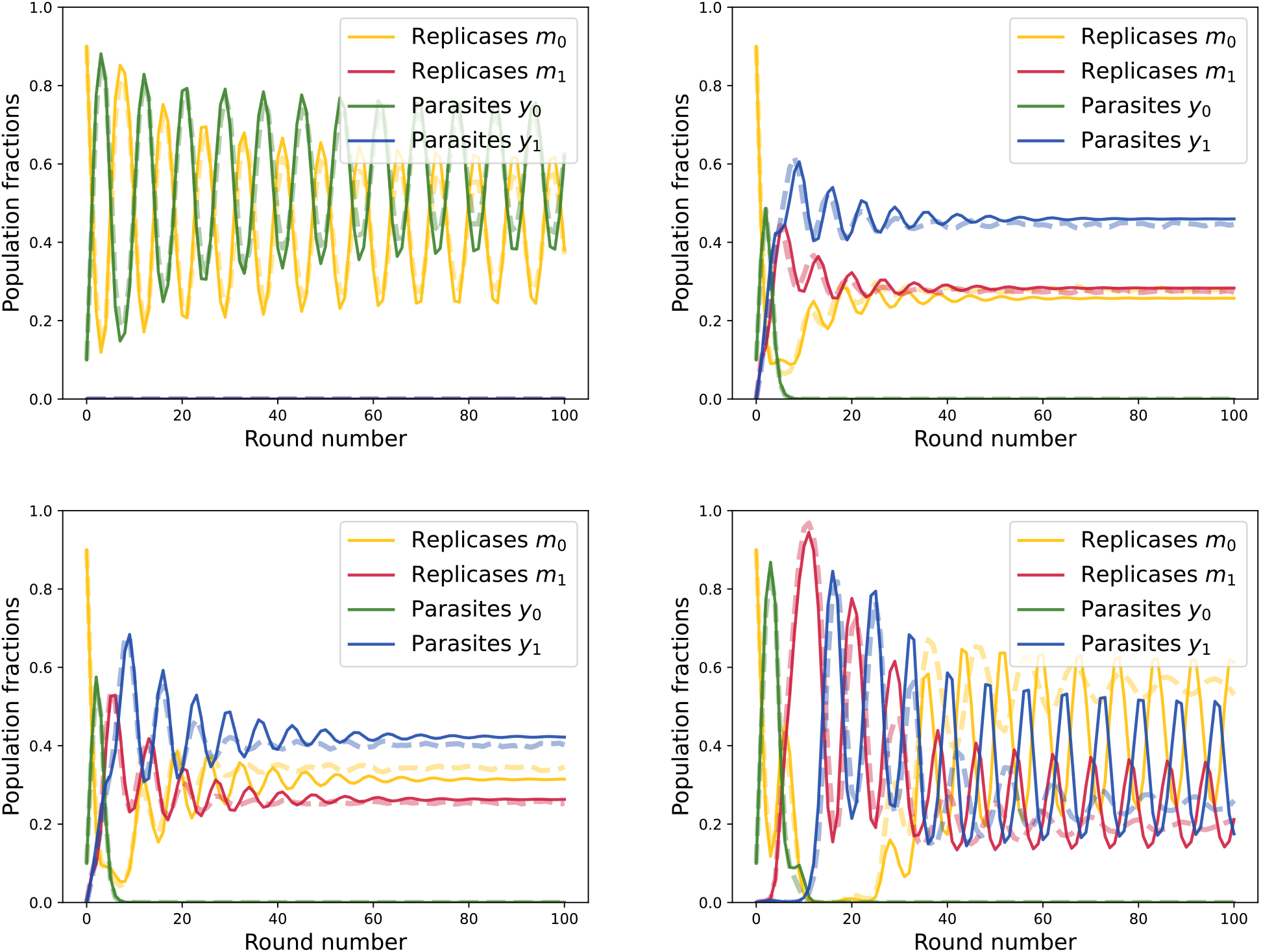
Theoretical prediction for model A (full) of the evolution of the system compared to Gillespie simulations (dotted) performed with 50000 compartments. We used for replication rates *α*_0,0_ = 5 × 10^−1^ *h*^−1^, *α*_1,1_ = 2 *h*^−1^, *γ*_0,0_ = 7 *h*^−1^, *γ*_1,1_ = 20 *h*^−1^. The carrying capacities are *K*_*p*_ = *K*_*r*_ = 30 and dilution factor *d* = 10. The different panels correspond to different mutation rates. **Upper left** *µ* = *ν* = 0 *h*^−1^. **Upper right** *µ* = *ν* = 10^−2^ *h*^−1^. **Lower left** *µ* = *ν* = 7.10^−3^ *h*^−1^. **Lower right** *µ* = *ν* = 2.10^4^ *h*^−1^.

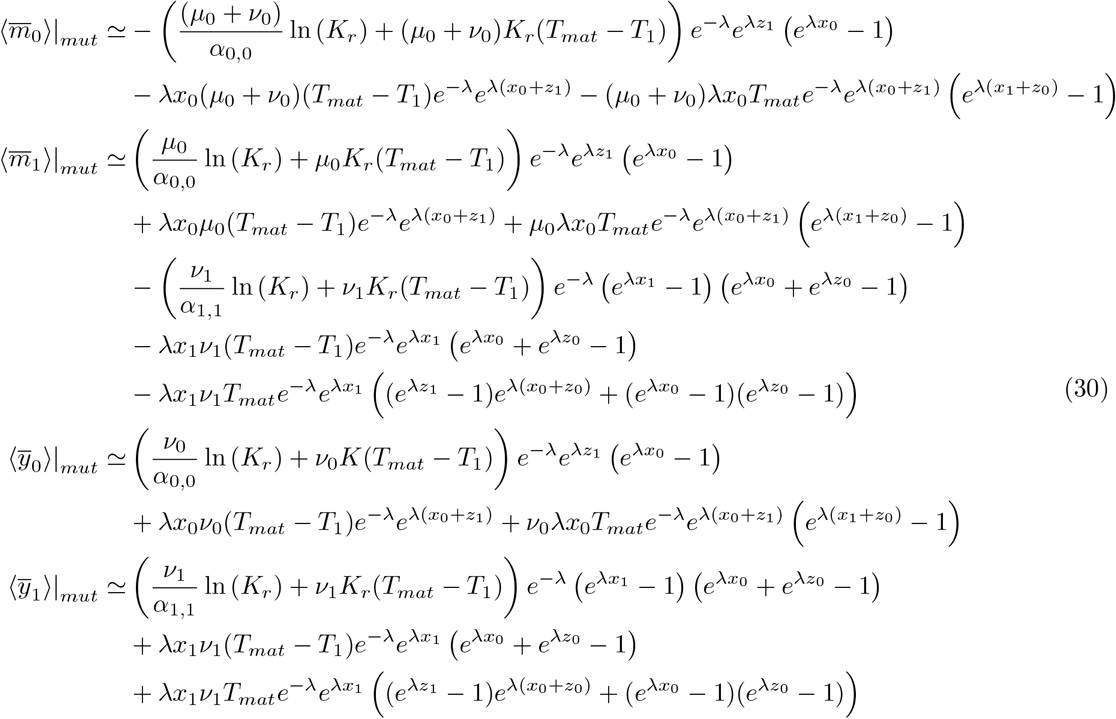

#### Pooling

Pooling consists in erasing the memory of individual compartments by putting their content together in a common pool. After pooling, the fraction of replicases depends on the repartition before maturation (compartmentalization step), then each compartment evolves according to the set of equations Eq.12. Therefore, the new fraction of replicases of type *k* is related the former one according to

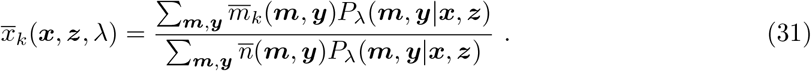

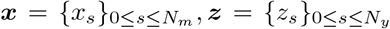 are the fraction of each mutant of replicases (where *N*_*m*_ is the number of mutant types for replicases) and parasites (where *N*_*y*_ is the number of mutant types for parasites), *λ* is the parameter of the Poisson distribution, and *P*_*λ*_(***m, y***|***x, z***) is the multinomial probability to have a vector ***m*** describing the numbers of each replicases and a vector ***y*** describing the numbers of each parasites starting with a pool with fractions ***x*** of replicases and fractions ***z*** of parasites and the average number of molecules per compartment is *λ*. 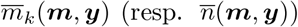 is the final number of replicases after maturation of type *k* (resp. final number of molecules after maturation) in the compartment given that it was initially filled with *n* molecules among which were ***m*** replicases and ***y*** parasites. We can set the time of maturation to be *T*, or fix the final number in each cell as a constant value 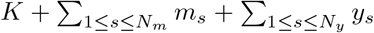. In the following we assume that *T* is large enough for both conditions to be equivalent.

**Figure S5.**
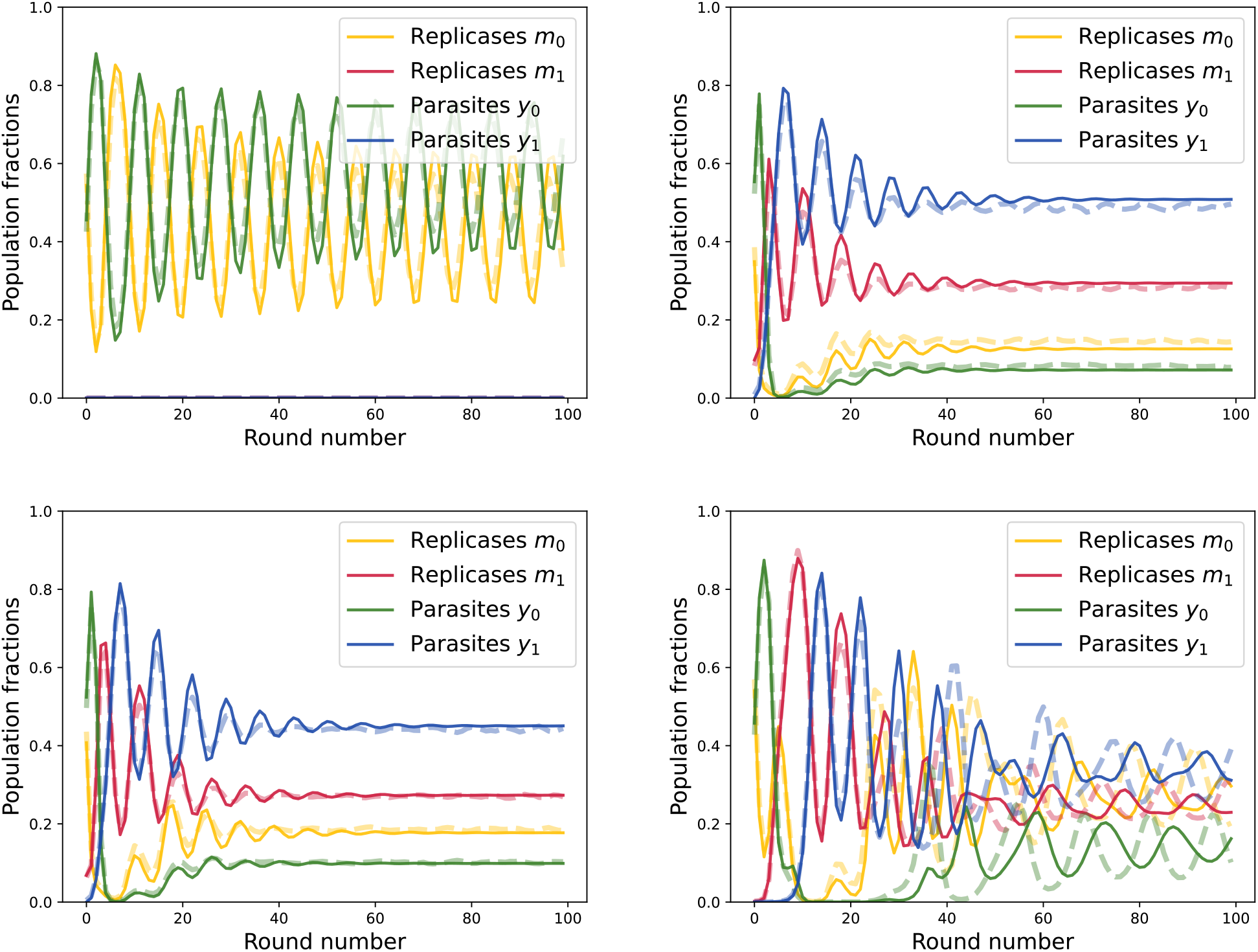
Theoretical prediction for model B (full) of the evolution of the system compared to Gillespie simulations (dotted) performed with 50000 compartments. We used for replication rates *α*_0,0_ = 5 × 10^−1^ *h*^−1^, *α*_1,1_ = 2 *h*^−1^, *γ*_0,0_ = 7 *h*^−1^, *γ*_1,1_ = 20 *h*^−1^. The carrying capacities are *K*_*p*_ = *K*_*r*_ = 30 and dilution factor *d* = 10. The different panels correspond to different mutation rates. **Upper left** *µ* = *ν* = 0 *h*^−1^. **Upper right** *µ* = *ν* = 10^−2^ *h*^−1^. **Lower left** *µ* = *ν* = 7.10^−3^ *h*^−1^. **Lower right** *µ* = *ν* = 2.10^4^ *h*^−1^.

#### Dilution

Dilution simply consists in dividing the total number of individuals by a dilution factor *d* to increase the total available volume. In experiments it also consists in refreshing the medium to allow for a new cycle. Therefore the average number of individuals per compartment is *λ/d*. Otherwise the updated fractions remain the same. If the dilution factor is larger than *λ*, the compartments will often be empty or contain 1 individual, thus favoring replicases. Therefore the new Poisson parameter is

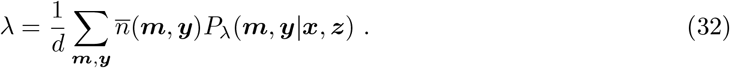

#### Mechanisms of oscillations

We typically observe oscillations (both experimentally and theoretically) between the fractions of the different species. Oscillations result from the variations of the average number of individuals per compartment. When *λ* is close to 1 replicases are favored because parasites tend to be isolated. On the contrary, when *λ* gets larger than 2 parasites are likely to be compartmentalized with a replicase, and the fraction of replicases starts decreasing. To estimate the period of the oscillations, we can thus consider two phases :

- A first phase (phase A) starting from *λ* ∼ 1 where the fraction of replicases is significant. In this case *λ* is likely to increase up to *λ* ∼ 2, when parasites start to invade the system.
- A second phase (phase B) starting from *λ* ∼ 2 where the fraction of replicases is close to 0. In this case *λ* decreases up to *λ* ∼ 1 and we back to the first phase.

To estimate the duration of the first phase, we first assume that the probability to have a replicase in a compartment is fixed equal to *p*_1_, such that *Kp*_1_ ≥ (*d* − 1) (in practice 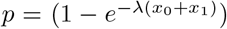 so this is wrong but we make this assumption to get an estimate). We can then write a recursive equation on *λ*_*t*_ where *t* is the cycle index

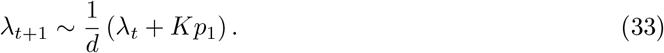

Starting from *λ*_0_ = 1, this phase will last *n*_1_ cycles with

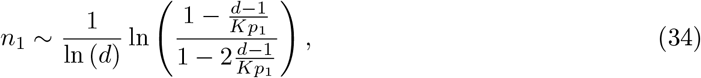

which is equal to ln((*K* − *d* − 1)*/*(*K* − 2(*d* − 1)))*/* ln(*d*) ∼ 0.24 when *p*_1_ ∼ 1 and diverges as *p*_1_ = 2(*d* − 1)*/K* = 0.6 for *d* = 10 and *K* = 30.

For the second phase, the recursion is similar, but the probability of having a replicase is now *p*_2_ ≪ 1 (in particular *Kp*_2_ *< d* − 1) and *λ* is initially equal to 2. The number of cycles required to get to *λ* ∼ 1 is thus

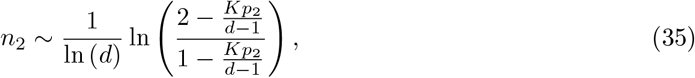

which is equal to ln(2)*/* ln(*d*) = 0.3 when *p*_2_ → 0, and diverges as *p*_2_ = (*d* − 1)*/K* = 0.3 for *d* = 10 and *K* = 30.

**Figure S6.**
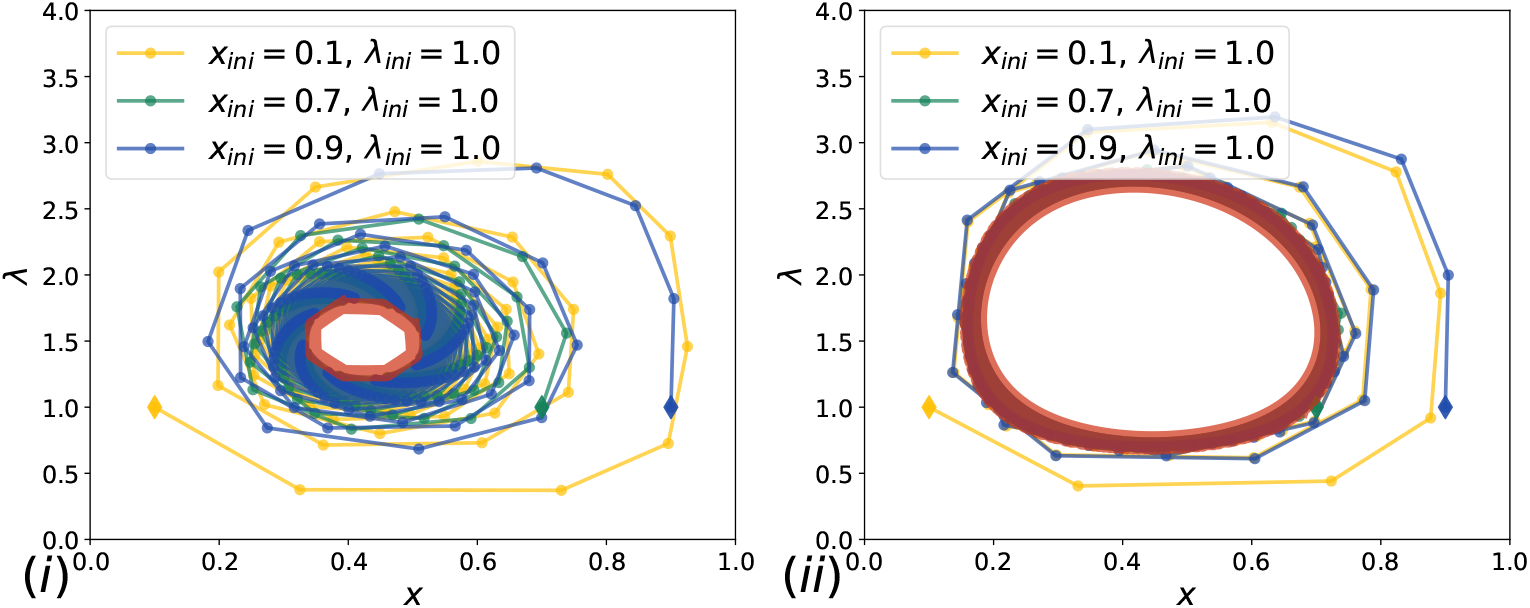
Oscillations between *λ* and *x*_*tot*_ with two species. The initial conditions are shown with diamonds. In red we represent the typical oscillation pattern towards which the system equilibrate, if *K/d* is large enough. For smaller *K/d* the system reaches a steady state. **(i)** *K* = 29, *d* = 10, **(ii)** *K* = 32, *d* = 10.

Therefore, the transition from phase B to phase A, and thus the period is set by the fraction of replicases *x*_0_ + *x*_1_. In particular, *λ* will spontaneously switch from *λ* ∼ 2 to *λ* ∼ 1 when

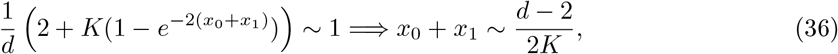

which is *x*_0_ + *x*_1_ ∼ 2*/*15 with *d* = 10 and *K* = 30. In practice we observe periods of a few cycles for all conditions. This results from the fact that once *λ* is larger than 2, parasites invade the system in a few cycles, then there is a transition to *λ* ∼ 1, then parasites are isolated and the fraction of replicases increases again for a few cycles until it starts again.

##### With two species

For two species, with one parasite (fraction 1 − *x*_*t*_ at time *t*) and one replicase (fraction *x*_*t*_ at time *t*), we have the following

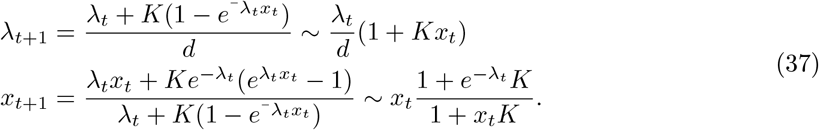

We plot a diagram of *λ* as a function of *x* in Fig. 6, where it is clear that there are oscillations between a state with large *K* and small *x*, and a state with small *K* and large *x*. Typically we see that a period corresponds to approximately 9 cycles. We also see that for various initial conditions, the system equilibrate towards a typical oscillation pattern, shown as a red area in Fig. 6. For smaller *K/d*, the system reaches a steady state. Indeed we can compute the equilibrium solution

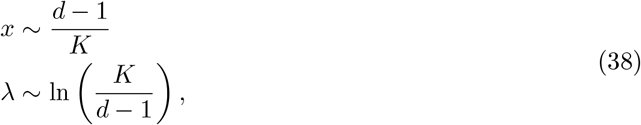

and introduce perturbations *x*_*t*_ = *x* + *ϵ*_*t*_, *λ*_*t*_ = *λ*(1 + *µ*_*t*_), then we find that stability is given by

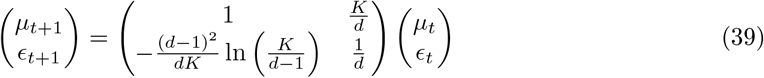

Therefore, the eigenvalues are

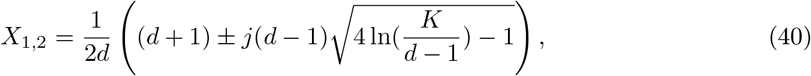

and their absolute values are

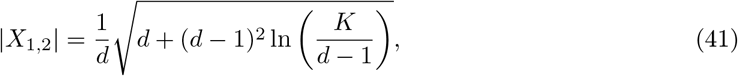

from which we deduce that the system becomes unstable for

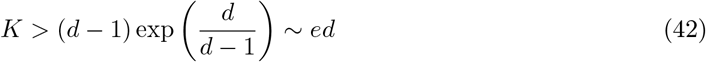

This is not exactly true because we linearized the equations to find these results.

##### With four species

In this case, we recover the same results in the limit where either wild-type or mutant dominate the system. However, when they coexist, the limit for instability is given by

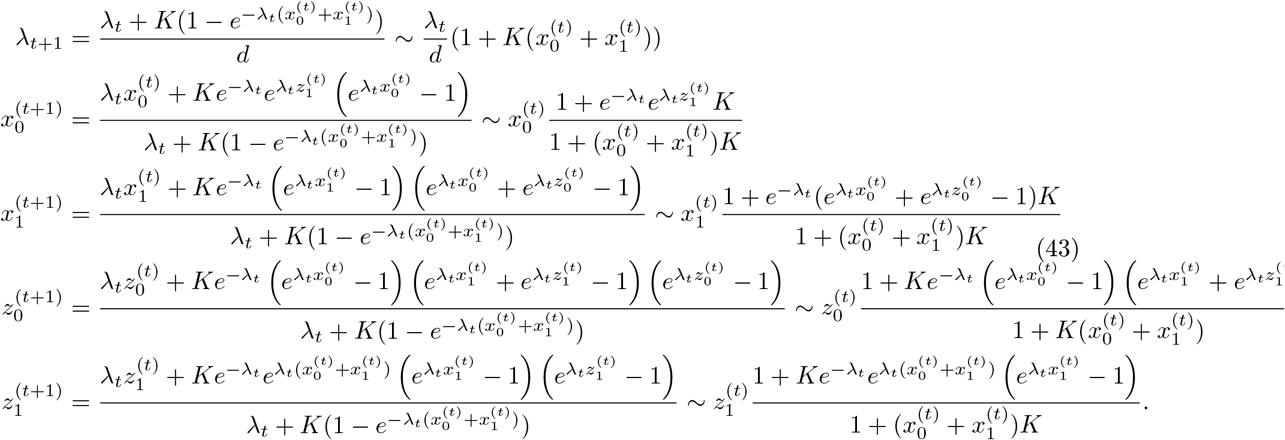

We observe in Fig. 7 that oscillation patterns are more complex with mutations. In addition, the threshold for instability is at larger values of *K/d*. We also observe that the system equilibrate at larger values of *λ*.

### Stirring

#### Modeling the stirring step

The experimental procedure of serial transfer is somewhat different from the dynamics we considered so far. The experimental protocol consists in:

**Figure S7.**
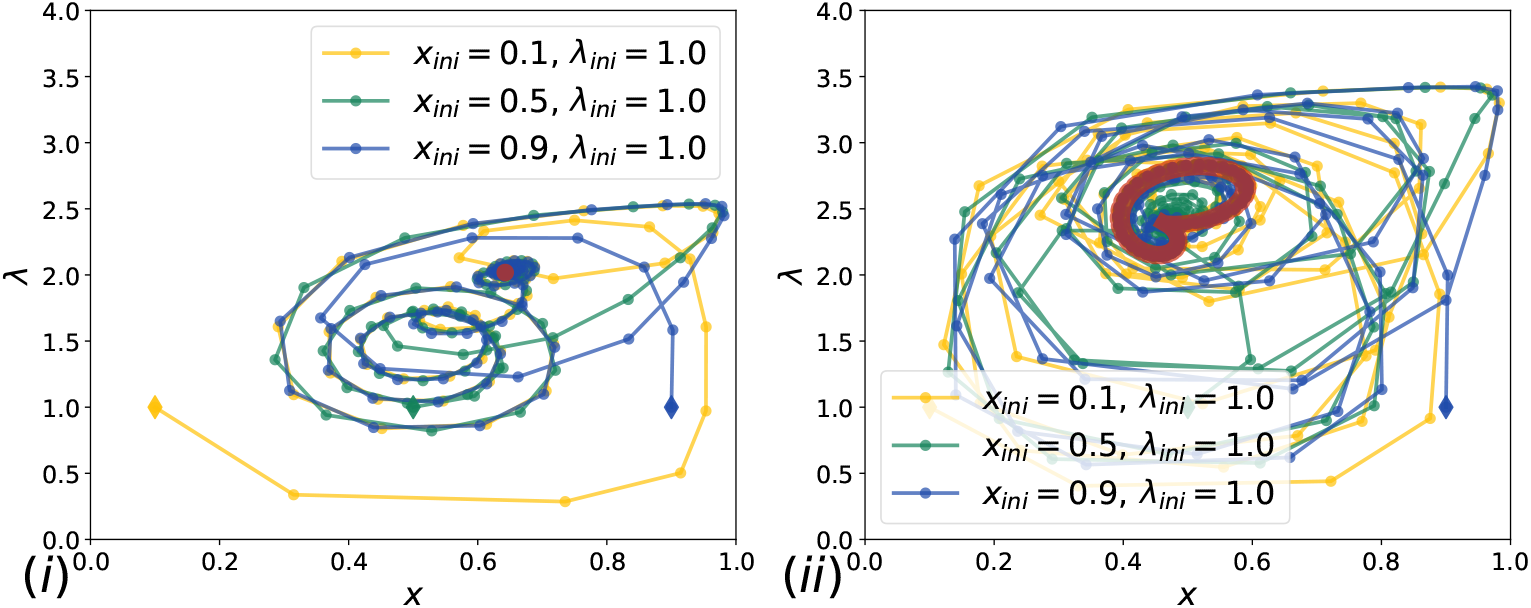
Oscillations between *λ* and *x*_*tot*_ with four species and within model B. The initial conditions are shown with diamonds. In red we represent the typical oscillation pattern towards which the system equilibrate, if *K/d* is large enough. For smaller *K/d* the system reaches a steady state. **(i)** *K* = 29, *d* = 10, **(ii)** *K* = 32, *d* = 10.

- Preparing compartments with different initial conditions
- Let compartments evolve until they reach the carrying capacity (maturation phase)
- Dilute (remove some mature compartments and add empty ones to increase the total available volume).
- Stir the solution to induce fusions and divisions of compartments.

In order to model this, we propose a dynamics, which incorporates a form of memory. Mathematically, we propose to do this by the following procedure:

- Starting from a distribution of initial conditions in the cells, we let them evolve (maturation phase)
- We randomly remove mature cells and replace them by empty cells (dilution)
- We update the composition of compartments according to an interpolation between their former compositions (memory effect) and the average composition of the whole population (mean field effect due to division/fusion events). This approach is mean field as we consider that every compartment is interacting with the average composition of the total population. Indeed, stirring induces homogenization of the compartments towards the average fractions in the whole population, but in reality the new compositions of every compartment will also be determined by its former composition as stirring is not strong enough to fully homogenize the system (some information is preserved). In particular we introduce a stirring parameter *s* ∈ [0, 1] such that the compartment *i* after maturation, 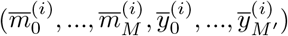, (with *M, M* ^′^ the number of mutants of replicases and parasites respectively) becomes 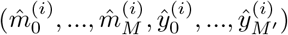. The total number of individuals in the cell 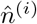 is drawn according to a Poisson distribution of parameter 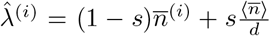 and the new numbers 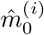 (similar for all other species) are drawn according to a multinomial distribution with parameters (in the limit on an infinite number of compartments)

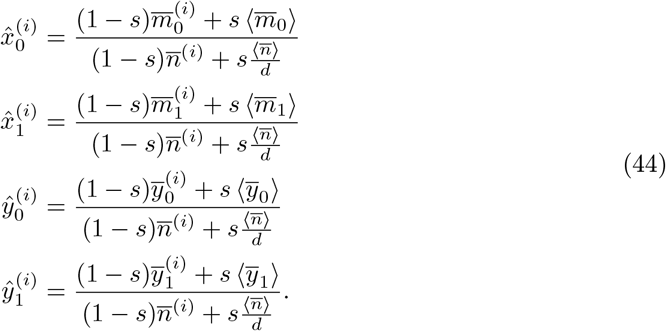

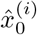 is the fraction of WT replicases in the compartment depending on the level of stirring. We keep the notations 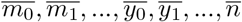 for the numbers of each species in one compartment after maturation and before dilution, *n*^(*i*)^ is the number of individuals in compartment *i*. If *s* = 1, we recover the dynamics discussed in the previous sections and the distribution is homogeneous (no memory). Whenever *s <* 1, we keep track of inhomogeneities (memory effect). This imposes to treat each compartment separately in simulations. We also see that *s* = *d/*(*d* + 1) is an important value as it corresponds to the stirring where the influence of the mean field effect has the same weight as the memory effect. For *s > d/*(*d* + 1) we expect to recover the perfectly homogeneous case.

We find that the total fraction of replicases surviving in the system depends on the level of stirring. In particular, it increases with the stirring parameter *s* because higher stirring tends to isolate parasites, unless it leads to extinction (see next Section). We recover this trend both for 2 species (one parasite and one replicases) in Fig. 9, as well as for 4 species (two parasites and two replicases) in the main text.

#### Assessing the effect of stirring for two types of individuals

To get insights on the stirring process, we study experimentally stirring for a population with two types of individuals. Typically one population is indicated with green fluorophores and the other with red fluorophores. In practice we assume that during fusion division events, compartments exchange a fraction of their content. We assume that they exchange packets of molecules proportional to the average fluorescence intensity in the population and call *G*_*k*_ the number of packets of green individuals in compartment *k* before stirring, and 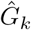 after stirring, We also call *R*_*k*_ the number of red individuals in compartment *k* before stirring, and 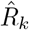 after stirring. We call *N*_*k*_ = *G*_*k*_ + *R*_*k*_ (resp. 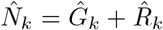) the total number of packets of individuals in compartment *k* before stirring (resp. after stirring). We model packets of molecules instead of individual molecules because we observed that even with no manually induced stirring, we get distribution of droplets contents where the probability of having half green and half red is non zero (starting from a perfectly bimodal distribution, that is either fully green or fully red). This suggests that even with a few fusion/division events, some droplets will have exchange half of their initial content. In practice, we obtain the distribution of numbers of packets by rescaling the distributions of fluorescence intensities by the average total fluorescence, we justify this assumption in Fig. 10A.

We have the following rules to model exchanges of contents

**Figure S8.**
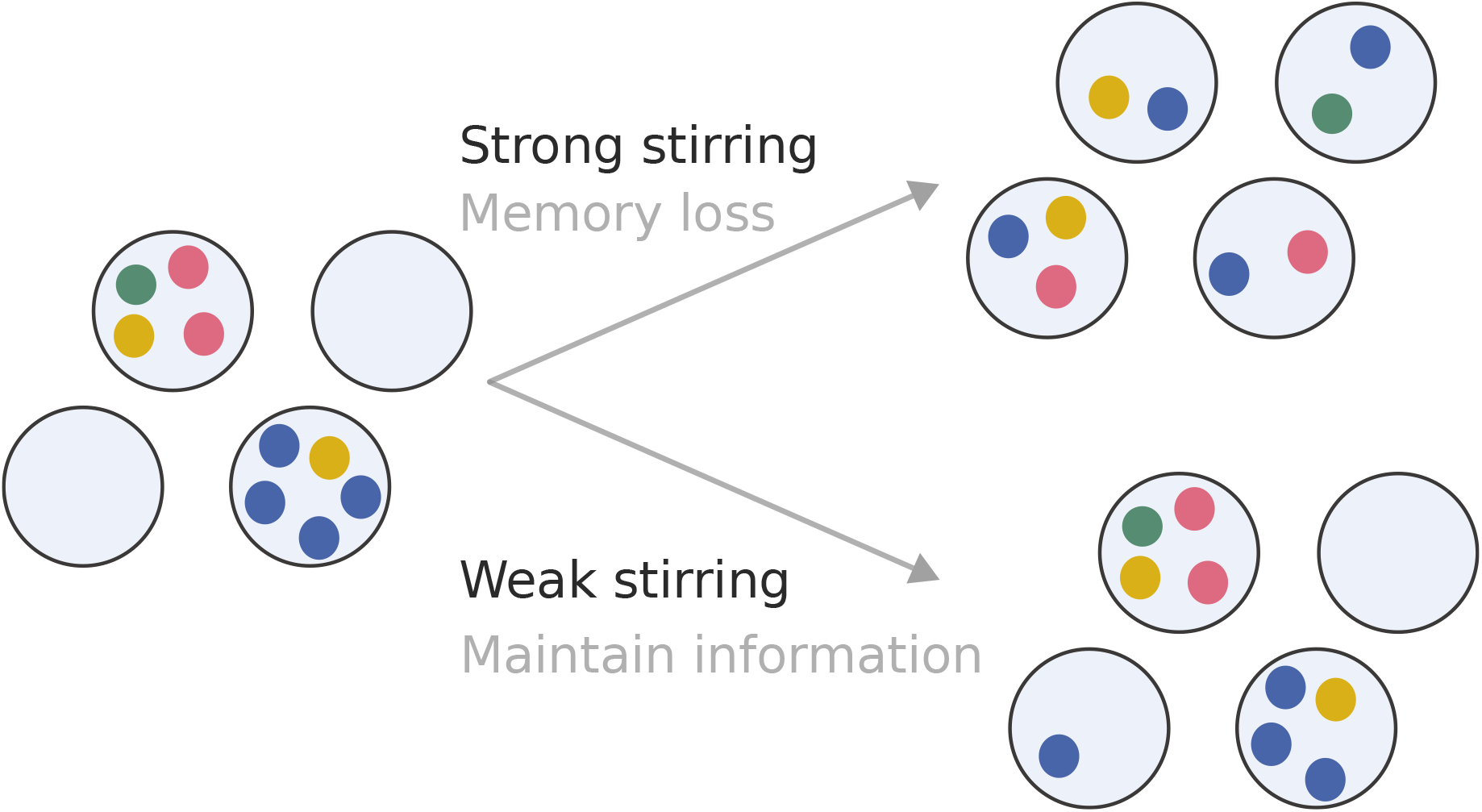
Memory for stirring. With strong stirring, the compartments are homogenized and partially erase their compositional memories. With weak stirring, compartments inherits from their former content after each stirring step and retain som compositional memory.

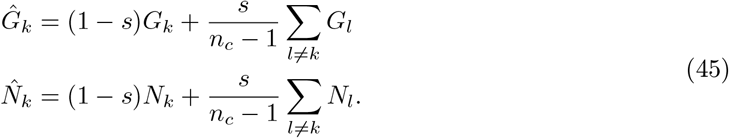

In this case, the cumulative probability distribution of the updated fraction after mixing is the probability that the updated fraction in a compartment *k*, 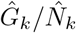 is lower than a given value *x*

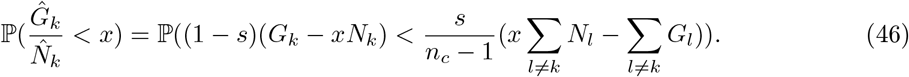

theorem. Accounting for the fact that *G*_*l*_ and *N*_*l*_ are correlated we obtain that

We approximate the sums ∑_l≠*k*_ *G*_*l*_ and ∑_*l* ≠*k*_ *N*_*l*_ as normally distributed using the central limit

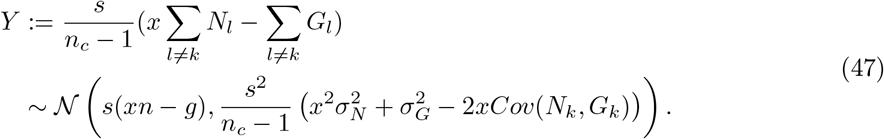

Now, we can write the cumulative distribution probability, using the law of total probability

**Figure S9.**
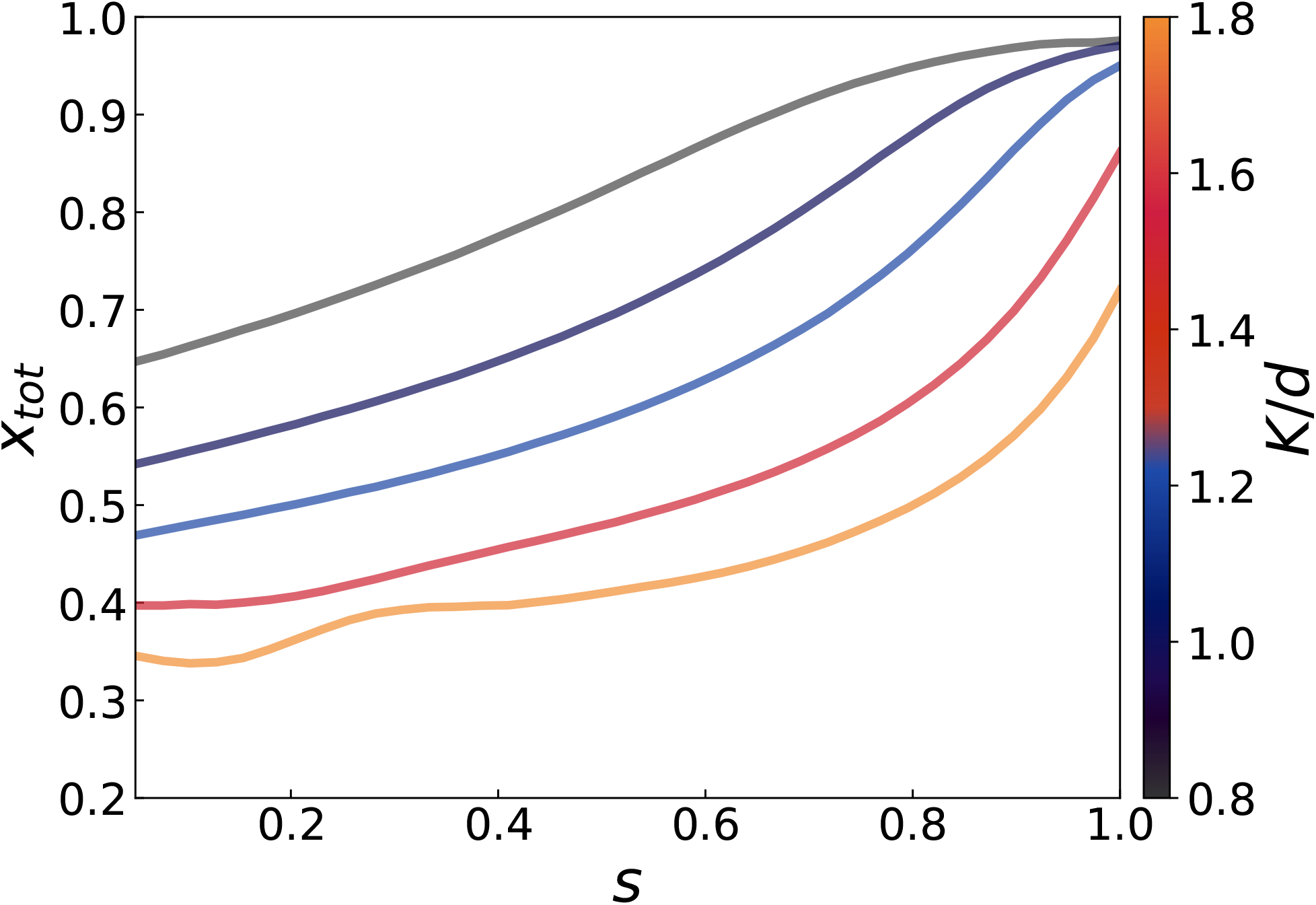
Total fraction of replicases *x*_*tot*_ depending on the stirring parameter *s* with two species (one replicase and one parasite). The colour bar indicates the value of the ratio carrying capacity to dilution *K/d*. The lower it i, the less compartments should be populated.

**Figure S10.**
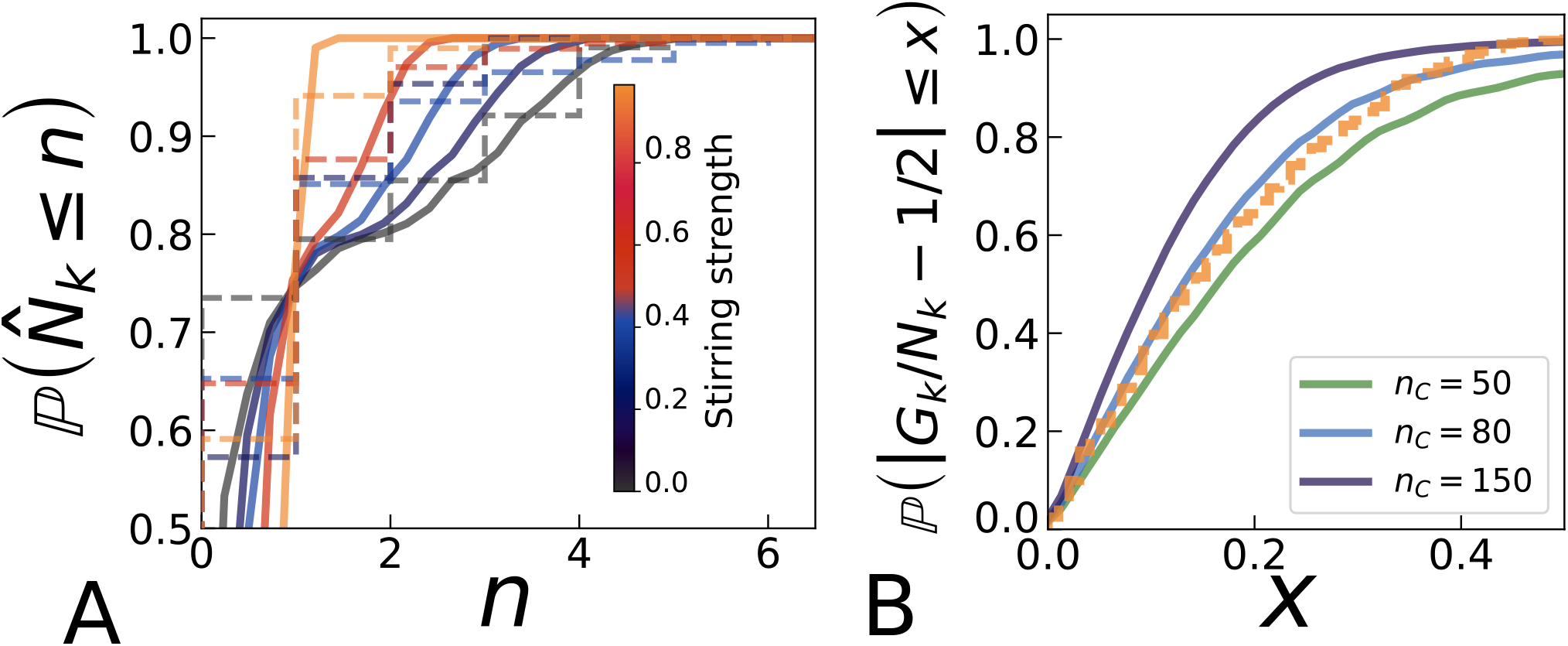
**A** Experimental distributions of 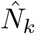 (total fluorescence rescaled by the average total fluorescence per compartment) and theoretical predictions. We conclude that taking packets of molecules corresponding to the average total fluorescence per compartment seems to describe well the exchanges between compartments. In dotted lines we show the experimental cumulative probability distributions (discretized because it respresents the number of packets of molecules emitting a fluorescence equal to the average total fluorescence per compartment). In solid lines we show expectations from the theory. **B** Comparison of the experimental distribution with maximal stirring (30 krpm) and theory starting from a perfectly bimodal distribution for *s* → 1, used to calibrate the value *n*_*C*_ = 80.

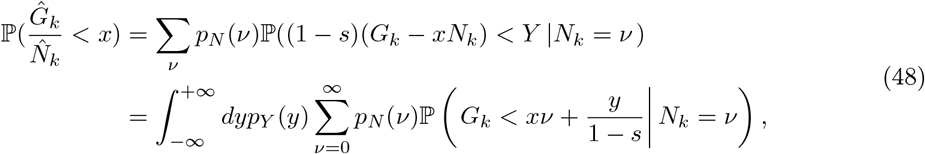

with

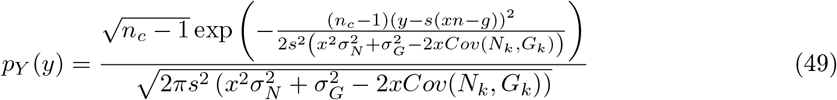

where *p*_*N*_ is the distribution probability of *N*_*k*_. With this method we obtain Fig.4 of the main text, which is in good agreement with the data. This suggests strong values for the stirring s, of about *s* = 0.9. If we start from a perfectly bimodal distribution for the conditional probability *p*_*G,ini*_(*γ*|*N*_*k*_ = *ν*) = (*δ*_*γ*,0_ + *δ*_*γ,ν*_)*/*2, and an initial experimental distribution for *N* with ⟨*N*_*k*_⟩ = *n* and standard deviation *σ*_*N*_, we get ⟨*G*_*k*_⟩ = *n/*2 and 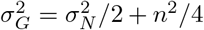. In addition 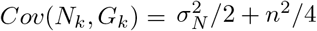. We get

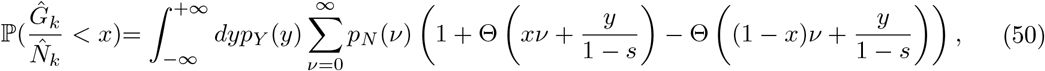

To study the strength of stirring, we quantify the “bimodality” of the distribution by studying the distribution of 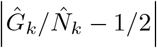. If the distribution is centered around 0, it means that the distribution is rather monomodal, while if it’s centered close to 1*/*2, it’s rather bimodal. This way we remove the bias due to the fact that the distribution of 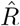 is not exactly the same as that of *Ĝ*. To do this we must introduce 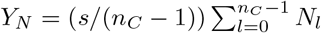 and 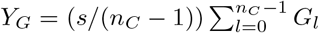. We assume that both follow normal distributions 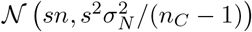 and 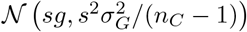, and *Cov*(*Y*_*N*_, *Y*_*G*_) = *s*^2^*Cov*(*N*_*k*_, *G*_*k*_)*/*(*n*_*C*_ − 1). Therefore

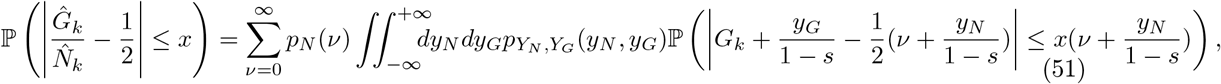

where we take *Y*_*N*_, *Y*_*G*_ to be bivariate. Therefore, with 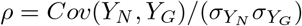 and

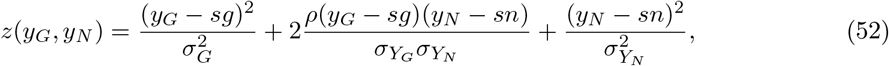

we get

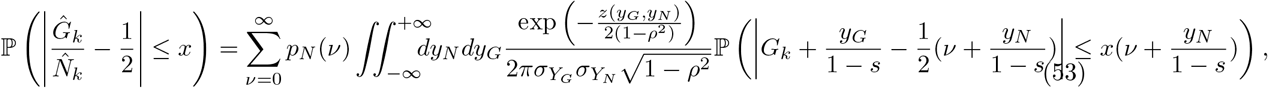

To estimate the value of *n*_*C*_ from data, we use the experimental system with strongest stirring (after one round with active stirring at 30 krpm) and assume this corresponds to *s* = 0.95 between an initial bimodal distribution and this final distribution. In the case *s* → 1, we have

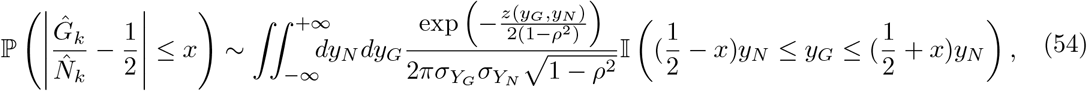

which is thus independent of the distribution of *G*_*k*_*/N*_*k*_. this means that for a given distribution of *N*_*k*_, the transformed distribution with *s* = 1 is a fixed point of the transformation. From this equation, assuming that this should correspond to data for one round of active stirring with 30 krpm, we estimate *n*_*C*_ ∼ 80 for strong stirring as can be seen in Fig. 10 B.

in Fig.4 of the main text, we see the result of the dye experiment and the comparison with the theory. We can compute the cumulative probability distribution 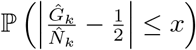, which should be 0 everywhere and then 1 at *x* = 0.5 for a perfectly bimodal distribution in 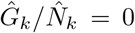 and 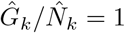, while it should be equal to 1 everywhere for a perfectly monomodal distribution centered in 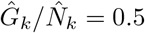. We see that as stirring strength increase, the distribution after stirring is closer to a monomodal one (meaning loss of memory of individual compartments).

From this description, starting from a bimodal distribution, we deduce *s* ∈ [0 : 0.2] for 2 krpm, *s* ∈ [0.2 : 0.4] for 5.2 krpm, *s* ∈ [0.5 : 0.6] for 16 krpm, and *s* ∈ [0.9 : 1] for 30 krpm (illustrated in Fig. 4B of the main text and used values in these intervals for Fig. 3A of the main text). We can also evaluate how robust are these results by comparing the experimental distributions after round 2 to the theoretical distributions starting from the experimental ones before round 2, with the inferred values of *s* for each stirring strengths. This is shown in Fig. 11, where we see that theoretical estimations are slightly closer to monomodality than the experimental ones. This might come from the fact that the size of packets is slightly larger in this case. However we conclude that our estimations of *s* mostly encapsulate the effect of stirring for 2 krpm, 5.2 krpm, 16 krpm and 30 krpm, thus validating the correspondencies we established above between stirring strengths and stirring parameters.

#### Stirring and extinction

In some cases, when *K/d* is smaller than 1, meaning than dilution tends to washout the system, we observe extinction of the whole population for high enough values of stirring. Lower stirring increases content heterogeneity and may thus lead to having statistically more crowded compartments, allowing the population to survive. The updated number of individuals in compartment *i* is drawn according to a Poisson distribution with parameter

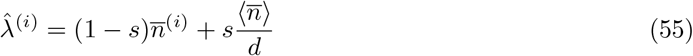

where 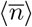 is the average content (number of individuals) after maturation and before dilution and 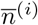 is the number of individuals in compartment *i* after maturation if it has not been removed during dilution, and 0 otherwise. This means that 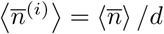. This leads to

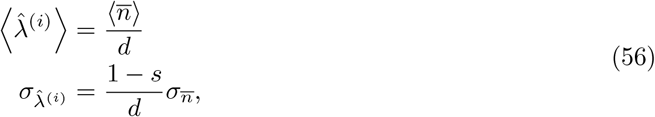

and thus we have a value of *s* that allows some compartments to contain more than 1 individual

**Figure S11.**
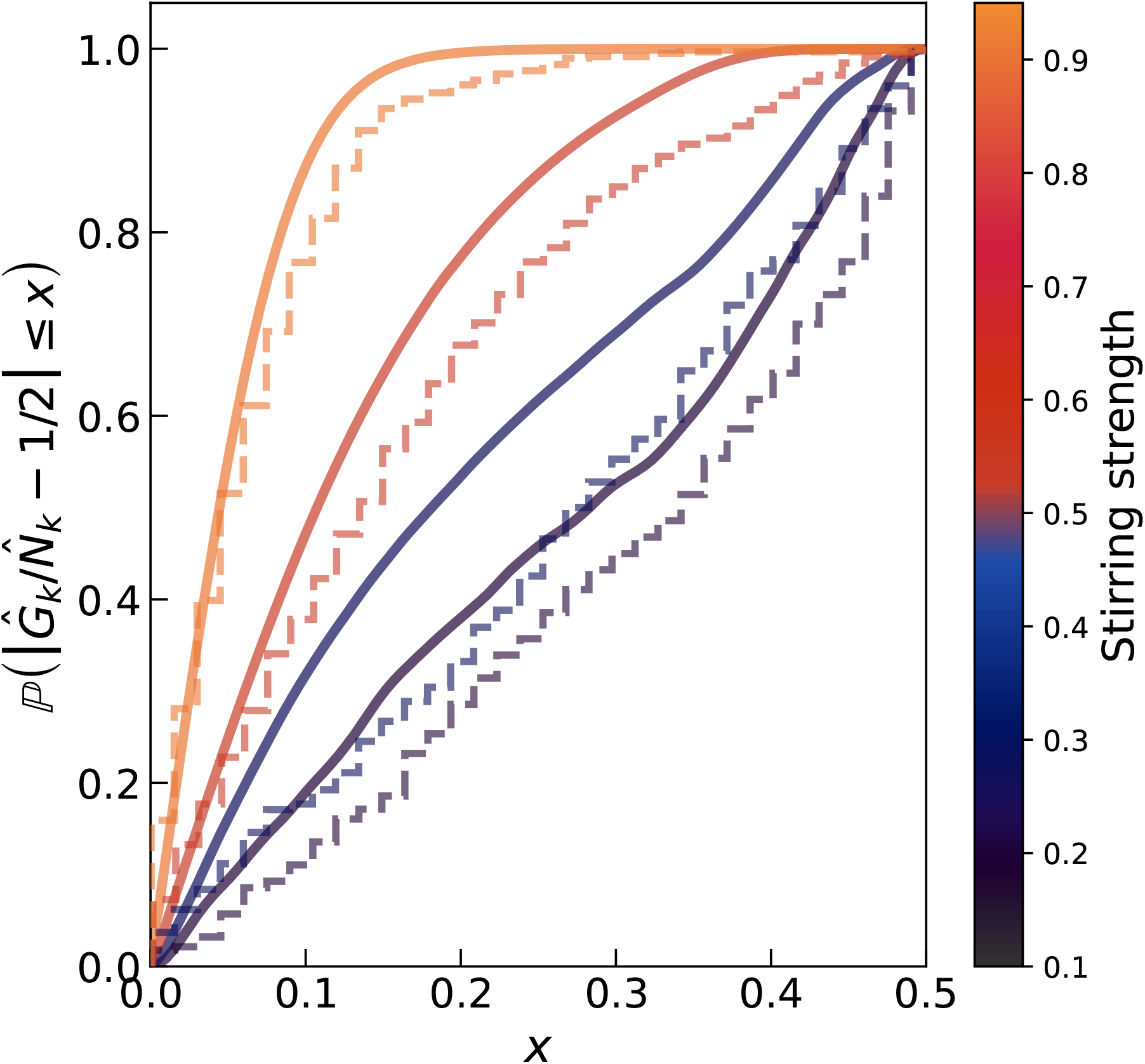
Assessment of the robustness of estimations of *s*. Starting from experimental distributions before round 2, we apply the theoretical transformation with *s* inferred from the first round (starting from a perfectly bimodal distribution of 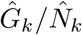) and compare it to the experimental distribution after round 2 for each stirring strengths.

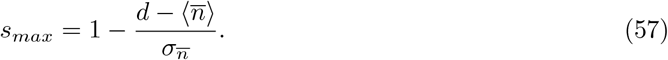

We can compute

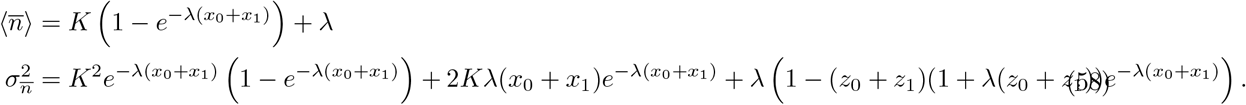

In terms of scaling, we thus get that

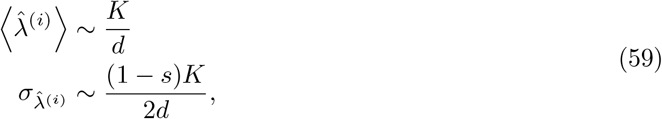

which sets a value of *s* for which we can hope to have a more than 1 individual, and thus the possibility to survive, in some compartments

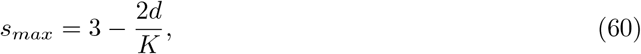

which explains qualitatively that the larger *d/K* is, the smaller *s* is required to observe extinction. In practice this also depends on the number of cycles and total number of compartments. Indeed for a finite number of compartments *n*_*C*_, the probability that at least one compartment has more than 1 individual is

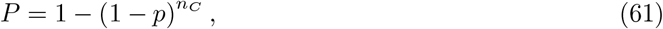

where *p* is the probability that a compartment contains more than 1 individual, which itself decreases with *s* (if *d > K*) as seen above, which can be approximated by 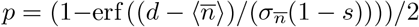, leading to

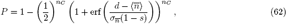

From which it is clear that it decreases with dilution *d* and increases with the stirring parameter *s* and the number of compartments *n*_*C*_. In addition, the number of cycles intervenes in the fact that *λ*^(*i*)^ successively decreases with time (it thus affects 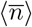 and 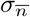, typically decreasing *P*). Indeed, for small *λ*^(*i*)^

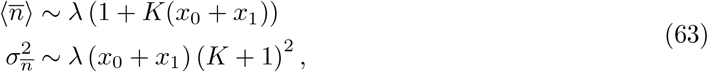

making *P* an increasing function of *λ* in this limit. This explains why heterogeneity favors survival, and the mechanism is sketched in Fig. 12.

**Figure S12.**
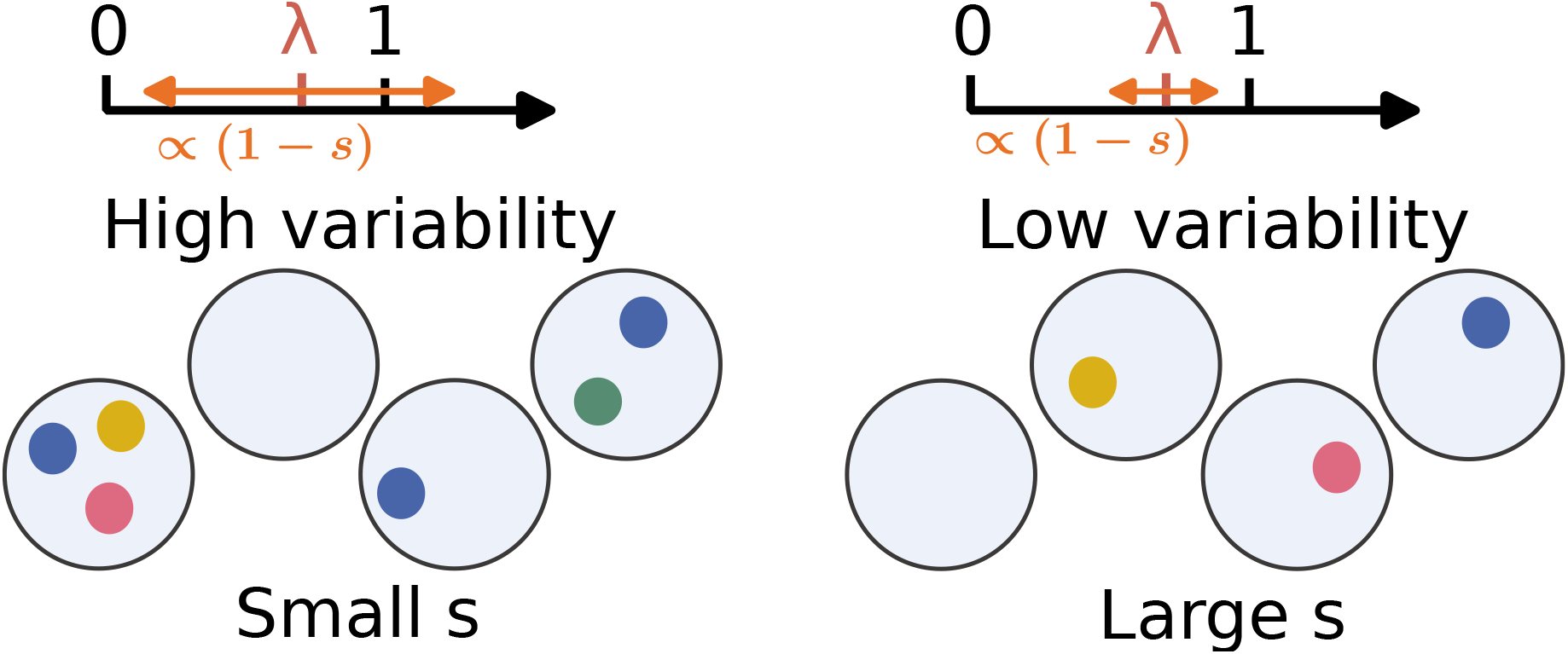
Mechanism for a small number of individuals and high stirring parameter *s*. When *s* decreases, the variability of the content of compartments allows for more populated compartments after dilution and stirring, and prevents from extinction. At large values of s, for *λ* ∼ 1, individuals are more likely isolated in one compartment. Isolated parasites cannot replicate, thus this limit typically favours replicases.

**Table S2.** Replication rates from experiments.

|  |  |  |
| --- | --- | --- |
| $\alpha_{0,0}$ | 2.6 | $h^{-1}$ |
| $\alpha_{0,1}$ | 0.0 | $h^{-1}$ |
| $\alpha_{1,0}$ | 1.4 | $h^{-1}$ |
| $\alpha_{1,1}$ | 2.8 | $h^{-1}$ |
| $\gamma_{0,0}$ | 5.7 | $h^{-1}$ |
| $\gamma_{0,1}$ | 3.0 | $h^{-1}$ |
| $\gamma_{1,0}$ | 0.0 | $h^{-1}$ |
| $\gamma_{1,1}$ | 3.0 | $h^{-1}$ |

#### Comparison to experiments

We compare the deterministic approach to experiments involving 4 species (see Fig.1D of the main text). We perform the following mapping with the deterministic model:

- *HL*1 ↦ Wild-type replicases *m*_0_.
- *HL*2 ↦ Mutant replicases *m*_1_.
- *PL*2 ↦ Specialist wild-type parasites *y*_0_.
- *PL*3 ↦ Generalist mutant parasites *y*_1_.

We use the replication rates in Table 2. We also use the experimental dilution factor *d* = 5, as well as mutation rates of *µ*_0_ = 1.3 × 10^−4^ *h*^−1^, *ν*_0_ = *ν*_1_ = 6.5 × 10^−5^ *h*^−1^, that are enough to ensure the progressive appearance of mutants in the system. We also fix *K* = 41.

Additionally, *HL*1 mutates to *HL*2, *HL*2 mutates to *PL*2 and *PL*2 mutates to *PL*3. *PL*3 is a generalist parasite, and *HL*1 is a generalist replicase in the sense that *HL*1 can be replicated by *HL*2. We assume the following deterministic rules:

- For a compartment with *HL*1 or *HL*2 only, they replicate until they reach the carrying capacity.
- For a compartment with *HL*1 and *PL*2 only, *HL*1 grows up to *K*_*r*_.
- For a compartment with both *HL*1 and *HL*2, but no parasites, *HL*1 will be the main species as they are generalist. However we assume that *HL*1 grows up to *r*_1_*K*_*r*_ and *HL*2 up to (1 − *r*_1_)*K*_*r*_, with *r*_1_ *>* 1*/*2.
- For a compartment with *HL*2 and *PL*2, *PL*2 grows up to the carrying capacity *K*_*p*_, regardless on the presence of other species.
- For a compartment with *HL*1 and *PL*3 only, *HL*1 grows up to *r*_2_*K*_*r*_ and *PL*3 up to (1−*r*_2_)*K*_*p*_.
- For a compartment with *HL*2 and *PL*3 only, *HL*2 grows up to *r*_3_*K*_*r*_ and *PL*3 up to (1−*r*_3_)*K*_*p*_.
- For a compartment with *HL*1, *HL*2 and *PL*3, *HL*1 grows up to *r*_1_*r*_4_*K*_*r*_, *HL*2 grows up to (1 − *r*_1_)*r*_4_*K*_*r*_ and *PL*3 up to (1 − *r*_4_)*K*_*p*_.

Those rules yield:

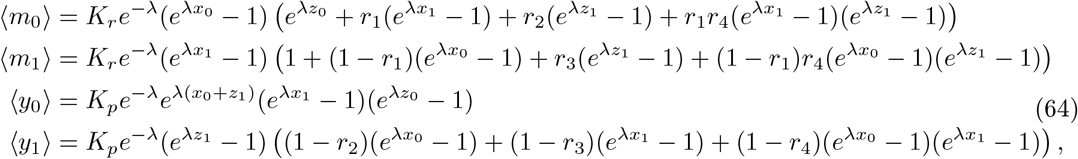

and the average numbers of mutants are

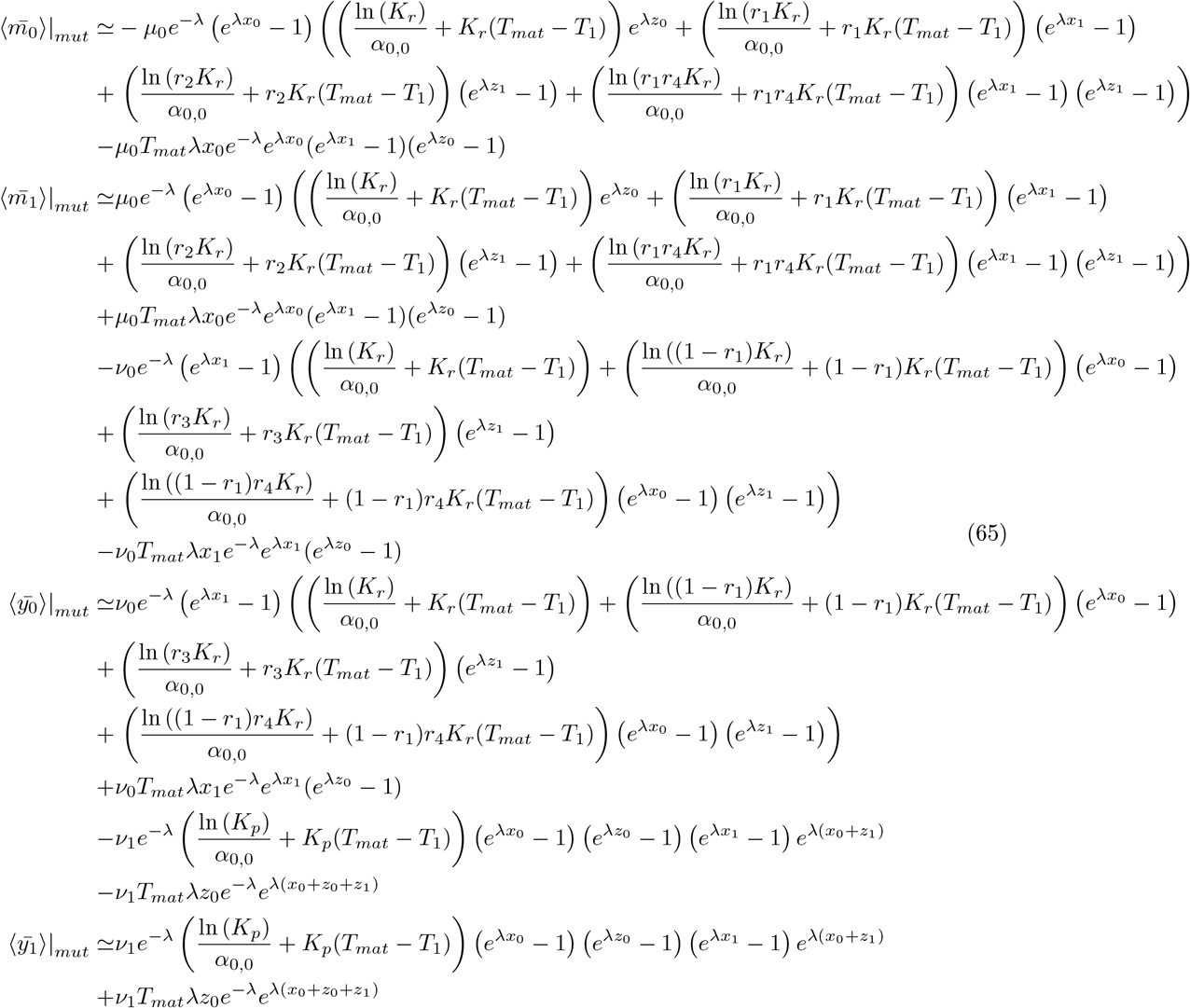

From such calculations, we obtain the evolution shown in Fig.1D of the main text and fit it with experimental data, where the fitting parameters are *r*_1_, *r*_2_, *r*_3_, *r*_4_. We find the values

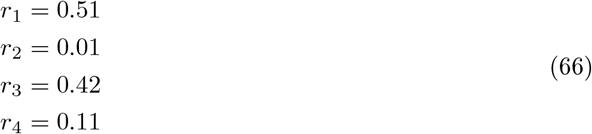

We use different values for *s*, the strength of stirring, and observe that the oscillations are slightly modified.

### Additional experimental data and materials

**Figure S13.**
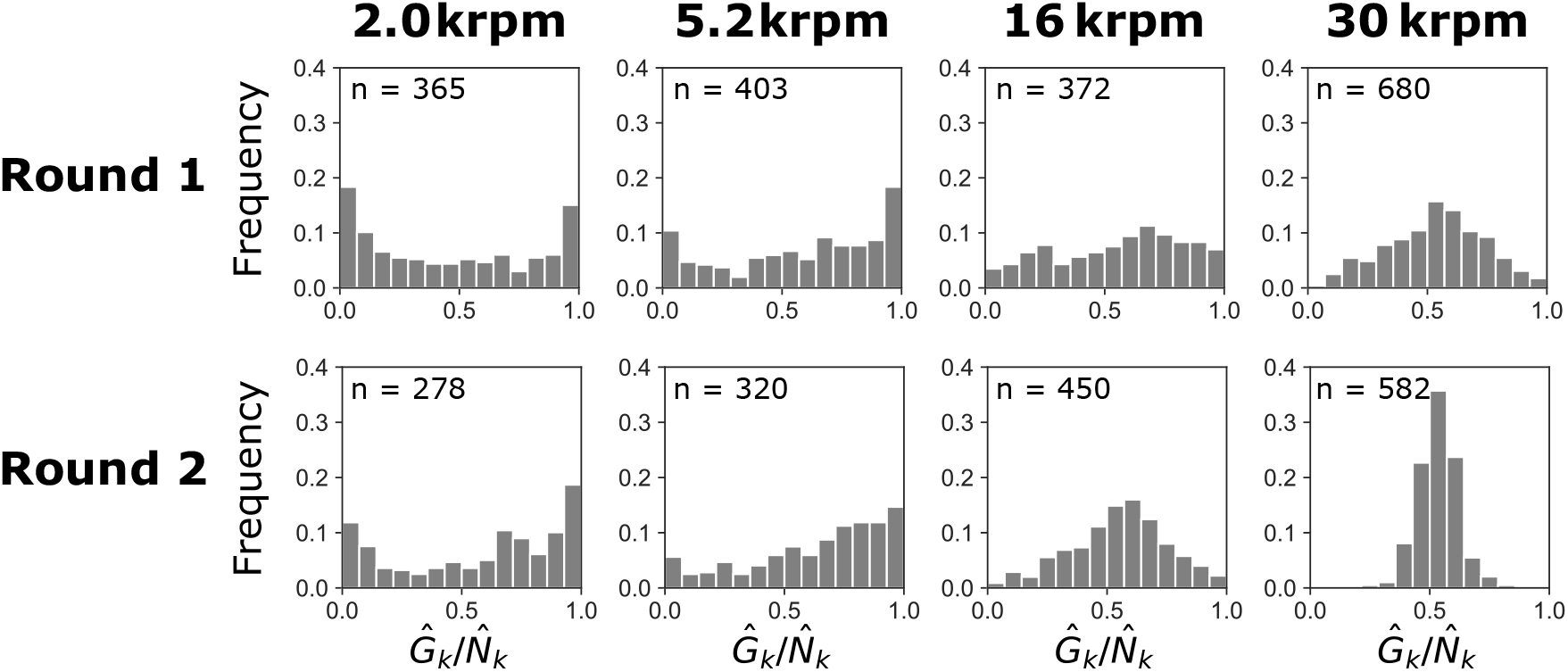
Distribution of fluorescent intensities over two rounds of the transfer experiment. The ratio of FITC fluorescence 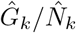 for each droplet containing FITC- or TRITC-labeled dextran was obtained by image analysis. Data for each condition were collected from three independent experiments. The number of analyzed droplets is indicated in each panel.

**Figure S14.**
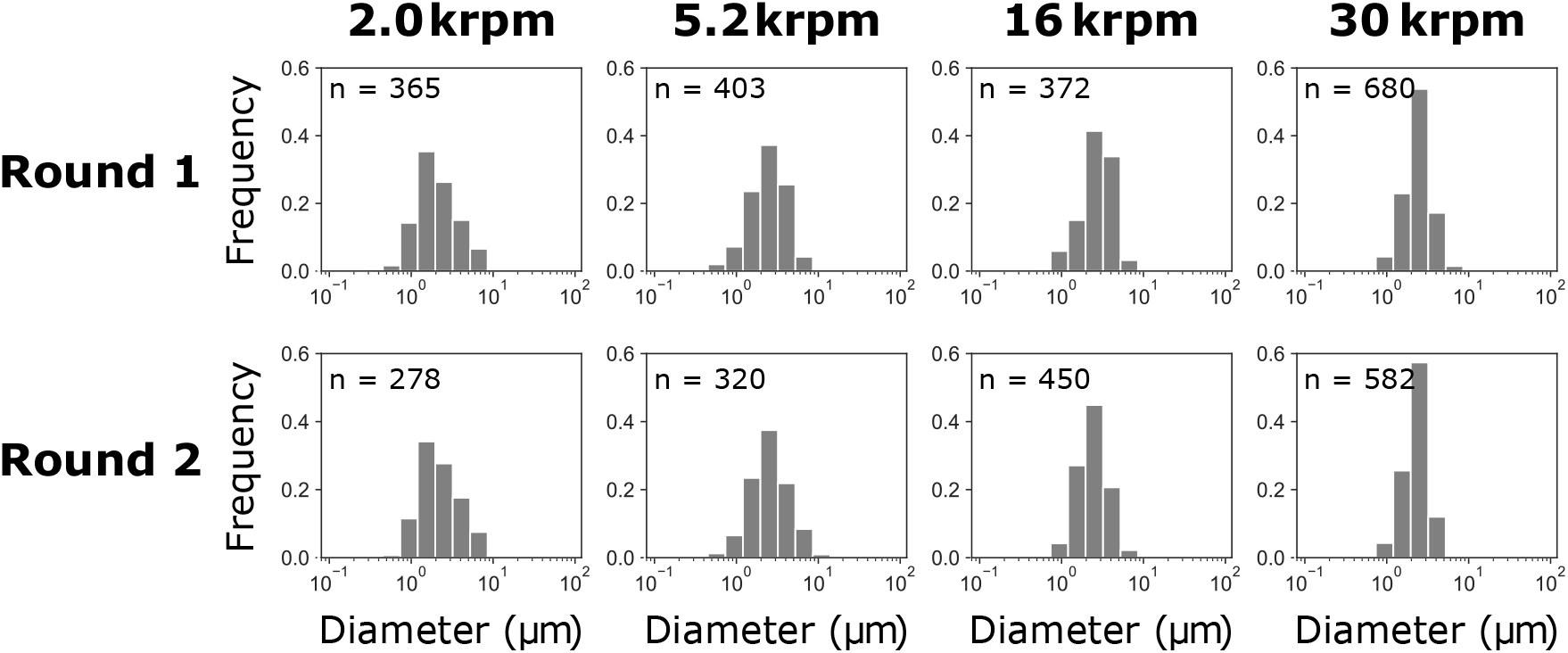
Size distribution of droplets over two rounds of the transfer experiment with fluorescent dyes. The diameters of individual droplets containing FITC- or TRITC-labeled dextran were obtained by image analysis. Data for each condition were collected from three independent experiments. The number of analyzed droplets is indicated in each panel.

**Table S3.**
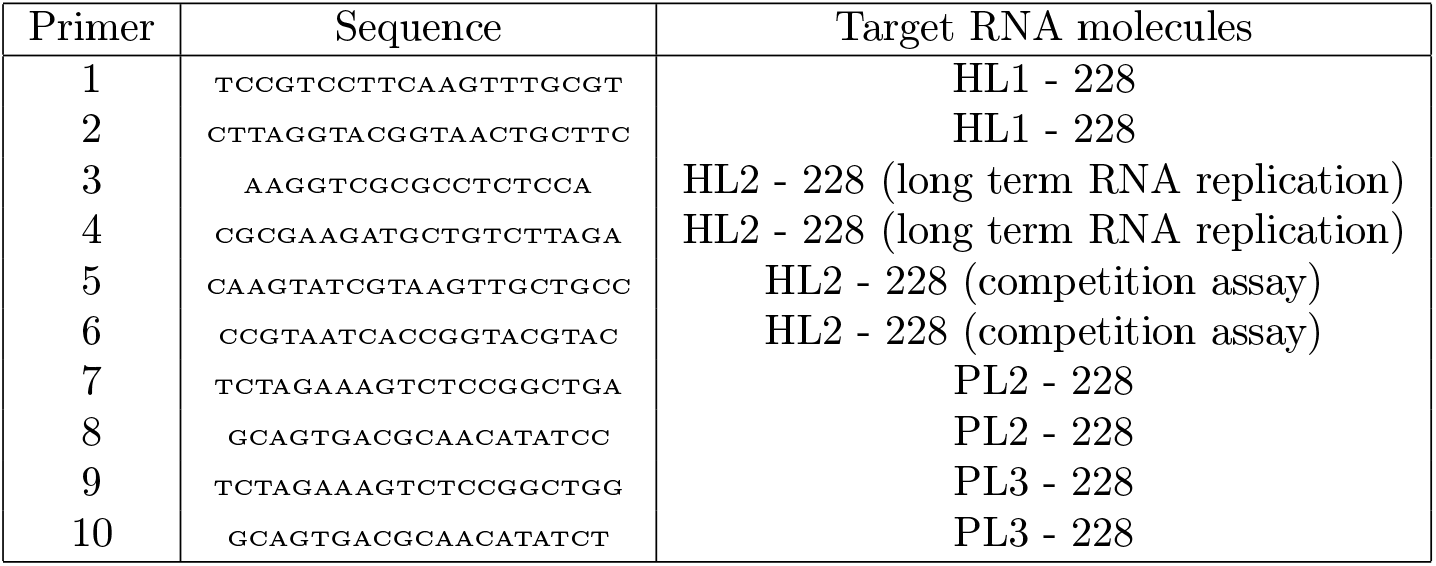
List of sequence-specific primers for quantitative RT-PCR (from 5’ end to 3’ end).

**Table S4.** Relative RNA concentrations estimated by RT-qPCR using different primer sets.

| Primers | HL1-228 | HL2-228 | PL2-228 | PL3-228 |
| --- | --- | --- | --- | --- |
| 1 & 2 | 1 | $2 \times 10^{-7}$ ($2 \times 10^{-8}$) | $2 \times 10^{-5}$ ($5 \times 10^{-6}$) | $1 \times 10^{-7}$ ($7 \times 10^{-8}$) |
| 3 & 4 | $9 \times 10^{-5}$ ($1 \times 10^{-5}$) | 1 | $2 \times 10^{-5}$ ($4 \times 10^{-6}$) | $2 \times 10^{-5}$ ($2 \times 10^{-6}$) |
| 5 & 6 | NA | 1 | $3 \times 10^{-5}$ ($6 \times 10^{-6}$) | NA |
| 7 & 8 | $5 \times 10^{-6}$ ($6 \times 10^{-7}$) | $6 \times 10^{-6}$ ($2 \times 10^{-6}$) | 1 | $4 \times 10^{-6}$ ($1 \times 10^{-6}$) |
| 9 & 10 | $6 \times 10^{-8}$ ($3 \times 10^{-8}$) | ND | $3 \times 10^{-4}$ ($2 \times 10^{-4}$) | 1 |

Values in parentheses indicate standard deviations (n=3). NA, not analyzed; ND, not detected (below the detection limit).

### Coexistence diagrams

#### Perfectly homogeneous limit

##### Theoretical coexistence diagram (*T, d*) (model A)

Using the deterministic approach, we can simulate the evolution of the system under transient compartmentalization and obtain a coexistence diagram depending on the dilution factor *d* and the maturation time *T*_*mat*_ in Fig. 15. In some phases, the system is out of equilibrium and oscillating as seen in Fig. 4, therefore we average the value of the different ratio over several repetitions and compare the average of the oscillating populations after enough repetitions to reach a limit cycle in Fig. 15. We observe that the appropriate timescale to study the system is 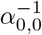. For high dilution factors, the compartments will initially have 0 or 1 individuals most likely, and the dominating replicase will depend on the maturation time (if maturation is too short, wild-type replicases will dominate). However, parasites can only replicate with replicases and mutant replicases dominate over wild-type replicases. Thus, in this limit, only mutant replicases survive. At large maturation times, the influence of mutations is larger, and we observe a decreasing trend with the maturation time in the fractions of wild-type populations. We also study the different states of the system in Fig. 16, highlighting the phases where some species are extinct, steady or oscillating. At low dilution factors, populations are oscillating more strongly as parasites are favored in the system (they will most likely be compartmentalized with replicases), and will periodically take over the replicases populations. But they will be diluted whenever there are no longer enough replicases to ensure parasite replication. At strong dilutions, wild-type populations go extinct and parasites are more likely isolated from replicases. We obtain steady coexistence for intermediate dilution factors, after a transient phase with damped oscillations. Steady states are more easily reached for large maturation times where enough mutants can appear to take over the system.

##### Theoretical coexistence diagram (*T, d*) (model B)

From the deterministic equation, and adding the estimated numbers of mutations for model B, we obtain the evolution of the system and compare to Gillespie simulations in Fig. 5. We build a coexistence diagram depending on the maturation time and dilution factor as done in the previous case. We observe that the coexistence diagrams Fig. 17 and Fig. 18 have common points with that of model A. The fact that the appearance of mutant parasites depends on the previous apparition of mutant replicases (in the previous case the two events were independent), results in a discrepancy at large dilution factor and large maturation time where mutant parasites can survive contrary to model A (explaining the increasing tren for *z*_1_ at large *T*_*mat*_). Indeed, for large maturation times, even if mutant replicases dominate the system, they will mutate to parasites and avoid their extinction. Again, the wild-type populations go extinct at high dilution factors and the mutant replicases dominate because they can replicate by themselves and parasites will be isolated because of dilution. Larger maturation times favour the appearance and survival of parasites in the system. As observed in Fig. 18, the intermediate steady co-existence is obtained over a wider range of dilution factors.

#### With stirring

##### Model A

We can study how stirring modifies the results. The maturation phase can be treated in the same manner, but the fractions will be different for each compartment. Thus, with 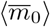 the mean number of WT replicases after the next maturation phase, we have

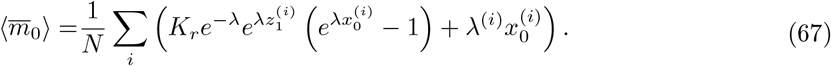

The sum is performed over the *N* compartments. We recognise the result from the precedent model but the fractions are now cell-dependent, and we average over all cells. Obviously, for *s* = 1 (complete stirring), all the terms in the sum are equal and the system loses its memory. In this case we recover the previous result with total pooling. On the other end, if *s* = 0 (no stirring), all the compartments are conserved in time and the system does not evolve (except for mutations). We can simulate these equations, assuming deterministic maturation and keeping the stochasticity induced by stirring. We compare this semi-deterministic method to the previous deterministic one (complete pooling) for different values of *s* in Fig. 19. We observe that for unperfect stirring *s <* 1, parasite fractions tend to increase and replicases become a minority in the whole population.

We observe that stirring (and thus uncomplete pooling) allows to preserve WT populations from extinction and keeps the system away from steady state. In the limit of strong stirring we recover the complete pooling behaviour. We can plot coexistence diagrams for different stirring coefficients *s* and compare to the deterministic one Fig. 15. We obtain Fig. 20 for *s* = 0.5. This coexistence diagram is built from simulations where maturation is completely deterministic but every compartment must be modeled and has a stochastic inoculation. In this case, the diagram depends more on maturation time than dilution, indeed the system has a stronger memory when *s <* 1.

**Figure S15.**
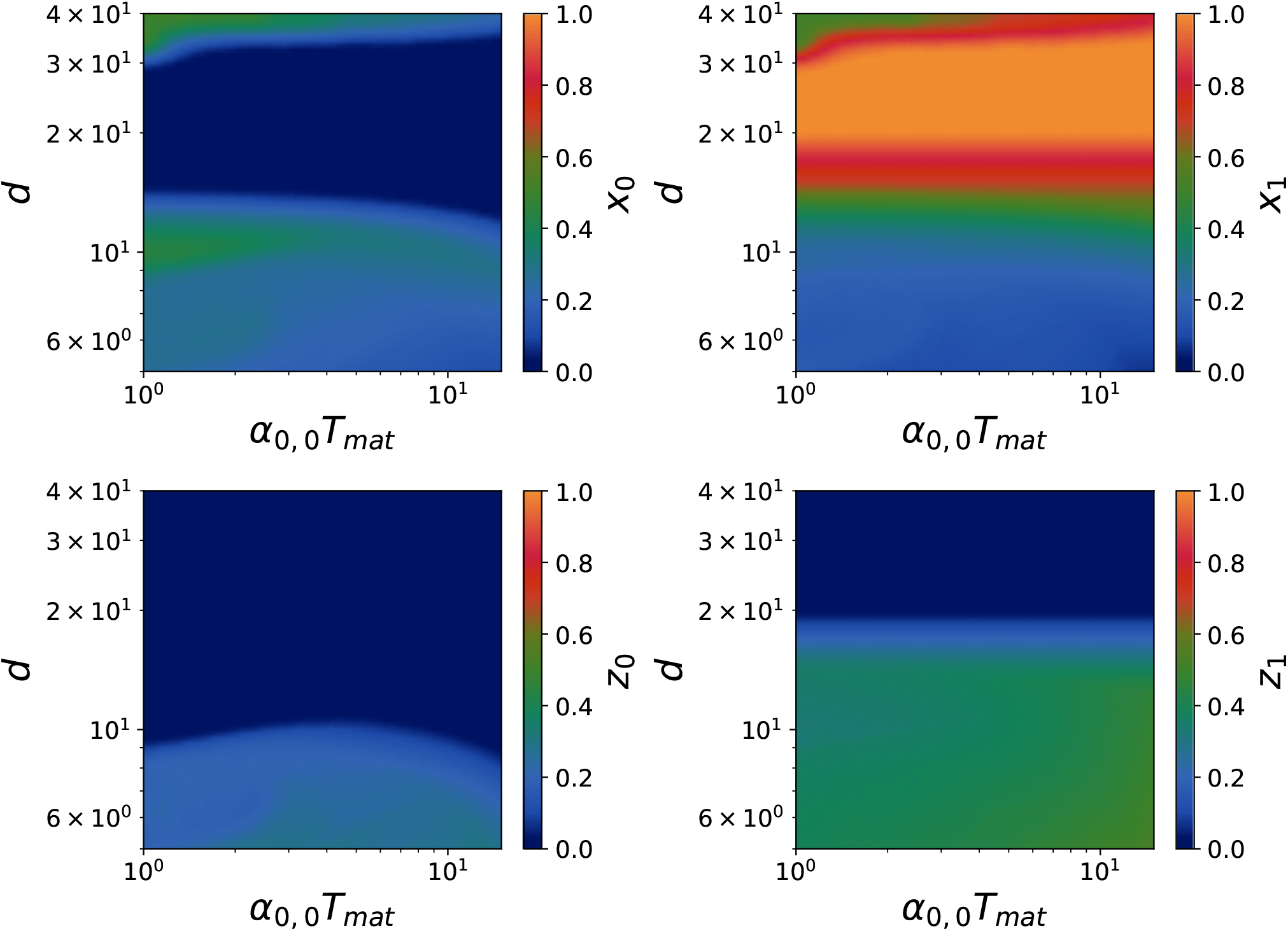
Coexistence diagram for the fractions in model A (theoretical expectations) after 300 cycles. We used for replication rates *α*_0,0_ = 5 × 10^−1^ *h*^−1^, *α*_1,1_ = 2 *h*^−1^, *γ*_0,0_ = 7 *h*^−1^, *γ*_1,1_ = 20 *h*^−1^, and for mutation rates *µ*_0_ = 7 × 10^−3^ *h*^−1^, *ν*_0_ = 7 × 10^−3^ *h*^−1^. The carrying capacities are *K*_*p*_ = *K*_*r*_ = 30 and dilution factor *d* = 10. The panels represent the fractions for *x*_0_, *x*_1_, *z*_0_, *z*_1_ from right to left and from top to bottom.

**Figure S16.**
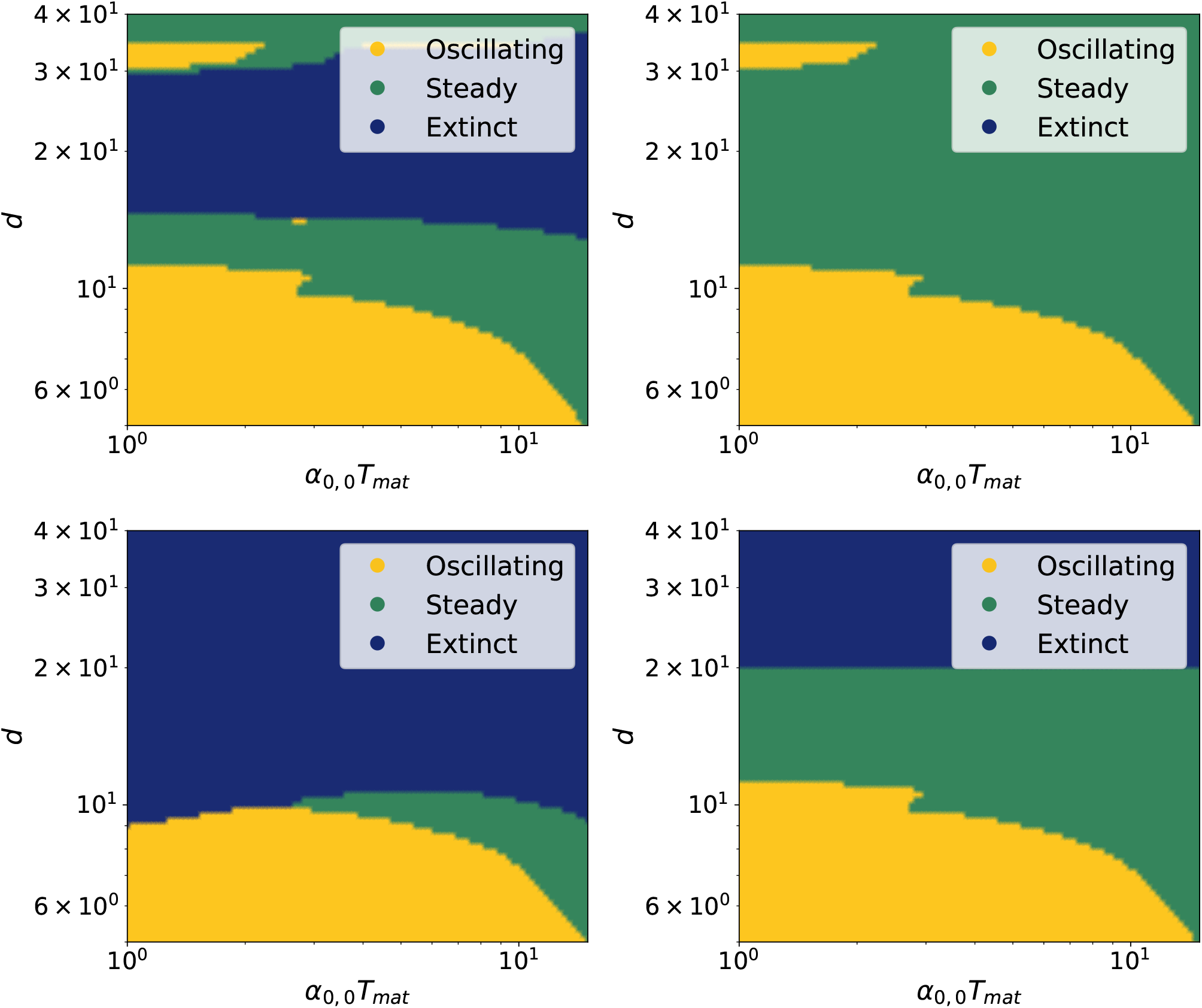
Coexistence diagram of the states of the system in model A (theoretical expectations) after 300 cycles. We used for replication rates *α*_0,0_ = 5 × 10^−1^ *h*^−1^, *α*_1,1_ = 2 *h*^−1^, *γ*_0,0_ = 7 *h*^−1^, *γ*_1,1_ = 20 *h*^−1^, and for mutation rates *µ*_0_ = 7 × 10^−3^ *h*^−1^, *ν*_0_ = 7 × 10^−3^ *h*^−1^. The carrying capacities are *K*_*p*_ = *K*_*r*_ = 30 and dilution factor *d* = 10. The panels represent the fractions for *x*_0_, *x*_1_, *z*_0_, *z*_1_ from right to left and from top to bottom.

**Figure S17.**
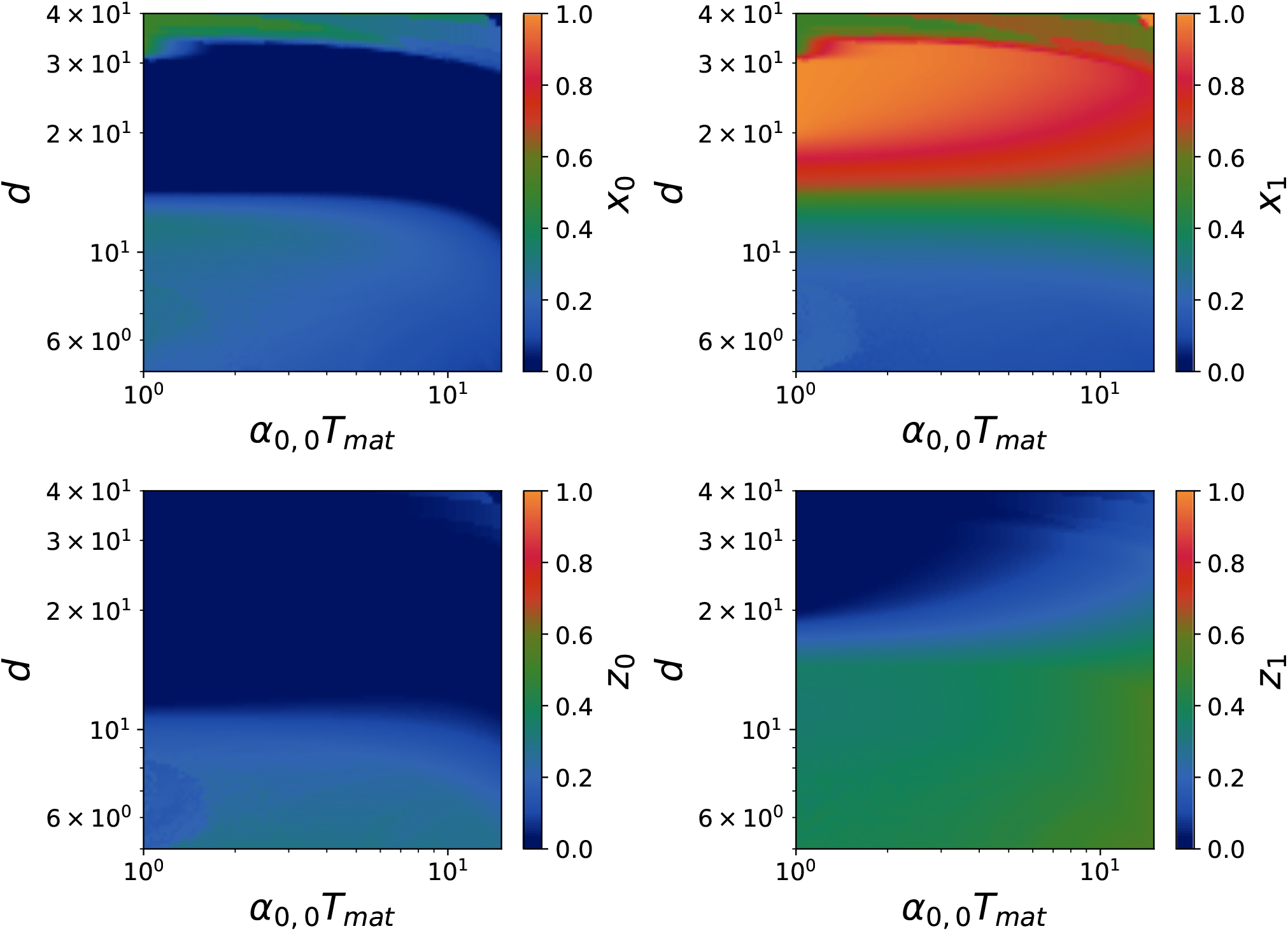
Coexistence diagram for the fractions with model B where hosts mutate to parasites (theoretical expectations) after 300 cycles. We used for replication rates *α*_0,0_ = 5 × 10^−1^ *h*^−1^, *α*_1,1_ = 2 *h*^−1^, *γ*_0,0_ = 7 *h*^−1^, *γ*_1,1_ = 20 *h*^−1^, and for mutation rates *µ*_0_, *ν*_0_, *ν*_1_ = 7 × 10^−3^ *h*^−1^. The carrying capacities are *K*_*p*_ = *K*_*r*_ = 30 and dilution factor *d* = 10. The panels represent the fractions for *x*_0_, *x*_1_, *z*_0_, *z*_1_ from right to left and from top to bottom.

**Figure S18.**
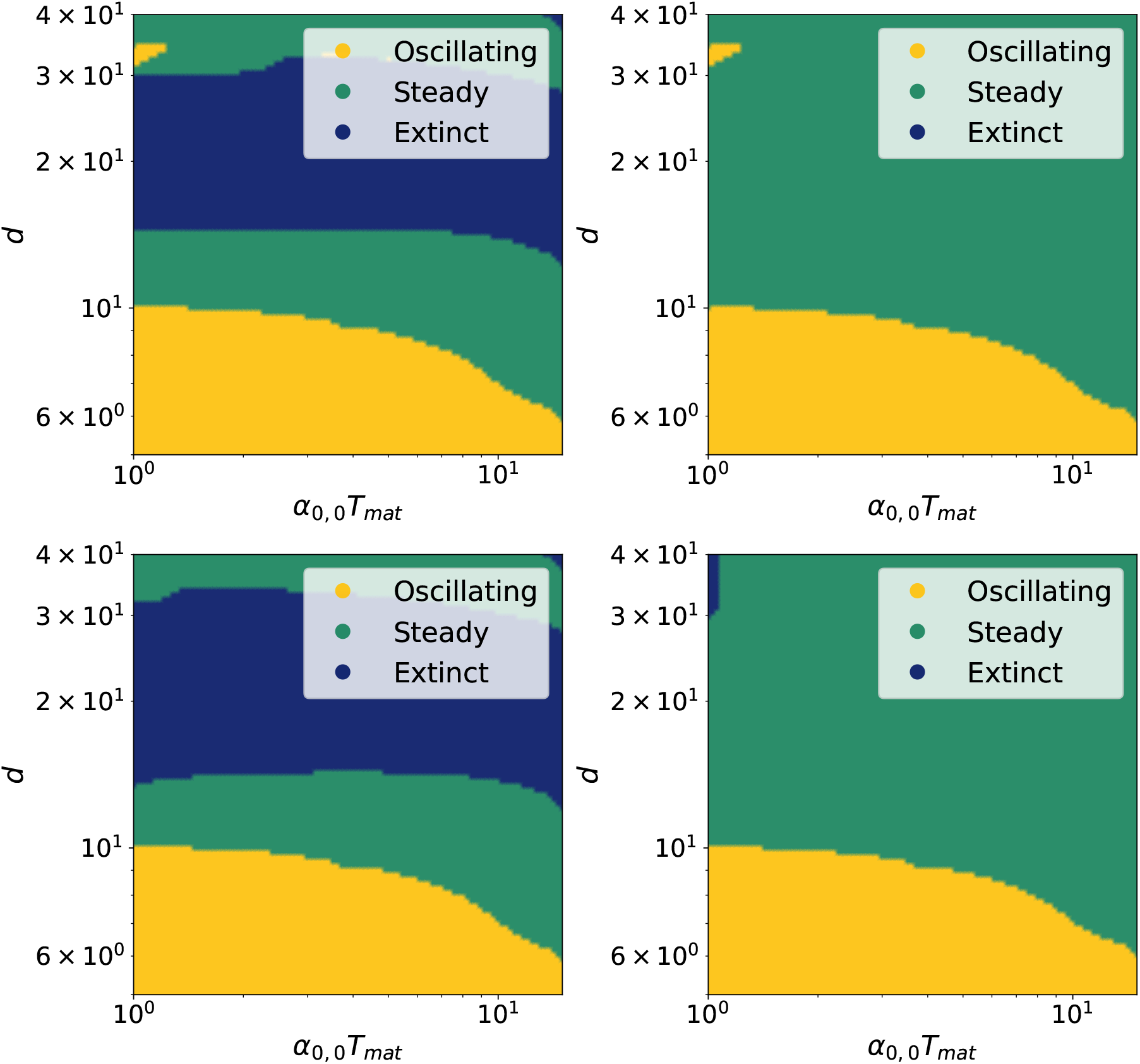
Coexistence diagram of the states of the system in model B where hosts mutate to parasites (theoretical expectations) after 300 cycles. We used for replication rates *α*_0,0_ = 5 × 10^−1^ *h*^−1^, *α*_1,1_ = 2 *h*^−1^, *γ*_0,0_ = 7 *h*^−1^, *γ*_1,1_ = 20 *h*^−1^, and for mutation rates *µ*_0_ = 7 × 10^−3^ *h*^−1^, *ν*_0_ = 7 × 10^−3^ *h*^−1^, *ν*_1_ = 7 × 10^−3^ *h*^−1^. The carrying capacities are *K*_*p*_ = *K*_*r*_ = 30 and dilution factor *d* = 10. The panels represent the fractions for *x*_0_, *x*_1_, *z*_0_, *z*_1_ from right to left and from top to bottom.

**Figure S19.**
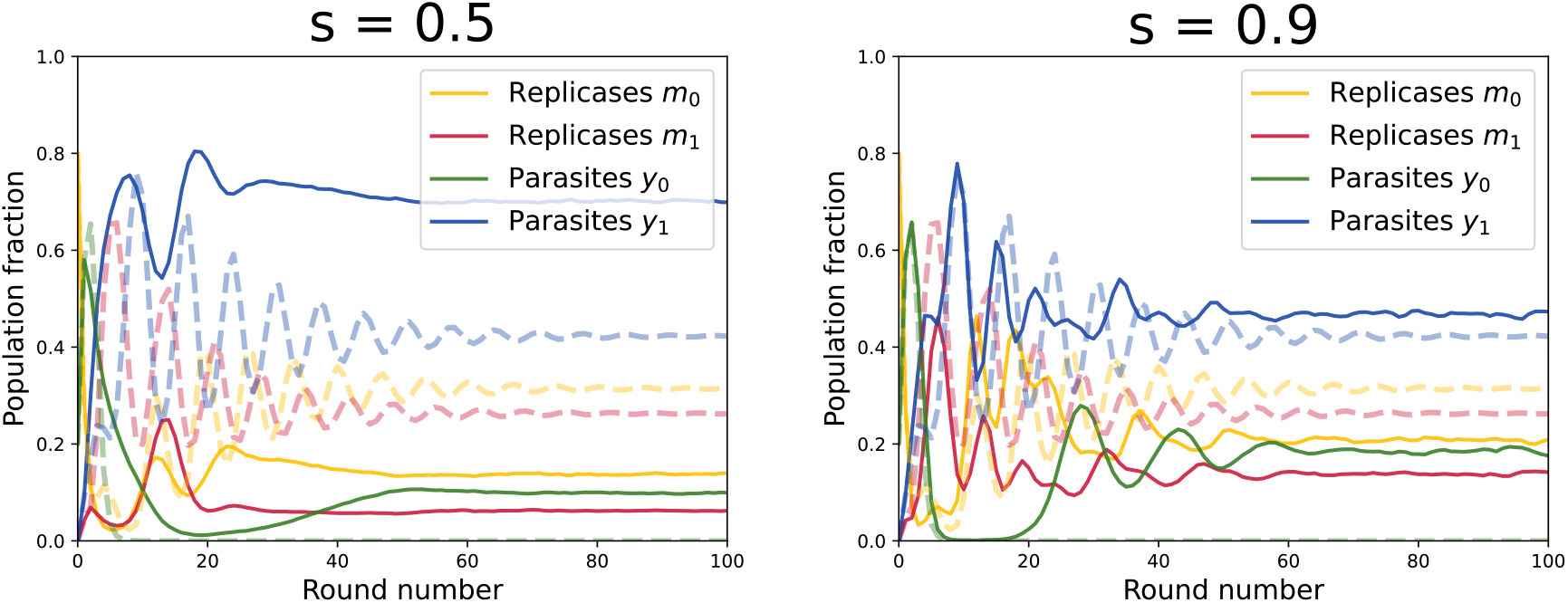
Comparison between complete pooling dynamics and stirring (simulations with stirring were performed with 50000 compartments). In experiments compartments are reinitialized by stirring, we model this in the last section. The stronger is stirring the more the system has a memory. This could explain discrepancies between our model of complete stirring and experiments. We used for replication rates *α*_0,0_ = 5 × 10^−1^ *h*^−1^, *α*_1,1_ = 2 *h*^−1^, *γ*_0,0_ = 7 *h*^−1^, *γ*_1,1_ = 20 *h*^−1^, and for mutation rates *µ*_0_ = 7 × 10^−3^ *h*^−1^, *ν*_0_ = 7 × 10^−3^ *h*^−1^. The carrying capacities are *K*_*p*_ = *K*_*r*_ = 30 and dilution factor *d* = 10. The panels represent different values of the stirring parameter *s*.

Weaker stirring tends to favour parasites, and the proportion of mutant parasites in the whole population then depends on the maturation time. We also recover the complete pooling limit for *s* → 1. If the system has a strong memory, dilution only weakly affects the fractions, but mutations occur even if the compartment is not diluted. We can compare this to the coexistence diagram with *s* = 0.95 (higher stirring) in Fig. 21, for large dilution factors and stronger stirring (*s* = 0.99 instead of *s* = 0.8), mutant replicases start dominating the system anew. For too large values of dilution factors, the finite number of compartments result in the extinction of the population, explaining that all species go to 0 at large *d*. Most of the discrepancies observed at small dilution factors between Fig. 21 and Fig. 15 results from a finite size effect in simulations with stirring (finite number of compartments in the simulations). Indeed, if we set a threshold on the fraction under which a population goes extinct (resulting from the finite number of compartments) in the deterministic, homogeneous model with an infinite number of compartments, we recover coexistence diagrams similar to that with stirring and a finite number of compartments.

##### Model B

We describe stirring in the same way for model B. For a stirring coefficient *s* = 0.5 (resp. *s* = 0.95), we obtain the coexistence diagram Fig. 22 (resp. Fig. 23). We additionally illustrate the discrepancy between a perfectly homogeneous stirring and the cases *s* = 0.5 and *s* = 0.95 in Fig. 24. As observed previously, the steady state fractions are modified due to stirring, allowing for a stronger resilience of wild-type species. In this model, increasing *s* allows to recover a phase where mutant replicases dominate at large dilution factors.

Replicases are mostly extinct when *s* is too weak in this model as well. Again, we observe in Fig. 22 that the coexistence diagram mostly depends on the maturation time, as dilution is no longer relevant if the system has a memory. With a weaker stirring, parasites are favored, because parasites will less likely be isolated from replicases. We draw the same conclusion as for model A concerning weaker stirring : for *s <* 1, parasite fractions are higher (it is clearly seen in Fig. 24.

**Figure S20.**
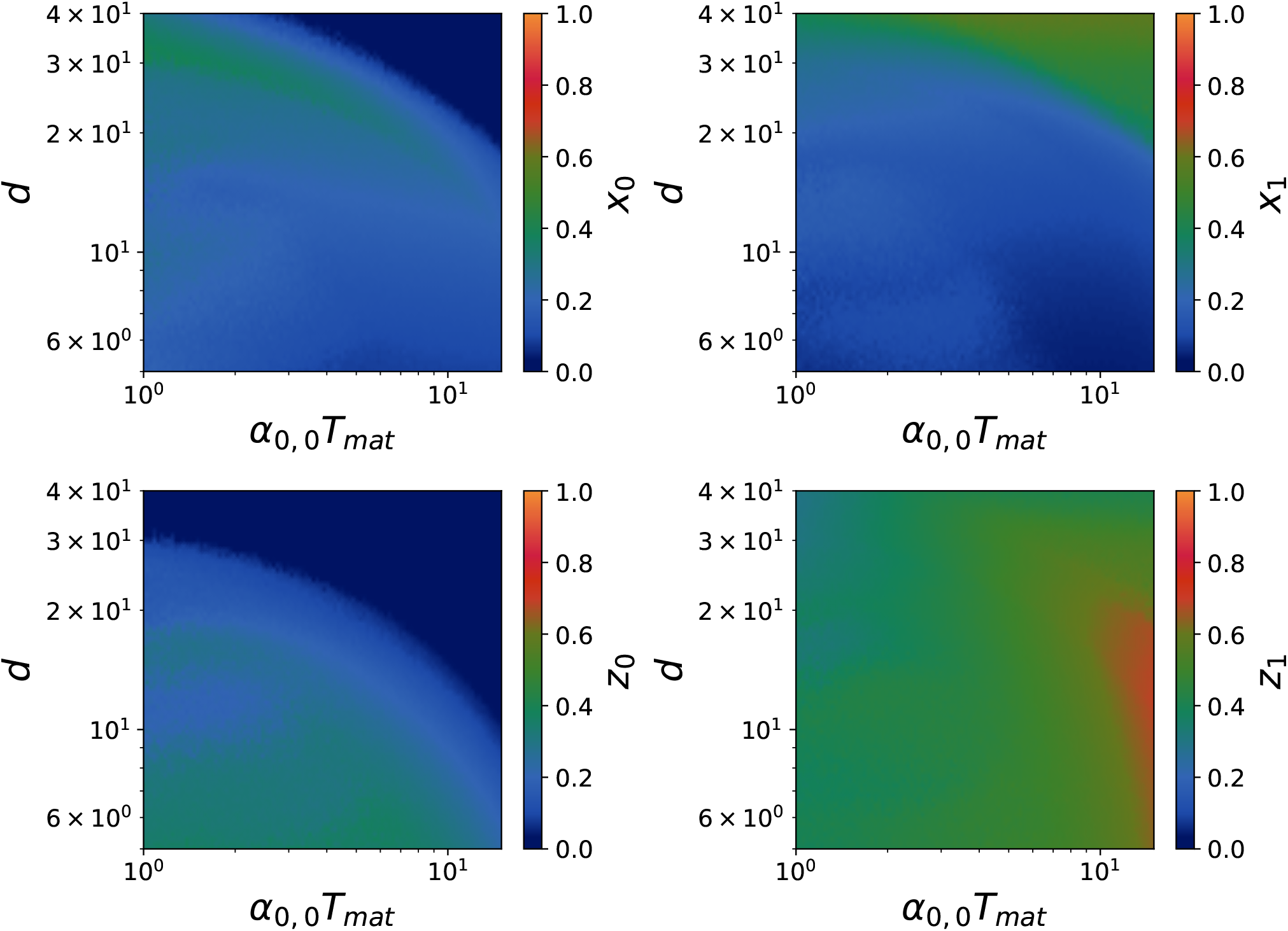
Coexistence diagram with ***s* = 0.7** for the fractions in model A with stirring after 100 cycles with 8000 compartments. We used for replication rates *α*_0,0_ = 5 × 10^−1^ *h*^−1^, *α*_1,1_ = 2 *h*^−1^, *γ*_0,0_ = 7 *h*^−1^, *γ*_1,1_ = 20 *h*^−1^, and for mutation rates *µ*_0_ = 7 × 10^−3^ *h*^−1^, *ν*_0_ = 7 × 10^−3^ *h*^−1^. The carrying capacities are *K*_*p*_ = *K*_*r*_ = 30 and dilution factor *d* = 10. The panels represent the fractions for *x*_0_, *x*_1_, *z*_0_, *z*_1_ from right to left and from top to bottom.

**Figure S21.**
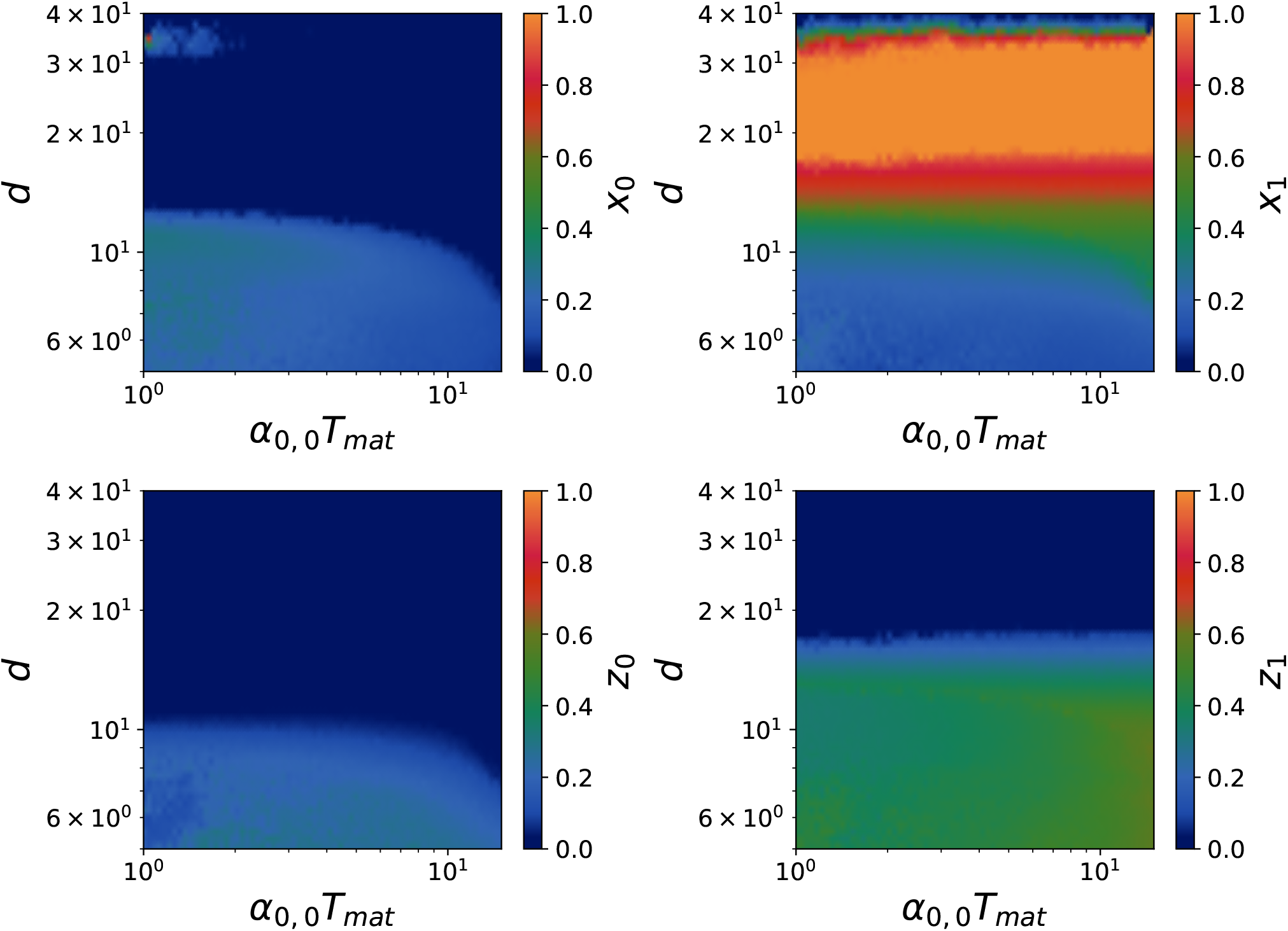
Coexistence diagram with ***s* = 0.95** for the fractions in model A with stirring after 100 cycles with 8000 compartments. We used for replication rates *α*_0,0_ = 5 × 10^−1^ *h*^−1^, *α*_1,1_ = 2 *h*^−1^, *γ*_0,0_ = 7 *h*^−1^, *γ*_1,1_ = 20 *h*^−1^, and for mutation rates *µ*_0_ = 7 × 10^−3^ *h*^−1^, *ν*_0_ = 7 × 10^−3^ *h*^−1^. The carrying capacities are *K*_*p*_ = *K*_*r*_ = 30 and dilution factor *d* = 10. The panels represent the fractions for *x*_0_, *x*_1_, *z*_0_, *z*_1_ from right to left and from top to bottom.

**Figure S22.**
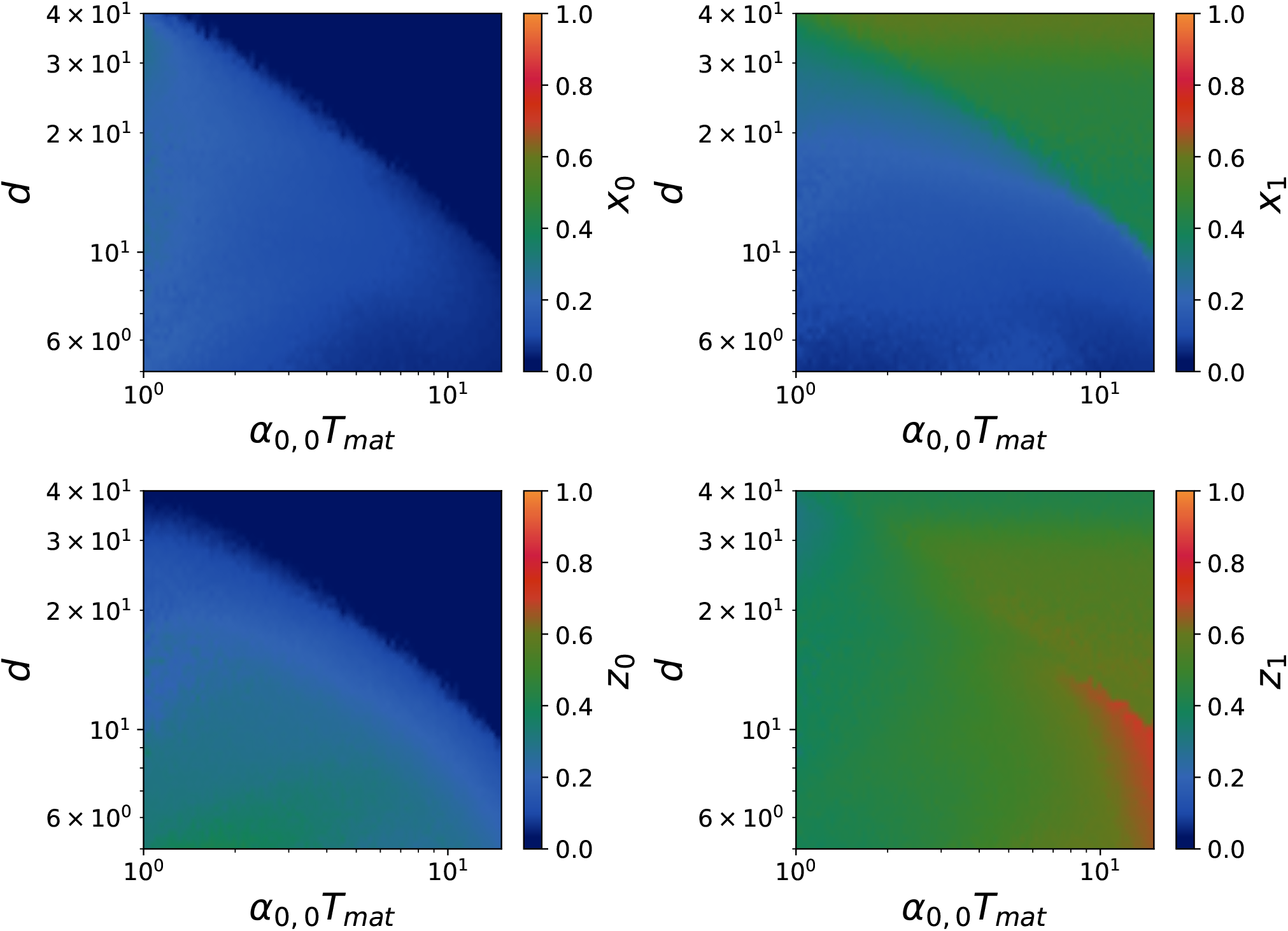
Coexistence diagram with ***s* = 0.7** for the fractions in model B with stirring after 100 cycles with 8000 compartments, where *α*_0,0_ = 5 × 10^−1^ *h*^−1^, *α*_1,1_ = 2 *h*^−1^, *γ*_0,0_ = 7 *h*^−1^, *γ*_1,1_ = 20 *h*^−1^, and *µ*_0_, *ν*_0_, *ν*_1_ = 7 × 10^−3^ *h*^−1^. The carrying capacities are *K*_*p*_ = *K*_*r*_ = 30 and dilution factor *d* = 10. The panels represent the fractions for *x*_0_, *x*_1_, *z*_0_, *z*_1_ from right to left and from top to bottom.

**Figure S23.**
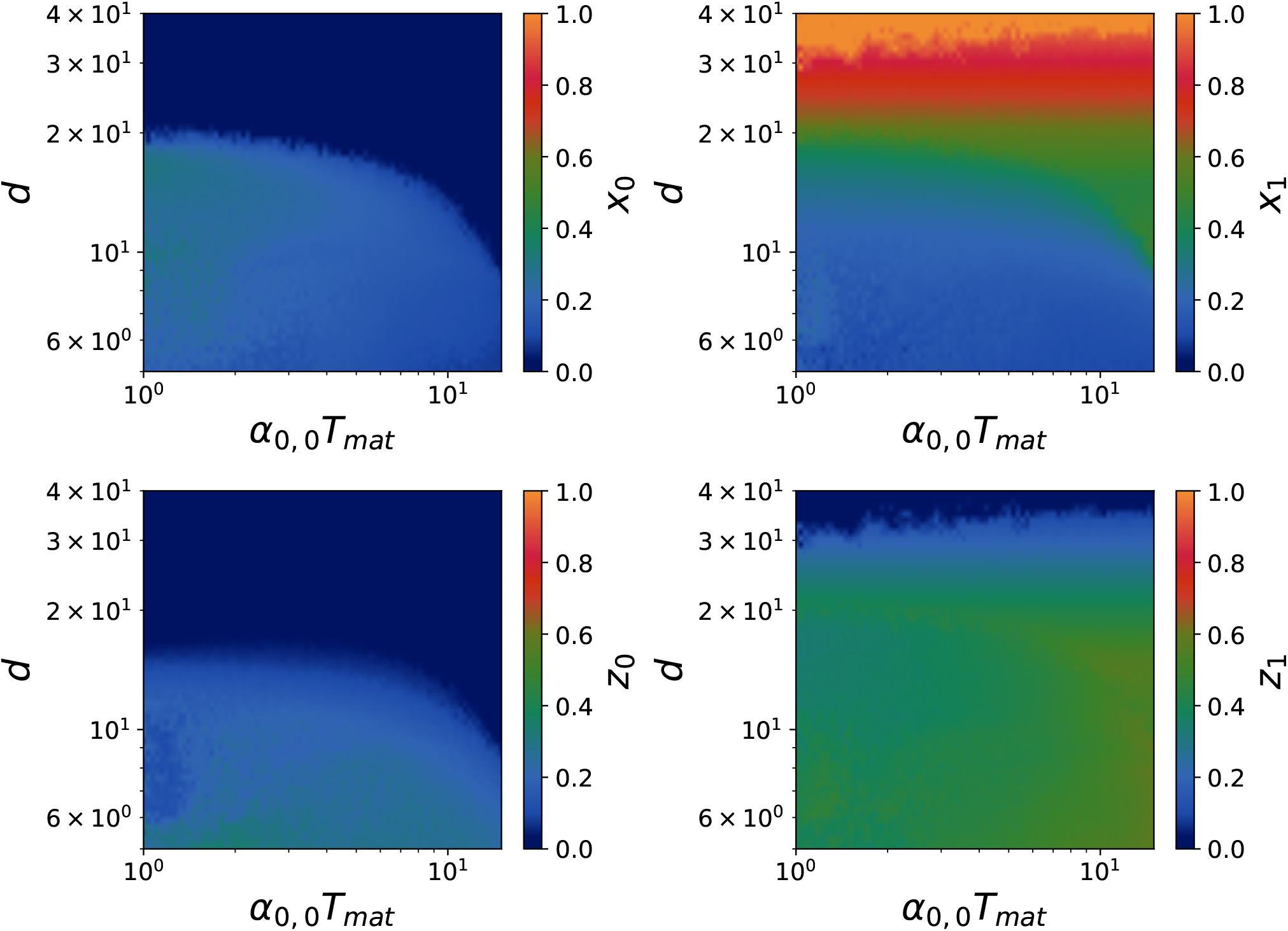
Coexistence diagram with ***s* = 0.95** for the fractions in model B with stirring after 100 cycles with 8000 compartments, where *α*_0,0_ = 5 × 10^−1^ *h*^−1^, *α*_1,1_ = 2 *h*^−1^, *γ*_0,0_ = 7 *h*^−1^, *γ*_1,1_ = 20 *h*^−1^, and *µ*_0_, *ν*_0_, *ν*_1_ = 7 × 10^−3^ *h*^−1^. The carrying capacities are *K*_*p*_ = *K*_*r*_ = 30 and dilution factor *d* = 10. The panels represent the fractions for *x*_0_, *x*_1_, *z*_0_, *z*_1_ from right to left and from top to bottom.

**Figure S24.**
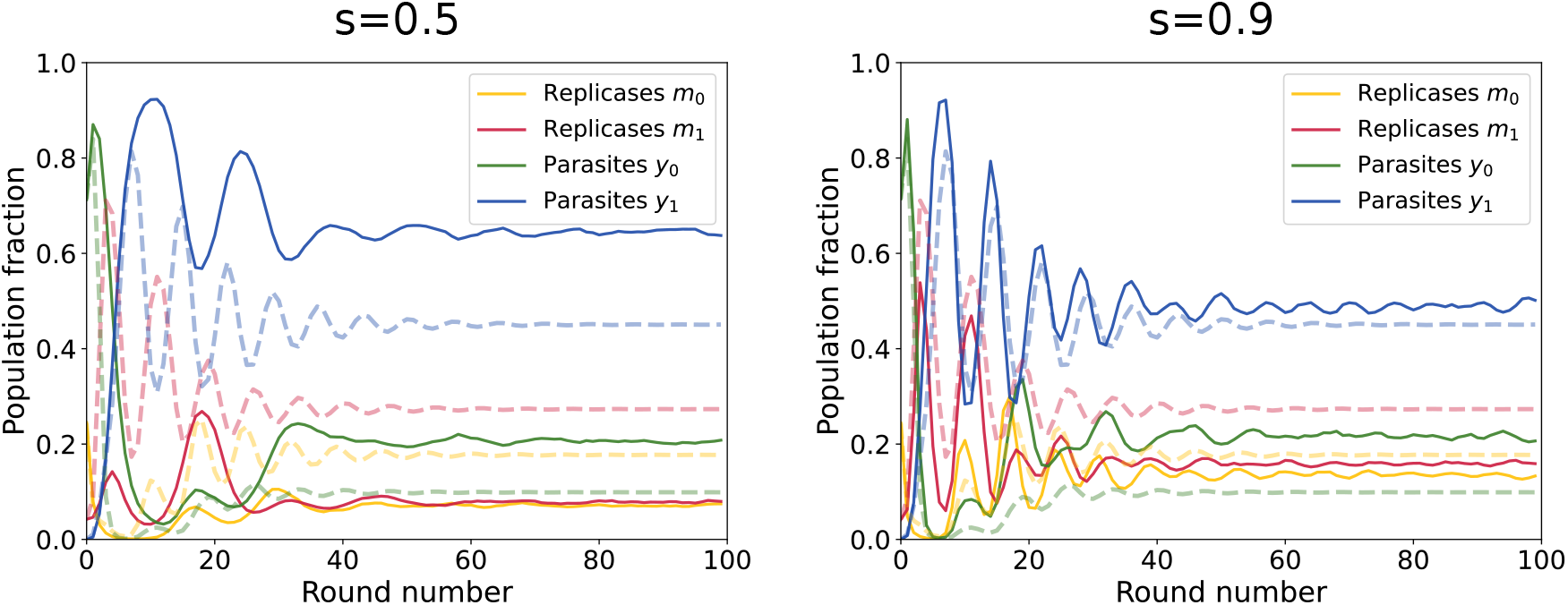
Illustration of the discrepancy between *s* = 1 and imperfect stirring with model B for *α*_0,0_ = 5 × 10^−1^ *h*^−1^, *α*_1,1_ = 2 *h*^−1^, *γ*_0,0_ = 7 *h*^−1^, *γ*_1,1_ = 20 *h*^−1^., and *µ*_0_, *ν*_0_ = *ν*_1_ = 7 × 10^−3^ *h*^−1^ and 50000 compartments. The carrying capacities are *K*_*p*_ = *K*_*r*_ = 30 and dilution factor *d* = 10. The panels represent different values of the sirring parameter *s*.

### Generalist parasites

#### Generalist parasites versus specific parasites (model A)

Generalist parasites evolve in such a way that they remain able to replicate using wild-type replicases as well as mutant replicases. Compartments with only WT replicases and mutant parasites will be filled with *y*_1_ thus leading to the extinction of WT replicases. In this case *γ* is no longer diagonal but upper triangular. With one level of mutations, it means *γ*_0,1_ ∼ *γ*_0,0_ and we obtain new deterministic rules:

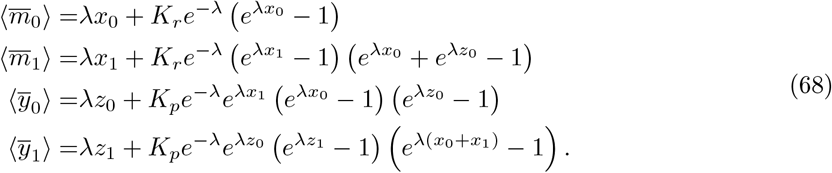

This leads to an easier extinction of wild-type populations, as the fractions are given by:

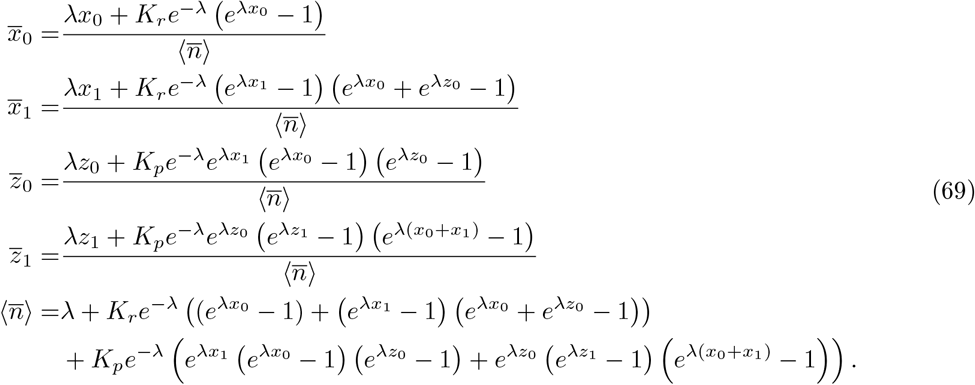

The effect of mutations is also slightly modified in this situation as the conditions for a population taking over the compartment are different. In particular, we obtain the average numbers of mutations

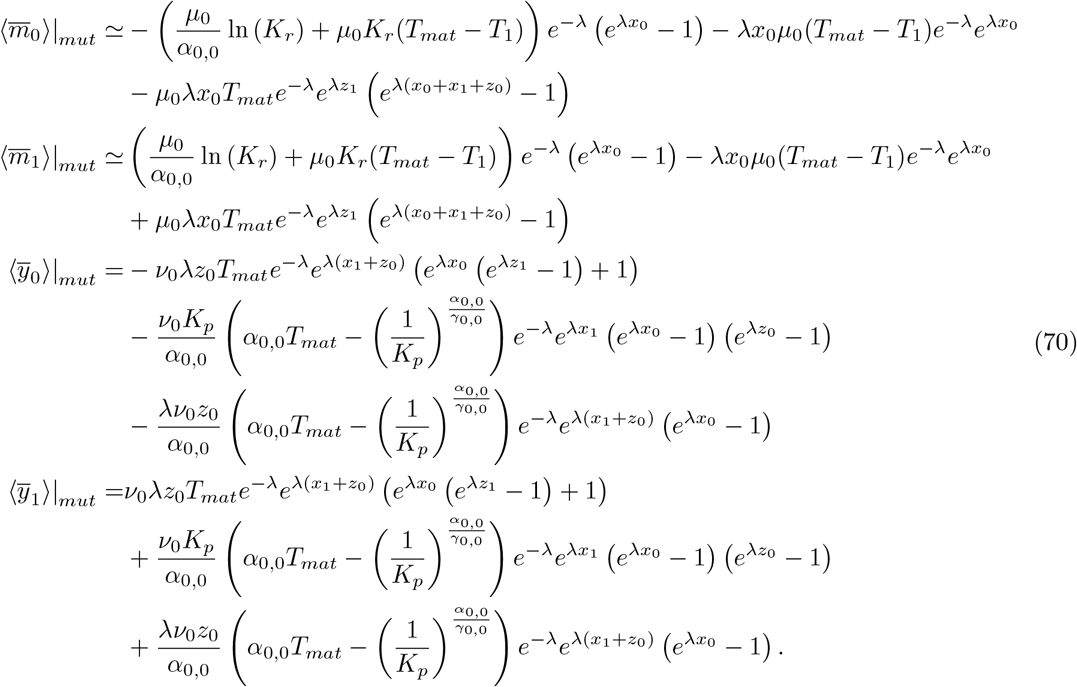

From this we can plot new coexistence diagrams in the case of general mutations in Fig. 25. We observe that for generalist parasites, the results become mostly independent of the maturation time (comparing to Fig. 15), especially at large dilution factors. The dilution factor modifies the state of the system and wild-type species are extinct in most of the diagram. Indeed generalist parasites can replicate with any replicase and the reason why WT replicases could survive with specific parasites was that whenever a compartment contained WT replicases and mutant parasites, the replicases were taking over the compartment. This leads to the extinction of wild-type replicases (for generalist parasites), and thus of wild-type parasites that cannot replicate on their own. It is enough to have non zero mutation rates to have mutants taking over the system unless dilution is weak enough to keep wild-type individuals in the system or large enough to wash out all mutants at each cycle.

#### Generalist parasites versus specific parasites (model B)

We can also compare the case where the mutant parasite is generalist to that where it is specific to the mutant replicases in model B. Generalist parasites will be able to take over the system more easily as seen in the previous case. However the difference lies in the fact that mutant parasites can only appear from mutant replicases. We build a new coexistence diagram in this case in Fig. 26. In this model, mutations are also modified as

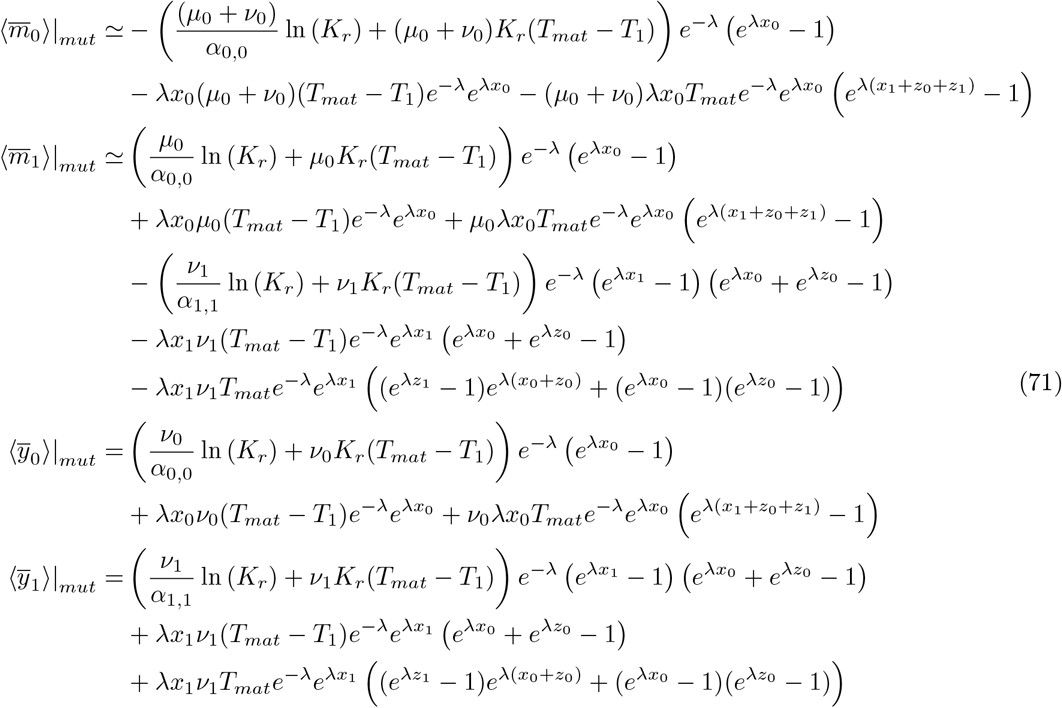

Again, we observe that the WT species go extinct when the mutant parasite is generalist in most of the diagram. We also recover that the diagram becomes mostly independent of maturation time, as mutant parasites can replicate with every hosts and do not need to reach a threshold in the total population to survive. However, for model B, there is a dependency in maturation time as mutant replicases can still mutate to parasites. For large maturation time, the fraction of parasites becomes larger. At large dilution factors, mutants are washed out at each cycle and wild-type replicases can survive.

**Figure S25.**
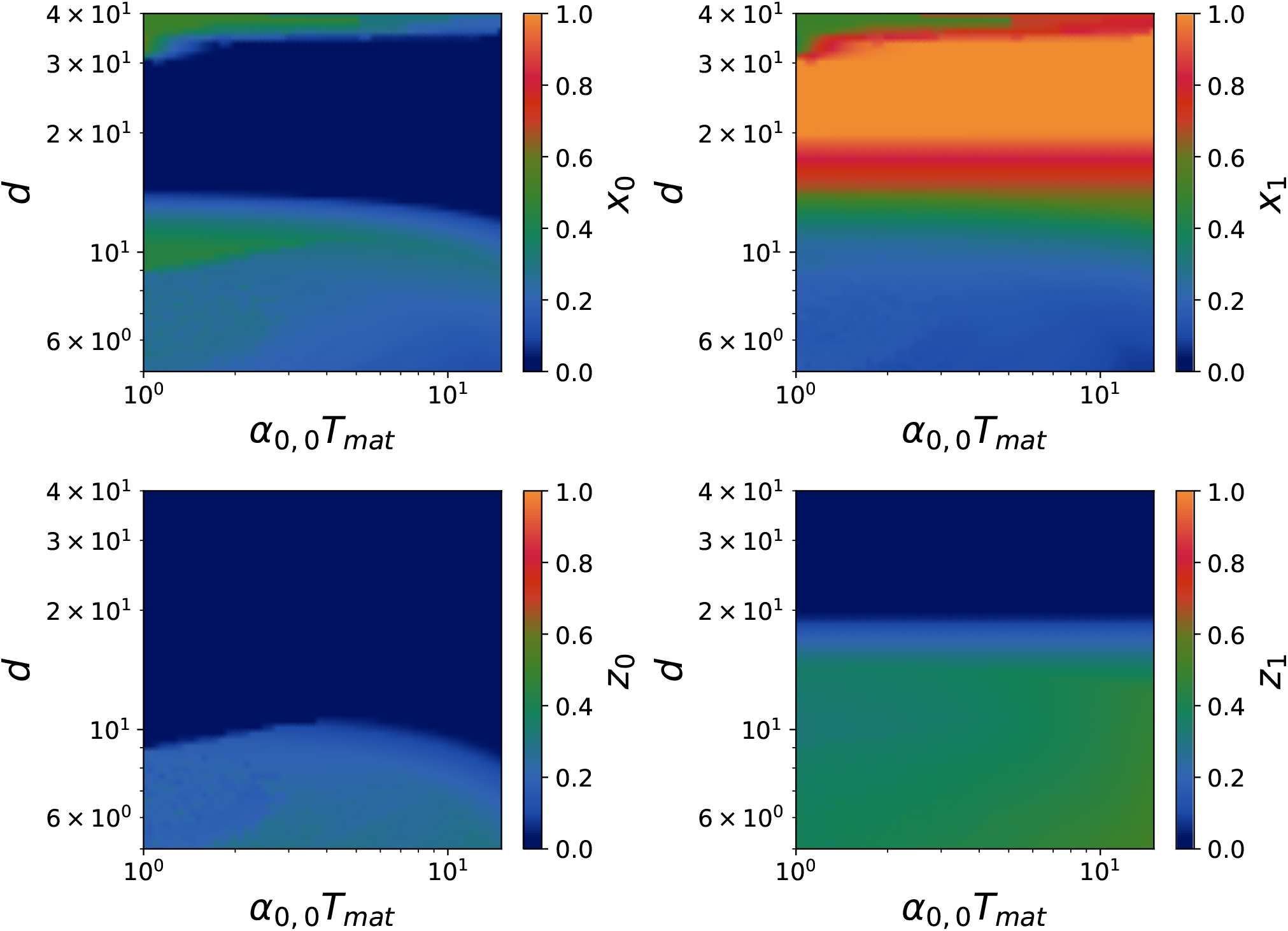
Coexistence diagram for the fractions in model A with generalist parasites (theoretical expectations) after 300 cycles. We used for replication rates *α*_0,0_ = 5 × 10^−1^ *h*^−1^, *α*_1,1_ = 2 *h*^−1^, *γ*_0,0_ = 7 *h*^−1^, *γ*_1,1_ = 20 *h*^−1^, and for mutation rates *µ*_0_ = 7 × 10^−3^ *h*^−1^, *ν*_0_ = 7 × 10^−3^ *h*^−1^. The carrying capacities are *K*_*p*_ = *K*_*r*_ = 30 and dilution factor *d* = 10. The panels represent the fractions for *x*_0_, *x*_1_, *z*_0_, *z*_1_ from right to left and from top to bottom.

**Figure S26.**
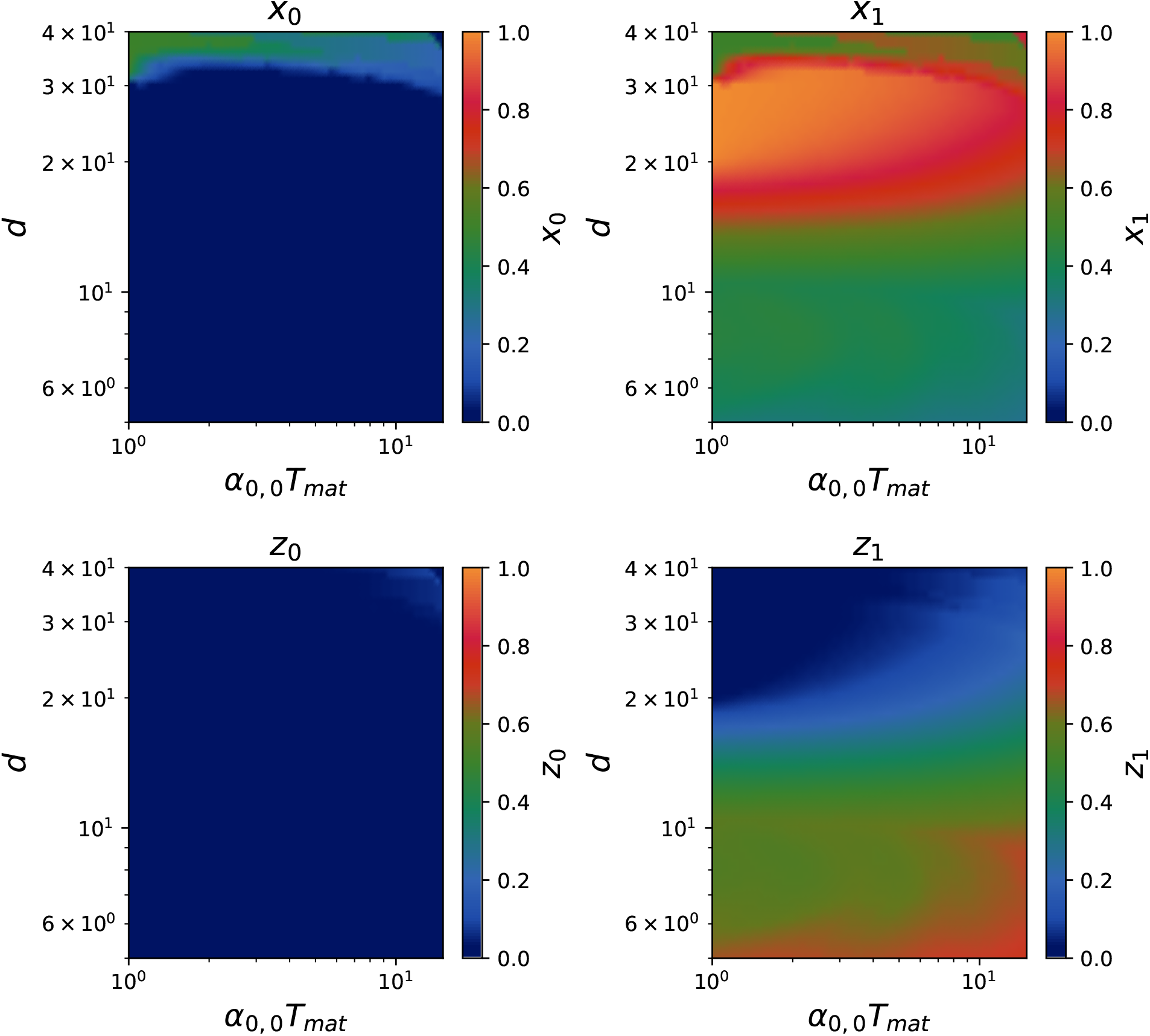
Coexistence diagram in model B with generalist parasites (theoretical expectations) after 300 cycles, where *α*_0,0_ = 5 × 10^−1^ *h*^−1^, *α*_1,1_ = 2 *h*^−1^, *γ*_0,0_ = 7 *h*^−1^, *γ*_1,1_ = 20 *h*^−1^, and *µ*_0_, *ν*_0_, *ν*_1_ = 7 × 10^−3^ *h*^−1^. The carrying capacities are *K*_*p*_ = *K*_*r*_ = 30 and dilution factor *d* = 10. The panels represent the fractions for *x*_0_, *x*_1_, *z*_0_, *z*_1_ from right to left and from top to bottom.

### Improvement of resistance (model B)

We can build another model where hosts mutate in order to acquire a better resistance to parasites, at the cost of a smaller replication rate. In order to model this with the deterministic approach we can assume the following rules: (1) If a compartment is initially filled with wild-type hosts, and no wild-type parasites (regardless of the rest), the hosts replicate until they reach the carrying capacity. (2) If a compartment is initially filled with wild-type hosts and wild-type parasites (regardless of the rest), the wild-type parasites replicate until they reach the carrying capacity. (3) If a compartment has no wild-type hosts, no mutant parasites, and is filled with mutant hosts (regardless of the amount of wild-type parasites), mutant hosts will replicate until they reach the carrying capacity. (4) If a compartment has no wild-type hosts, and is filled with mutant hosts and mutant parasites (regardless of the amount of wild-type parasites), we assume that mutant hosts and parasites will replicate together until the whole population reaches the carrying capacity (thus hosts grows by *K*_*r*_*/*2 individuals and parasites grows by *K*_*p*_*/*2 individuals). From those rules, we can compute the new updated average numbers after maturation:

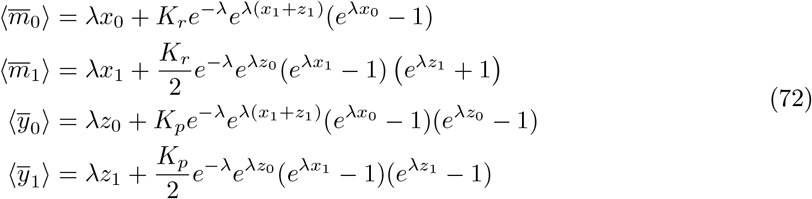

From this we can plot new coexistence diagrams (see Fig. 27), at intermediate dilution factors wild-type replicases take over as they have the highest relative fitness and are able to self-replicate. Yet at high dilution factors, mutant replicases are likely to be alone in a compartment, facilitating their growth. Therefore we get a co-existence of wild-type and mutant replicases at large dilution. Mutant parasites remain a minority as they do not have a strong relative fitness as compared to mutant replicases (due to the increased resistance of mutant replicases). We recover the influence of maturation time which favors mutant populations.

### Trade-off Generality/Replicability (model B)

We can also study what happens when parasites mutate to acquire generality, at the cost of replicability. In this case the deterministic rules are: (1)In a compartment with WT replicases and WT parasites, WT parasites take over the cell whatever the concentration of other species. (2)In a compartment with WT replicases and mutant replicases only, mutant replicases take over the compartment. (3)In a compartment with mutant parasites and WT replicases only, both populations grow until they reach half the carrying capacity. (4)In a compartment with mutant parasites, and mutant replicases but no WT replicases or no WT mutants, both populations grow until they reach half the carrying capacity. From those new rules, we can compute the new updated fractions (independent of the model):

**Figure S27.**
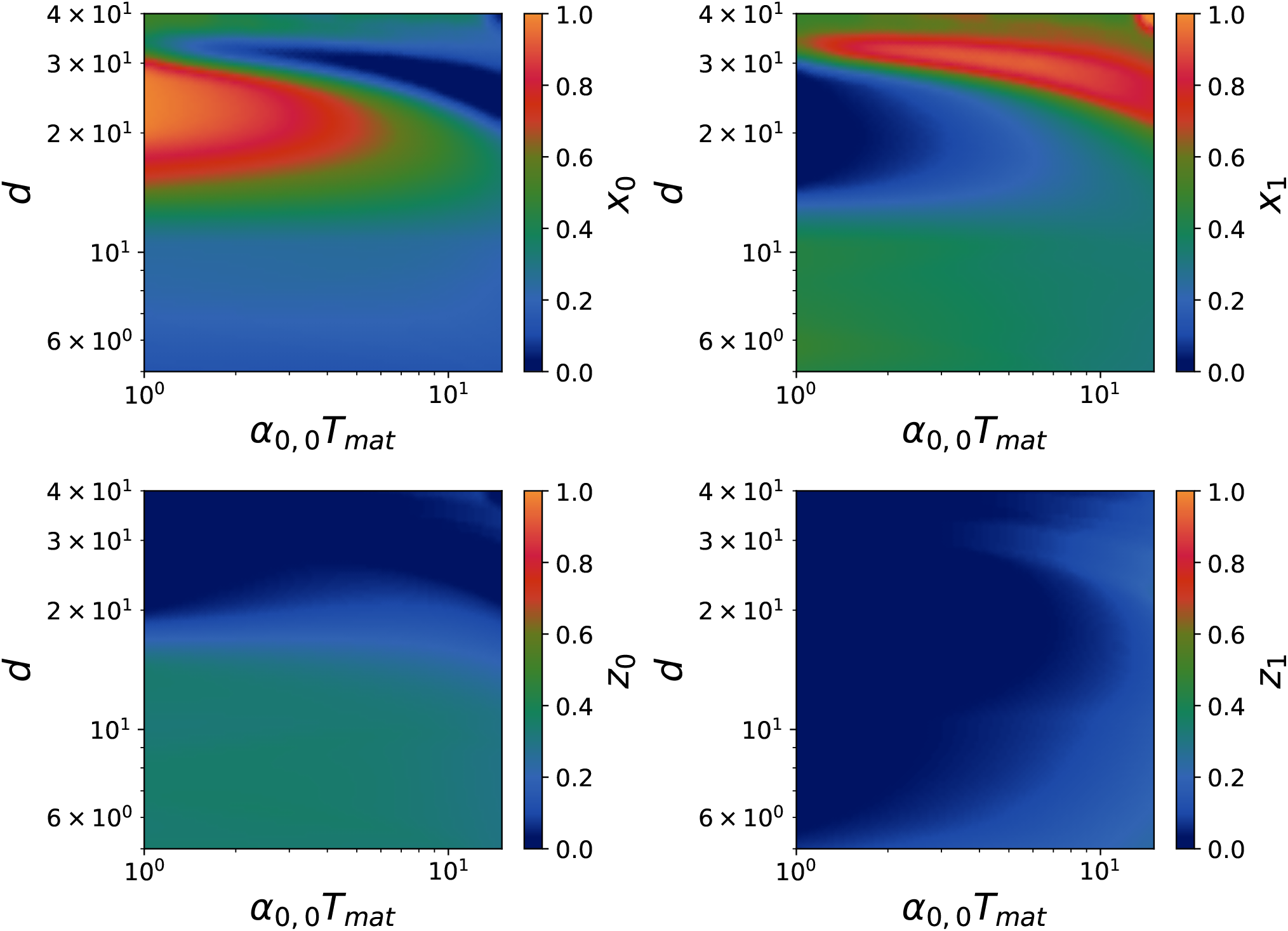
Coexistence diagram of the fractions of each species in the system with the improvement of resistance in model B (theoretical expectations) after 300 cycles. We used for replication rates *α*_0,0_ = 5 × 10^−1^ *h*^−1^, *α*_1,1_ = 5 × 10^−1^ *h*^−1^, *γ*_0,0_ = 10 *h*^−1^, *γ*_1,1_ = 5 × 10^−1^ *h*^−1^, and for mutation rates *µ*_0_ = 7 × 10^−3^ *h*^−1^, *ν*_0_ = 7 × 10^−3^ *h*^−1^. The carrying capacities are *K*_*p*_ = *K*_*r*_ = 30 and dilution factor *d* = 10. The panels represent the fractions for *x*_0_, *x*_1_, *z*_0_, *z*_1_ from right to left and from top to bottom.

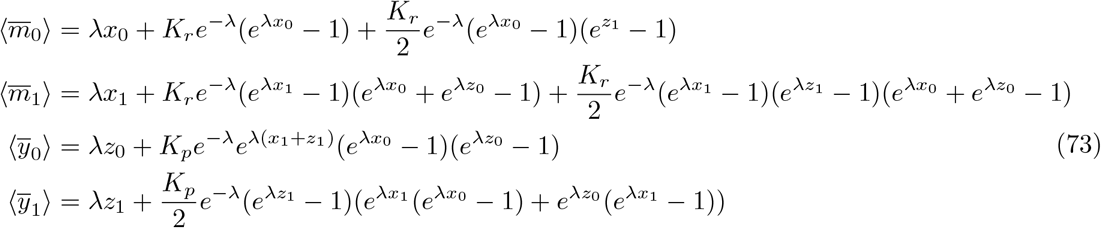

We also have the numbers of mutants

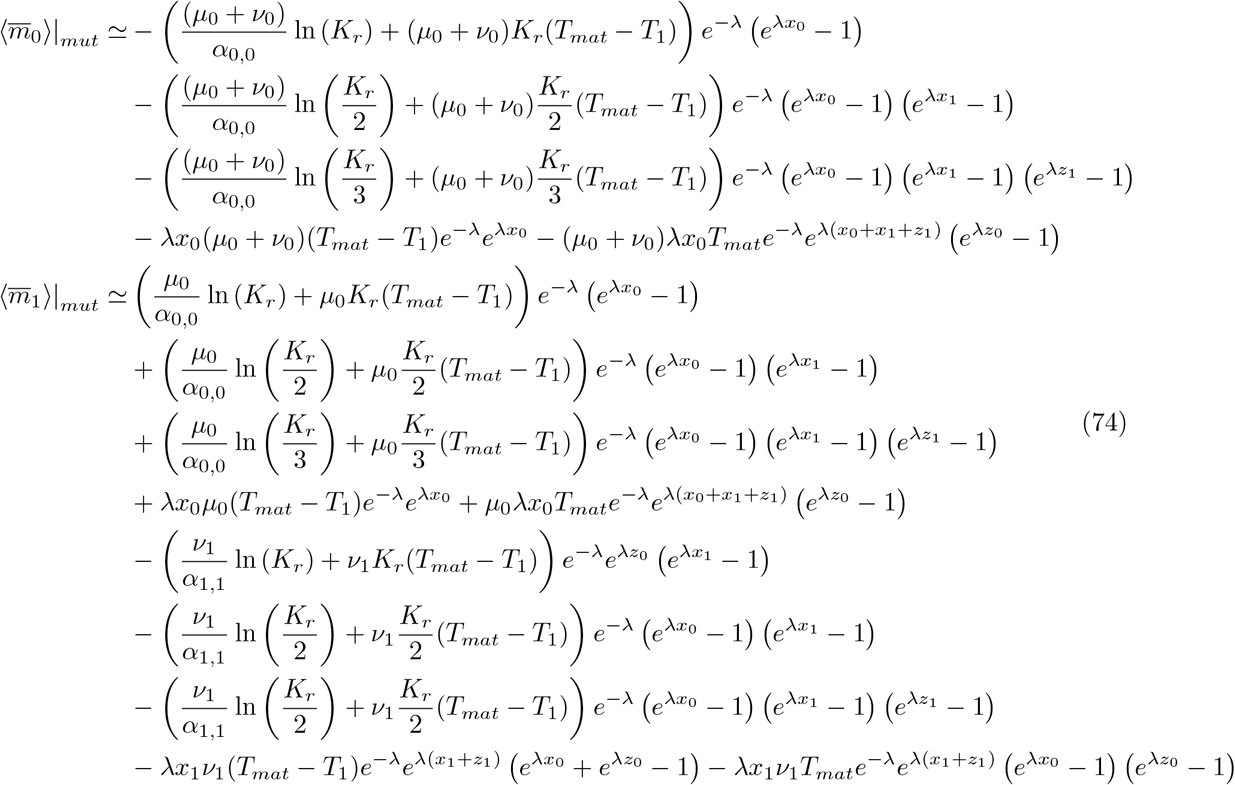

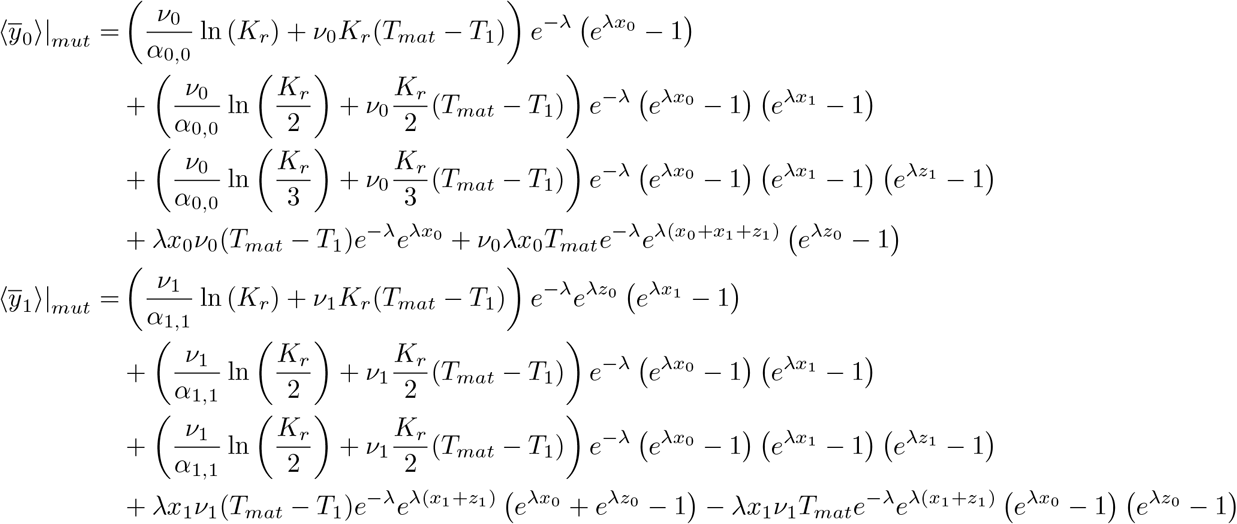

With such updated fractions, we can plot coexistence diagrams as done in Fig. 28. We still observe a complete extinction of the wild-type populations, except at large dilution factors (same as for aggressive generalist parasites). The diagrams also become mostly independent of the maturation time as the dynamics are not influenced by mutations anymore (WT populations being extinct), except from the flux from mutant replicases to mutant parasites. From this we deduce that generalist and non-aggressive parasites have a stronger fitness than specific and aggressive parasites in transient compartmentalization dynamics. This is also resulting from the fact that we assume that mutant replicases have a higher replication rate than wild-type replicases, and are therefore favored in the dynamics. Having a large population of mutant replicases leads to favoring mutant parasites.

### Trade-off Generality/Replicability with slow mutants (model B)

If we assume that parasites mutate to acquire generality, at the cost of replicability, but that mutant replicases have a similar growth rate as wild-type replicases. Therefore, mutant replicases will not take over wild-type parasites, and thus wild-type parasites can remain in the total population. We obtain regimes where all species co-exist at small dilution factors, with parasites and replicases having comparable fractions. There is a transition depending on the dilution factor from a large fraction of replicases and a dependency on maturation time, to a higher fraction of parasites independent of the maturation time. The rules here are: (1) In compartment with wild-type replicases and wild-type parasites, wild-type parasites take over the compartment regardless of the rest of the composition. (2) In compartments with wild-type replicases and mutant replicases only, they both grow up the half of the carrying capacity. (3) In compartments with wild-type replicases or mutant replicases and mutant parasites, but no wild-type replicases, all the species grow until they reach the same proportions of the carrying capacity. (4) In compartments with no replicases, nothing can grow. In this case we obtain for the updated fractions:

**Figure S28.**
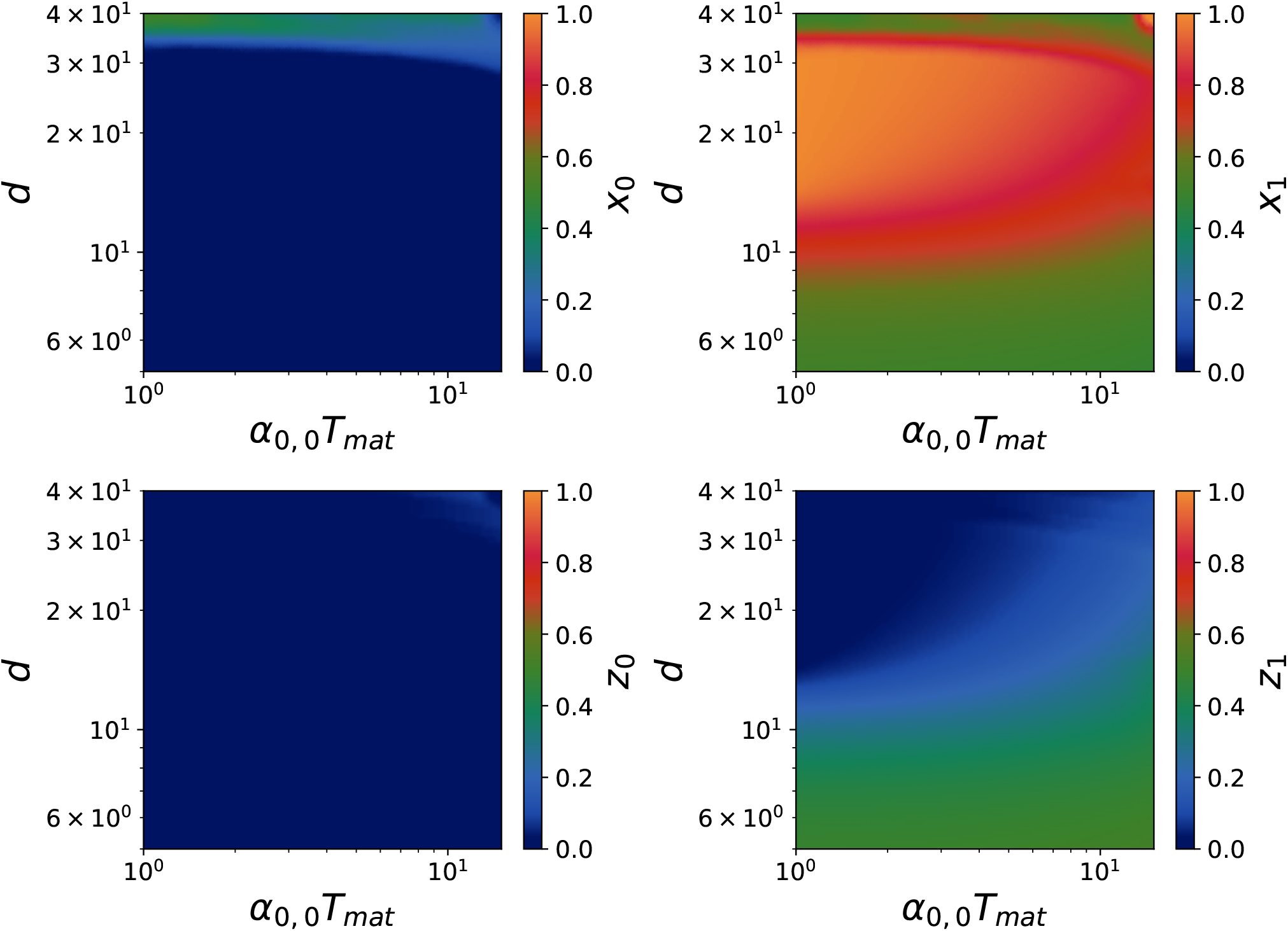
Coexistence diagram of the fractions of each species in the system with a Generality/Replicability trade-off in model B (theoretical expectations) after 300 cycles. We used for replication rates *α*_0,0_ = 5 × 10^−2^ *h*^−1^, *α*_1,1_ = 5 × 10^−2^ *h*^−1^, *γ*_0,0_ = 1. *h*^−1^, *γ*_0,1_ = 1 × 10^−1^ *h*^−1^, *γ*_1,1_ = 2.5 × 10^−1^ *h*^−1^, and for mutation rates *µ*_0_ = 5 1× 0^−4^ *h*^−1^, *ν*_0_ = 5 × 10^−4^ *h*^−1^. The carrying capacities are *K*_*p*_ = *K*_*r*_ = 30 and dilution factor *d* = 10. The panels represent the fractions for *x*_0_, *x*_1_, *z*_0_, *z*_1_ from right to left and from top to bottom.

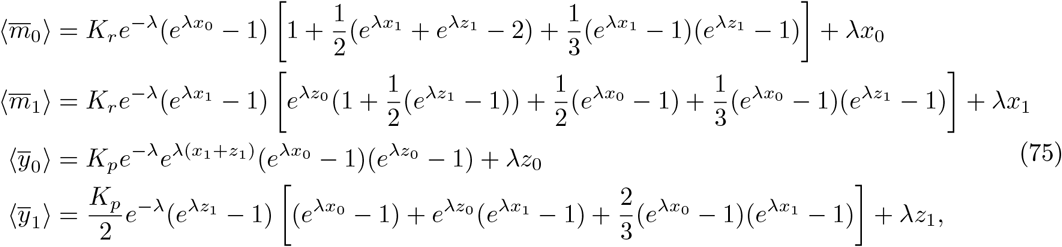

And for mutant numbers, we find that

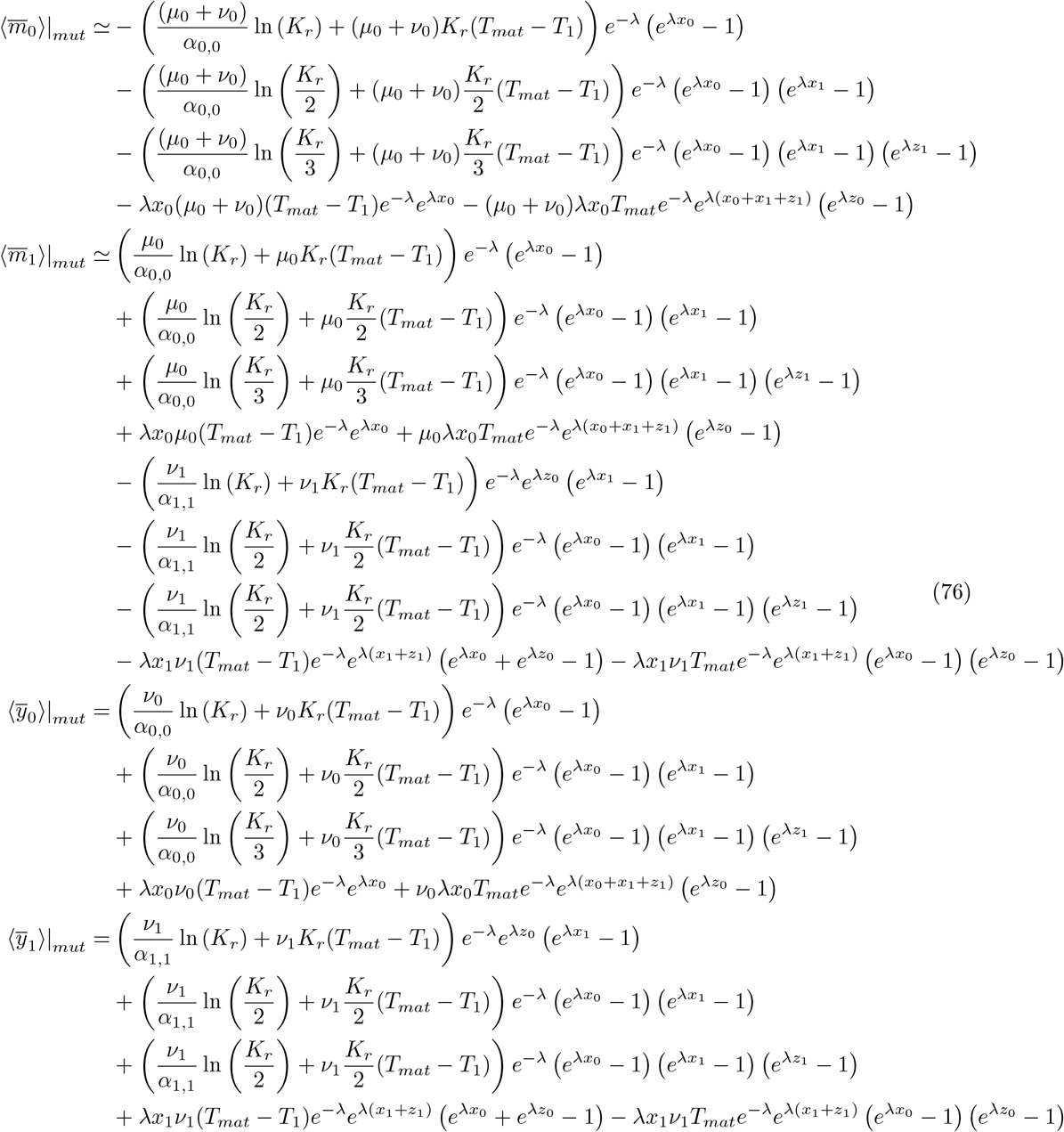

This model is in agreement with experimental observations where specialist and generalist parasites co-exist. The coexistence diagrams depending on the dilution factor *d* and maturation time *T* are given in 29, for the fully deterministic case (*s* = 1). Interestingly we also observe a phase where mutant replicases dominate at intermediate dilution factors and small maturation times. This corresponds to a phase where dilution is not strong enough to wash out wild-type parasites, which are aggressive on wild-type replicases.

### Influence of the carrying capacity

Up to this part, the carrying capacity *K*_*p*_ = *K*_*r*_ was fixed (*K* = 30). Increasing the carrying capacity typically favors parasites.

#### Coexistence diagram (*d, K*) (model A)

We can plot coexistence diagrams with a fixed maturation time *T* and varying the dilution factor *d* and carrying capacity *K*. Phase separations should be linear in a coexistence diagram (*d, K*) for transient compartmentalization dynamics. We find Fig. 30.

We observe that the phase separations are linear. We also have that increasing the carrying capacity *K* favors parasites. We can also plot state diagrams, and observe that at large dilution factors and carrying capacities, the dynamics are oscillating (unstable spirals). Decreasing the carrying capacity at a fixed dilution factor first lead to a phase with damped oscillations and to a phase with extinctions of some populations.

#### coexistence diagram (*d, K*) (model B)

In model B, the results are similar, as you can see in Fig. 31. The main difference is that replicases never completely take over the dynamics due to mutations from replicases to parasites.

Stable solutions are found by investigating steady states. Typically, in the large *K* limit, *K*_*p*_ = *K*_*r*_ :

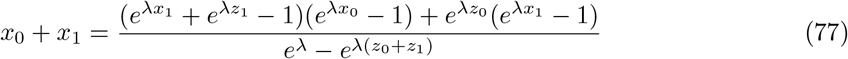

This equation setting the steady fraction of replicases in the system has several implications. In the limit of large *λ*, this yields *x*_0_ + *x*_1_ = 0. In the limit *λ* → 0, we obtain *x*_0_ + *x*_1_ = 1. In addition, *λ* is set by the equation:

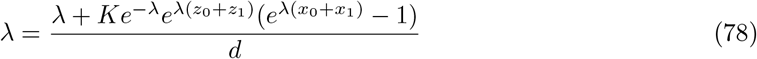

And thus, for *x*_0_ + *x*_1_ *>* 0, *λ* ∼ *K/d*. Thus we recover the expectations from the diagram Fig. 31 where *x*_0_ + *x*_1_ is small in the region where *K* ≫ *d*, and *x*_0_ + *x*_1_ → 1 in the region *d* ∼ *K*.

### From single compartments to total population

From our simulations we can compare the trajectories of single compartments to the average in the whole population. In practice we observe a strong variation from the trajectory of one compartment to another. We illustrate this strong variability in Fig. 32, where we plot the average fraction of each species, along with the content of single compartments (purple dots). This is done for model B, using simulations with stirring where each compartment follows a deterministic evolution but the stirring step in stochastic. At the scale of one compartment, the population is usually either close to 0 or close to the carrying capacity *K*. The average is rather depending on the number of full compartments (which can be estimated from the shadings of the dots) compared to the number of empty compartments, than on the populations per compartments.

**Figure S29.**
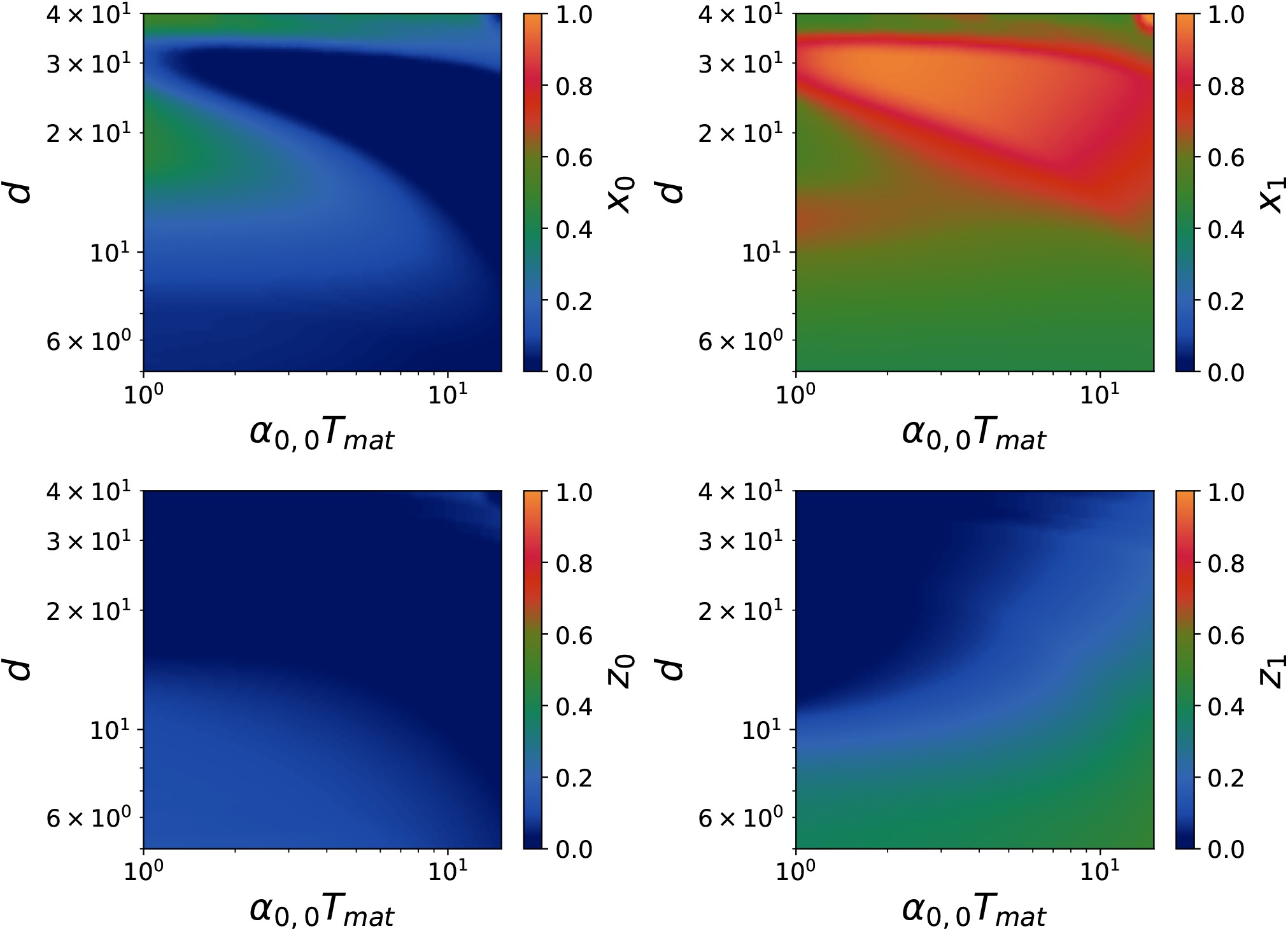
Trade-off Generality/Replicability (slow mutants) with the deterministic approach for *s* = 1. We used for replication rates *α*_0,0_ = 5 × 10^−1^ *h*^−1^, *α*_1,1_ = 5 × 10^−1^ *h*^−1^, *γ*_0,0_ = 10. *h*^−1^, *γ*_0,1_ = 5 × 10^−1^ *h*^−1^, *γ*_1,1_ = 5 × 10^−1^ *h*^−1^, and for mutation rates *µ*_0_ = 7 × 10^−3^ *h*^−1^, *ν*_0_ = 7 × 10^−3^ *h*^−1^. The carrying capacities are *K*_*p*_ = *K*_*r*_ = 30 and dilution factor *d* = 10.

**Figure S30.**
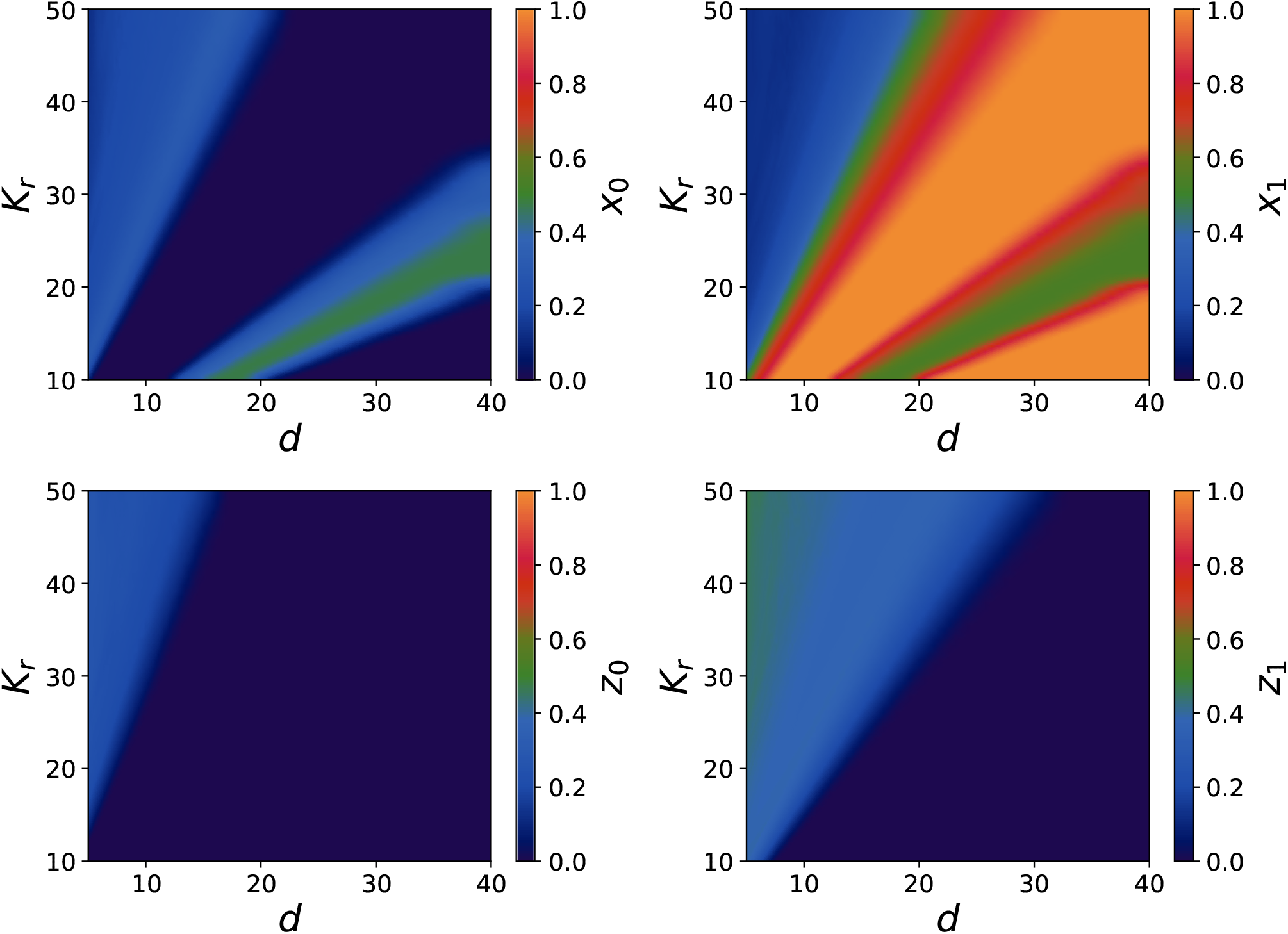
Coexistence diagram for the fractions in model A with stirring after 100 cycles with 8000 compartments, where *α*_0,0_ = 5 × 10^−1^ *h*^−1^, *α*_1,1_ = 2 *h*^−1^, *γ*_0,0_ = 7 *h*^−1^, *γ*_1,1_ = 20 *h*^−1^, and *µ*_0_, *ν*_0_, *ν*_1_ = 7 × 10^−3^ *h*^−1^. The carrying capacities dilution factor is *d* = 10. The panels represent the fractions for *x*_0_, *x*_1_, *z*_0_, *z*_1_ from right to left and from top to bottom.

**Figure S31.**
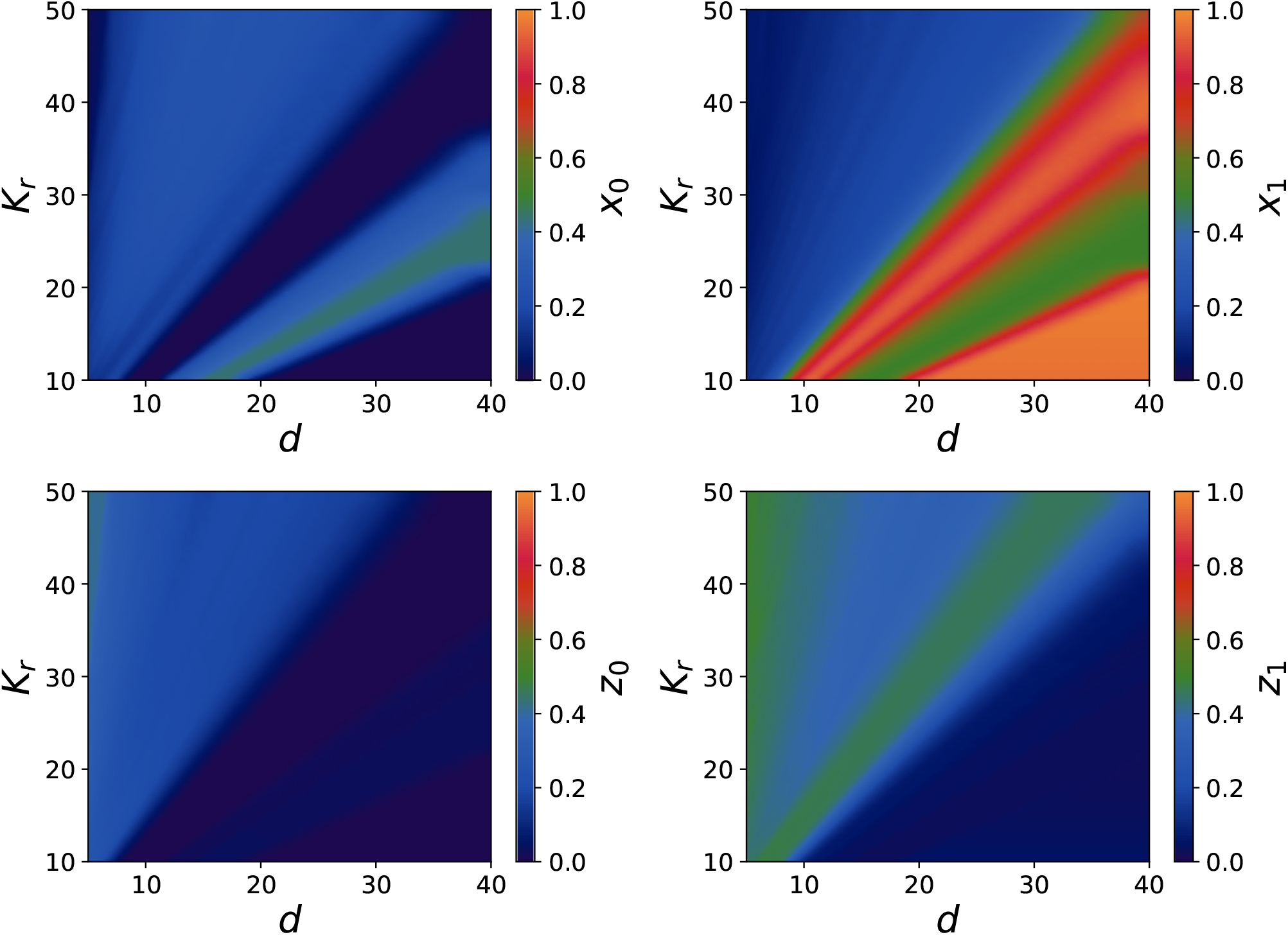
Coexistence diagram for the fractions in model B with stirring after 100 cycles with 8000 compartments, where *α*_0,0_ = 5 × 10^−2^ *h*^−1^, *α*_1,1_ = 3 × 10^−1^ *h*^−1^, *γ*_0,0_ = 1 *h*^−1^, *γ*_1,1_ = 2.5 *h*^−1^, and *µ*_0_, *ν*_0_, *ν*_1_ = 1 × 10^−3^ *h*^−1^. The carrying capacities are *K*_*p*_ = *K*_*r*_ = 30 and dilution factor *d* = 10. The panels represent the fractions for *x*_0_, *x*_1_, *z*_0_, *z*_1_ from right to left and from top to bottom.

**Figure S32.**
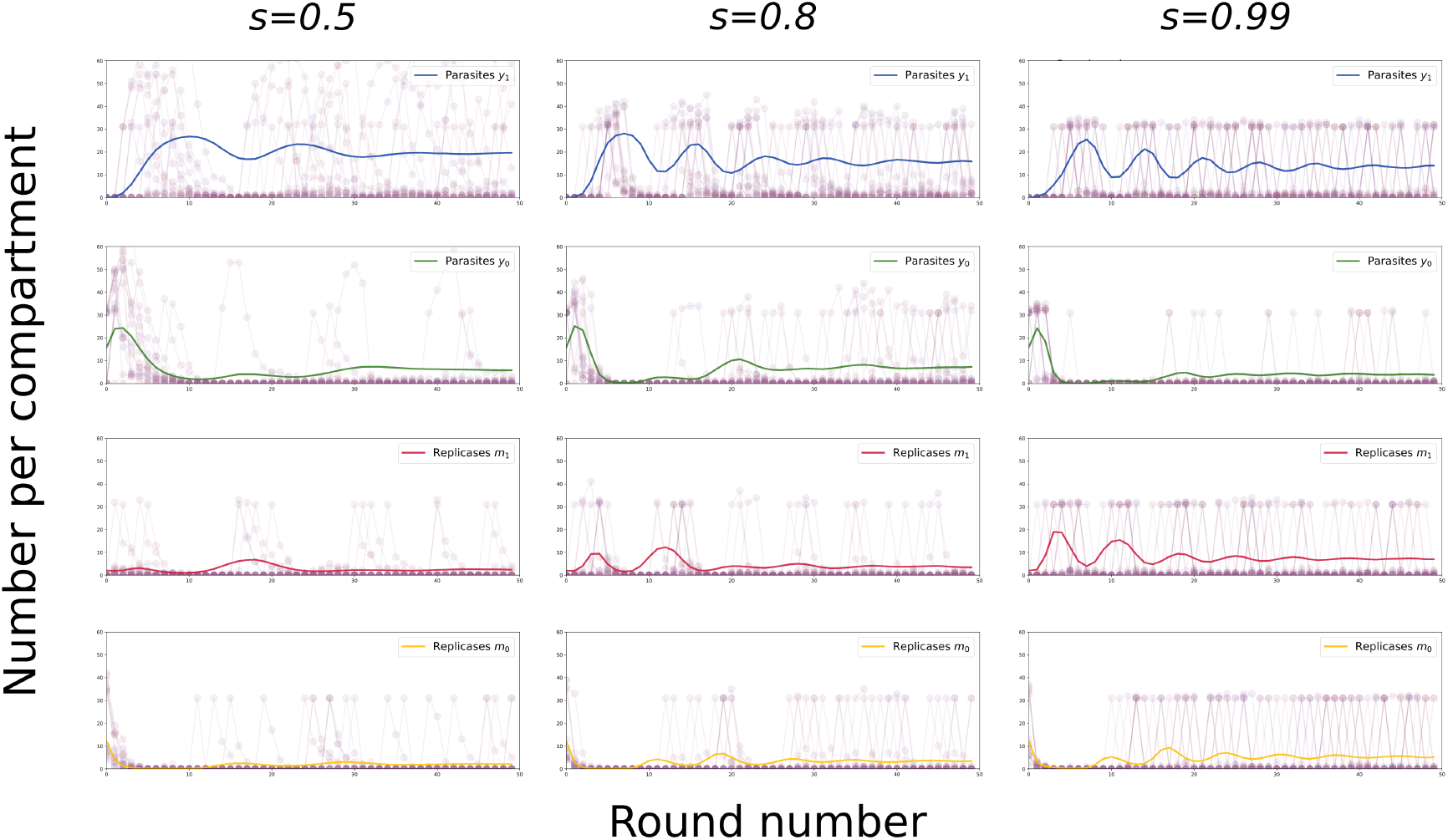
Volatility of the trajectories of single compartments (purple dots), and average fractions (colored lines), with *s* = 1. (model B). The columns represent different values of the stirring parameter *s*, and rows correspond to different species *x*_0_, *x*_1_, *y*_0_, *y*_1_ from bottom to top.

We observe that for weak stirring, the single compartments are more likely to have intermediary compositions where two species share the compartment instead of having one reaching the carrying capacity and the rest close to 1.

### Length dependence of the carrying capacity

Up to this section we assumed that *K*_*p*_ = *K*_*r*_, however, the lengths of parasites is smaller than that of replicases, so that the carrying capacity of parasites should be larger than that of replicases. In particular we can assume that the lengths of parasites is related to that of replicases through *L*_*r*_ ∼ 4*L*_*p*_. Indeed HL1-, HL2-, PL2-, and PL3-228 are respectively 2041, 2042, 514, and 514 nt long. If we assume that the number of resources required for one nucleotide is fixed, we have *K*_*p*_ ∼ 4*K*_*r*_.

**Figure S33.**
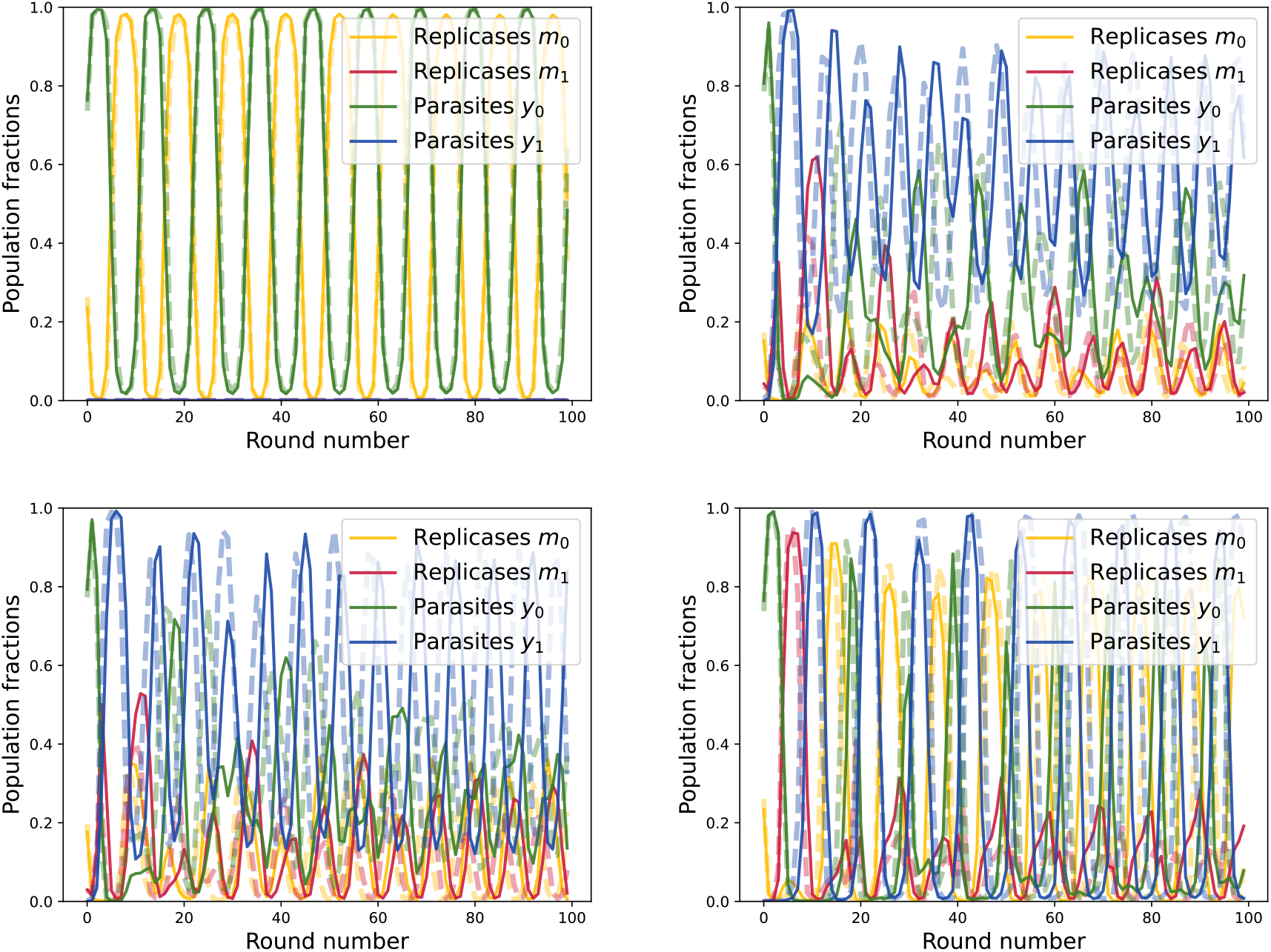
Dynamics with different carrying capacities for parasites and replicases within model B. We used for replication rates *α*_0,0_ = 5 × 10^−1^ *h*^−1^, *α*_1,1_ = 2 *h*^−1^, *γ*_0,0_ = 7 *h*^−1^, *γ*_1,1_ = 20 *h*^−1^. **Upper left** *µ* = *ν* = 0 *h*^−1^. **Upper right** *µ* = *ν* = 10^−2^ *h*^−1^. **Lower left** *µ* = *ν* = 7.10^−3^ *h*^−1^. **Lower right** *µ* = *ν* = 2.10^4^ *h*^−1^.

In this case, the results are modified, as parasites populate the compartments faster. In particular, oscillations are stronger and parasites periodically populate the system entirely or are alternatively completely removed. We obtain the dynamics shown in Fig. 33. We also observe that the theoretical deterministic approach captures the behaviour of the system in average, and in the oscillations amplitudes, but the finite size of Gillespie simulations generate a discrepancy between the periods of oscillations between Gillespie and the deterministic approach. The different states of the system are comparable to that for *K*_*p*_ = *K*_*r*_: at small dilution factors, mutant parasites dominate, at large dilution factors replicases dominate. Nevertheless, the domain of dominance of mutant replicases is smaller in this case and mutant parasites survive on a larger domain. in Fig. 35 mutant replicases are also present on the whole diagram.

**Figure S34.**
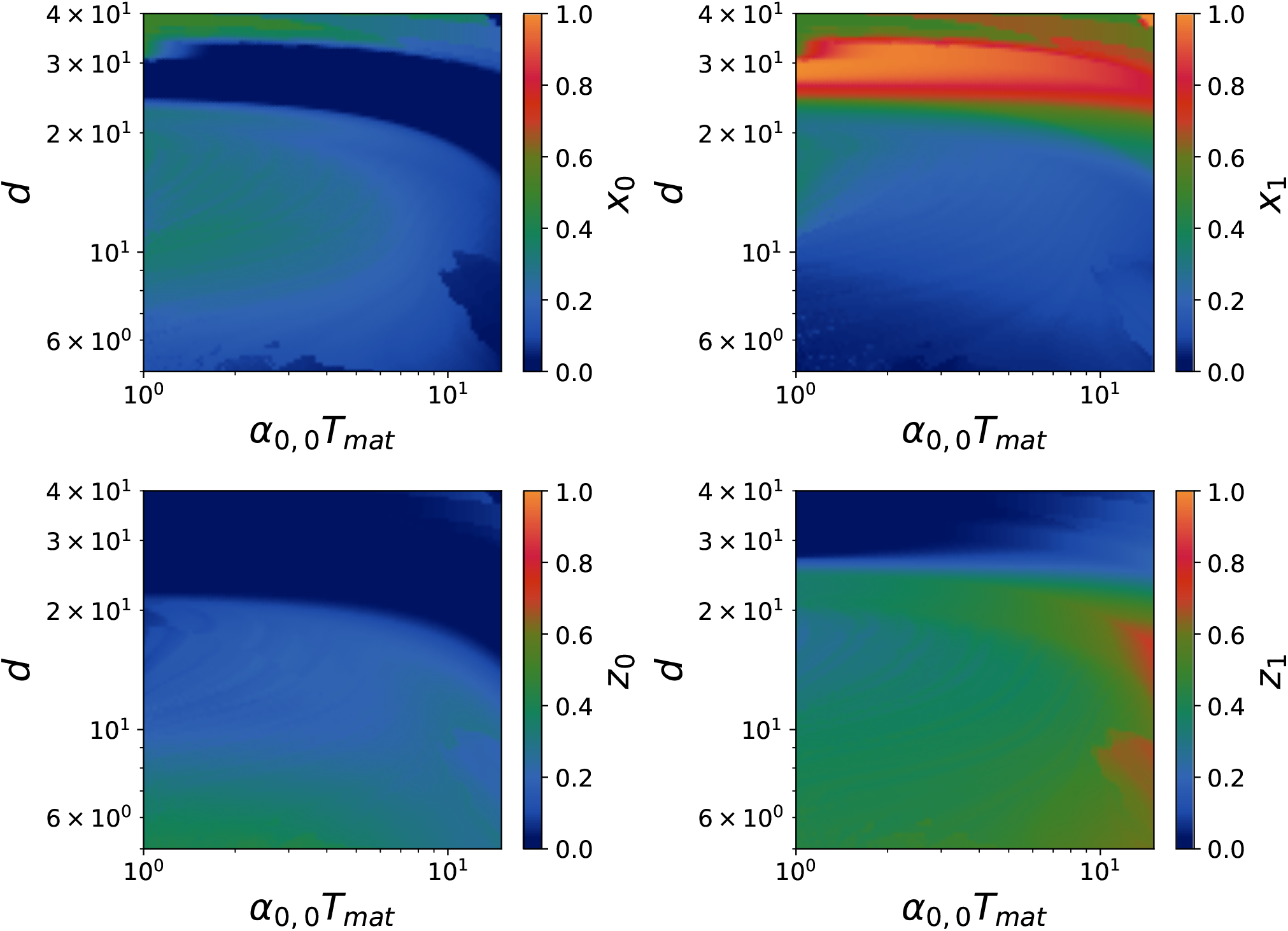
Coexistence diagram of the states of the system in model B where hosts mutate to parasites after 300 cycles. We used for replication rates *α*_0,0_ = 5 × 10^−2^ *h*^−1^, *α*_1,1_ = 5 × 10^−2^ *h*^−1^, *γ*_0,0_ = 1. *h*^−1^, *γ*_0,1_ = 1 × 10^−1^ *h*^−1^, *γ*_1,1_ = 2.5 × 10^−1^ *h*^−1^, and for mutation rates *µ*_0_ = 5 × 10^−4^ *h*^−1^, *ν*_0_ = 5 × 10^−4^ *h*^−1^. The carrying capacities are *K*_*p*_ = 4*K*_*r*_ = 120 and dilution factor *d* = 10. The panels represent the fractions for *x*_0_, *x*_1_, *z*_0_, *z*_1_ from right to left and from top to bottom.

**Figure S35.**
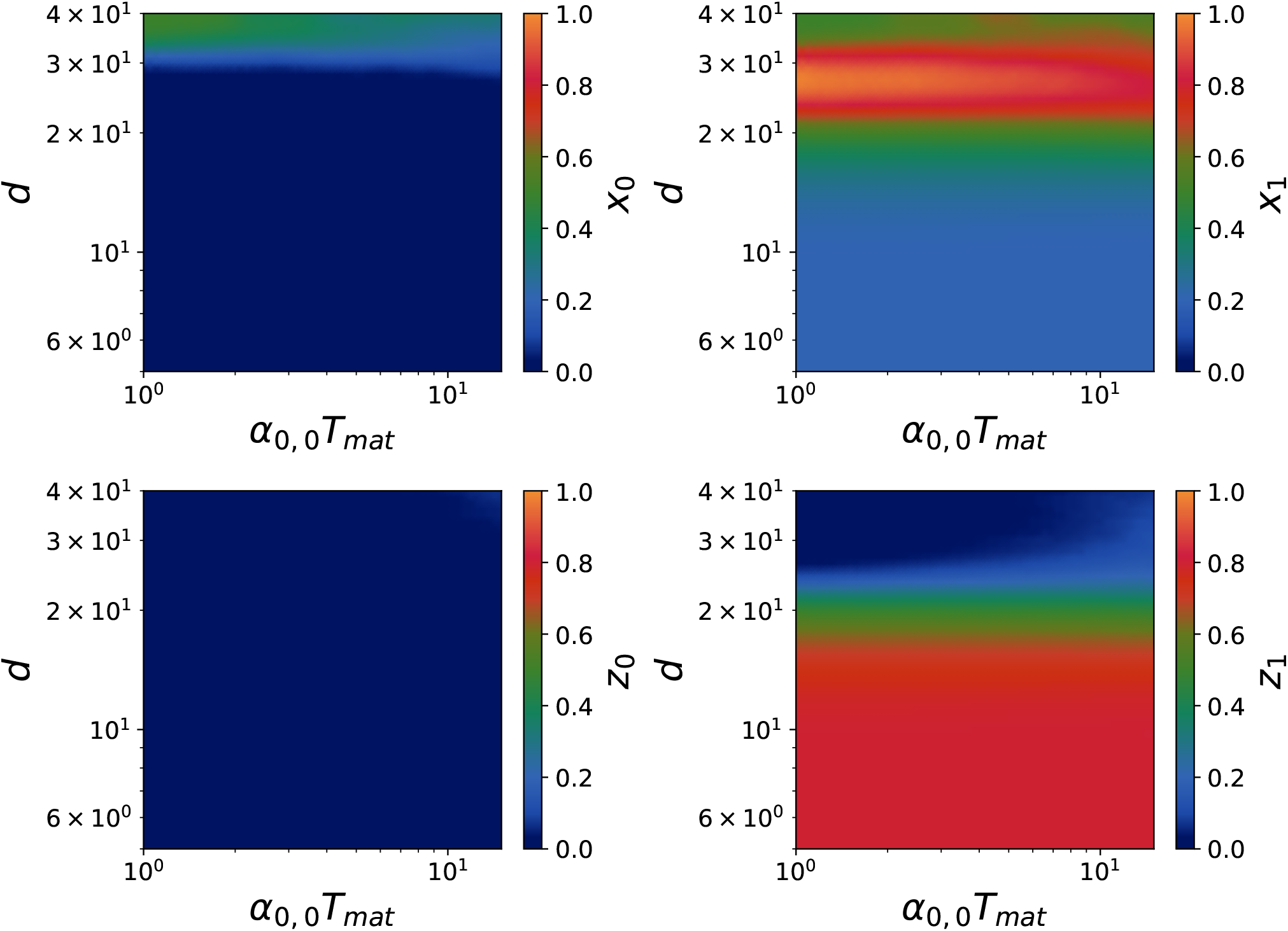
Coexistence diagram of the states of the system in model B in a Generality/Replicability trade-off after 300 cycles. We used for replication rates *α*_0,0_ = 5 × 10^−2^ *h*^−1^, *α*_1,1_ = 5 × 10^−2^ *h*^−1^, *γ*_0,0_ = 1. *h*^−1^, *γ*_0,1_ = 1 × 10^−1^ *h*^−1^, *γ*_1,1_ = 2.5 × 10^−1^ *h*^−1^, and for mutation rates *µ*_0_ = 5 × 10^−4^ *h*^−1^, *ν*_0_ = 5 × 10^−4^ *h*^−1^. The carrying capacities are *K*_*p*_ = 4*K*_*r*_ = 120 and dilution factor *d* = 10. The panels represent the fractions for *x*_0_, *x*_1_, *z*_0_, *z*_1_ from right to left and from top to bottom.

**Figure S36.**
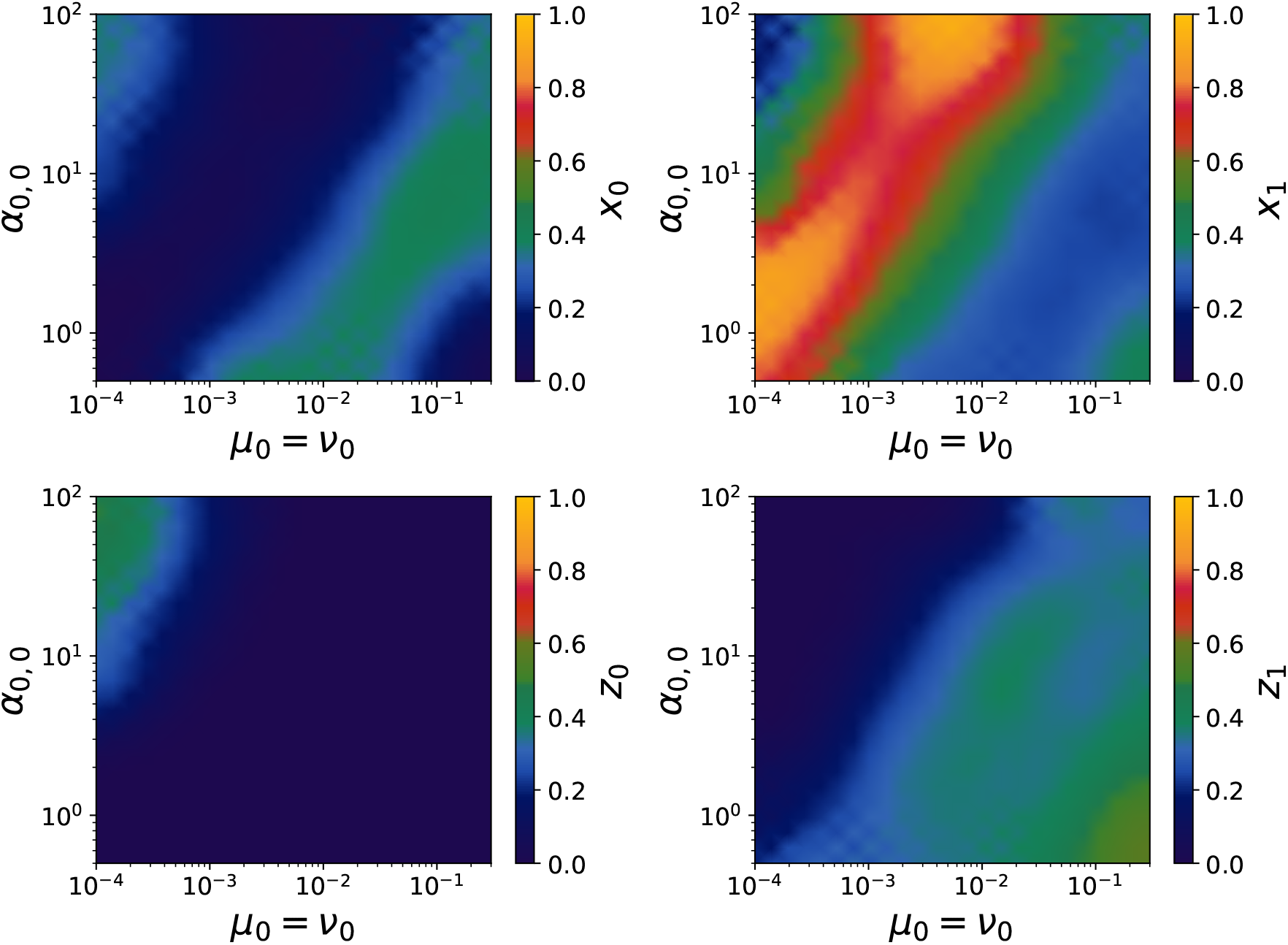
Coexistence diagram (*µ, α*) for model A. We study the state of the system (*x*_0_, *x*_1_, *y*_0_, *y*_1_) as a function of replication rates and mutation rate. Transitions between different states are linear.

### Boundaries in coexistence diagram (*µ, α*)

We can observe the influence of the difference between replication and mutation, by fixing the ratio between the replication rate of wild-type replicases and mutant replicases, the ratio between the replication rate of wild-type replicases and wild-type parasites and the ratio between the replication rate of wild-type replicases and mutant parasites. Replication and mutation rates will indirectly depend on the lengths of RNA strands. Those coexistence diagrams are realized using Gillespie simulations as it reaches phases where the deterministic model fails to describe the dynamics.

#### Model A

For model A, we observe linear phase transitions. This suggests that for fixed ratio *α*_1,1_*/α*_0,0_, *γ*_0,0_*/α*_0,0_ and *γ*_1,1_*/α*_0,0_, the value of the ratio *µ*_0_*/α*_0,0_ fixes the final repartition of the population.

This coexistence diagram results from the expression of the number of mutants appearing at each round. In particular, we that the number of mutants depends on the ratios *µ*_0_*/α*_0,0_ and *ν*_0_*/α*_0,0_ (as the maturation time is set to 10*/α*_0,0_ in the simulations by default). This explains the linear behaviour observed.

#### Model B

In model B, we also observe linear phase transitions. In this case, wild-type species mostly go extinct.

**Figure S37.**
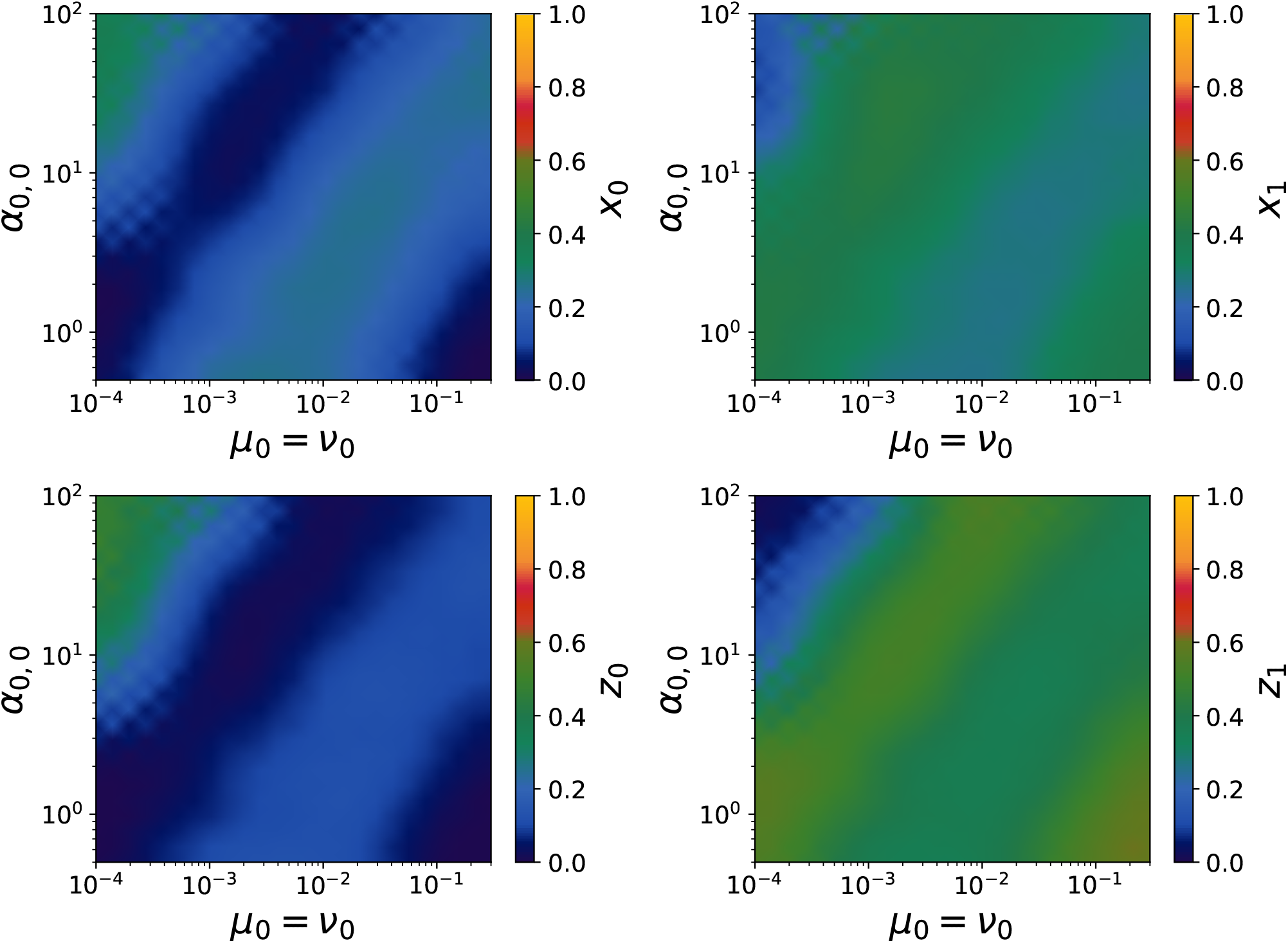
Coexistence diagram (*µ, α*) for model B. We study the state of the system (*x*_0_, *x*_1_, *y*_0_, *y*_1_) as a function of replication rates and mutation rate. Transitions between different states are linear.

## Notes

### Competing Interest Statement

The authors have declared no competing interest.

### Summary of Updates

I changed the aspect ratio in Figure 4 of the main text, plus added minor changes to the text.

